# The genetics of immune and infection phenotypes in wild mice, *Mus musculus domesticus*

**DOI:** 10.1101/2022.07.26.501599

**Authors:** Louise Cheynel, Luke Lazarou, Eleanor M. Riley, Mark Viney

**Affiliations:** Department of Evolution, Ecology and Behaviour, University of Liverpool, Liverpool, L69 7ZB, UK; School of Biological Sciences, University of Bristol, Bristol, BS8 1TQ, UK; Institute of Immunology and Infection Research, School of Biological Sciences, University of Edinburgh, Edinburgh, EH9 3FL, UK

## Abstract

Wild animals are under constant threat from a wide range of micro- and macroparasites in their environment. Animals make immune responses against parasites, and these are important in affecting the dynamics of parasite populations. Individual animals vary in their anti-parasite immune responses. Genetic polymorphism of immune-related loci contributes to inter-individual differences in immune responses, but most of what we know in this regard comes from studies of humans or laboratory animals; there are very few such studies of wild animals naturally infected with parasites. Here we have investigated the effect of polymorphism in immune-related loci (the MHC, and genes coding for cytokines and Toll-like receptors) on a wide range of immune and infection phenotypes in UK wild house mice, *Mus musculus domesticus*. We found strong effects of polymorphisms in various MHC and cytokine coding loci on both immune measures (antibody concentration and cytokine production) and on infection phenotypes (infection with mites, worms and viruses). Our study provides a comprehensive view of how polymorphism of immune-related loci affects immune and infection phenotypes in naturally infected wild rodent populations.

## 1 Introduction

Wild animals are under constant threat from a wide range of micro- and macroparasites present in their environment. Animals use a variety of behavioural and physiological responses to avoid and / or resist infection and the harm that it can cause. A key aspect of this are immune responses that actively defend individuals against infection. However, parasites also evolve to avoid these immune responses. The ubiquity of parasites and their continual evolution represents a strong selective force that hones the immune responses that animals make (Moran, 2002; Pilosof et al., 2014). This ongoing co-evolution between hosts and parasites shapes both parasites’ genetic diversity and the genetic diversity of host loci whose products mediate the immune response, henceforth immune-related loci (Anderson & May, 1982; Frank, 2000).

In vertebrates, some immune-related loci, such as those in the major histocompatibility complex (MHC), are highly polymorphic (Radwan et al., 2020) and this genetic diversity is thought to be the result of the parasite-driven selection (Spurgin & Richardson, 2010). The MHC encodes proteins that present antigens to T lymphocytes, which can then result in an immune response (Klein, 1986). MHC heterozygosity is thought to be advantageous by facilitating the presentation of a wide variety of antigens, so widening the range of potential immune responses that can be generated; however, experimental work to test this hypothesis has provided mixed results (Doherty & Zinkernagel, 1975; Penn et al., 2002; Ilmonen et al., 2007). Moreover, there is a much wider repertoire of immune-related loci that may also affect individuals’ immune responses (Acevedo-Whitehouse & Cunningham, 2006). Animals use Toll-like receptors (TLRs) to detect bacterial and viral pathogen associated molecular patterns (PAMPs), and TLRs then initiate signalling events, including via cytokines and chemokines, that ultimately activate innate and adaptive immune responses, so contributing to parasite resistance (Kawai & Akira, 2006). Most mammals have 10 – 12 TLR coding loci that detect different pathogen-associated moieties (Roach et al., 2005). In humans, numerous studies have described associations between polymorphisms in TLR and cytokine coding loci, and susceptibility or resistance to a range of infections (Smith & Humphries, 2009; Mukherjee et al., 2019). In humans, the cumulative contribution of non-MHC loci, such as those coding for cytokines or PAMP receptors, to the immune phenotype likely exceeds that of the MHC (Jepson et al., 1997; Roederer et al., 2015).

Most of our understanding of the effect of polymorphism in immune-related loci on immune and / or infection phenotypes comes from studies of humans or laboratory animals; few empirical studies have been conducted with wild animals. Major differences exist between the infection and immune states of wild animals and their laboratory-reared counterparts, with this extensively elaborated for mice, including showing the significantly different immunological effect of wild vs. lab gut microbiomes (Beura et al., 2016; Abolins et al., 2017; Rosshart et al., 2017). Outbred lines of laboratory mice are available, though these do not reflect the full genetic diversity of wild populations (Turner & Paterson, 2013). Given the disconnect between the immunological state, infection state, and genetic diversity of wild and lab animals, the paucity of our understanding of how genetic effects affect immune responses and infection phenotypes in wild animals is a notable knowledge gap, and points to the critical need to study these phenomena directly (Turner & Paterson, 2013).

To date there have been about thirty immunogenetic studies on 16 different genera of wild mammals (Table 1). Among these the effect of the MHC has been investigated most frequently and these studies commonly, but not universally, find effects of MHC diversity on micro- and macroparasites infection phenotypes. Studies of polymorphisms in TLR and cytokine coding loci have also found associations with infection phenotypes. Most of these studies have studied the effects of polymorphism in immune-related loci on infection phenotypes (especially of macroparasite burden), and so there remains major gaps in our understanding of effects on immune responses themselves (but see Charbonnel et al., 2010; Cutrera et al., 2011; Turner et al., 2011; Huang et al., 2021). Overall, polymorphism in rather few immune-related loci have been studied in wild mammals, again in notable contrast to the very large number of loci studied in laboratory animals. It is not yet known whether these laboratory observed effects also occur in the wild. An important remaining question is therefore whether – and if so, how – naturally occurring polymorphism in immune-related loci impacts both immune and infection phenotypes in wild mammals.

**Table 1.**
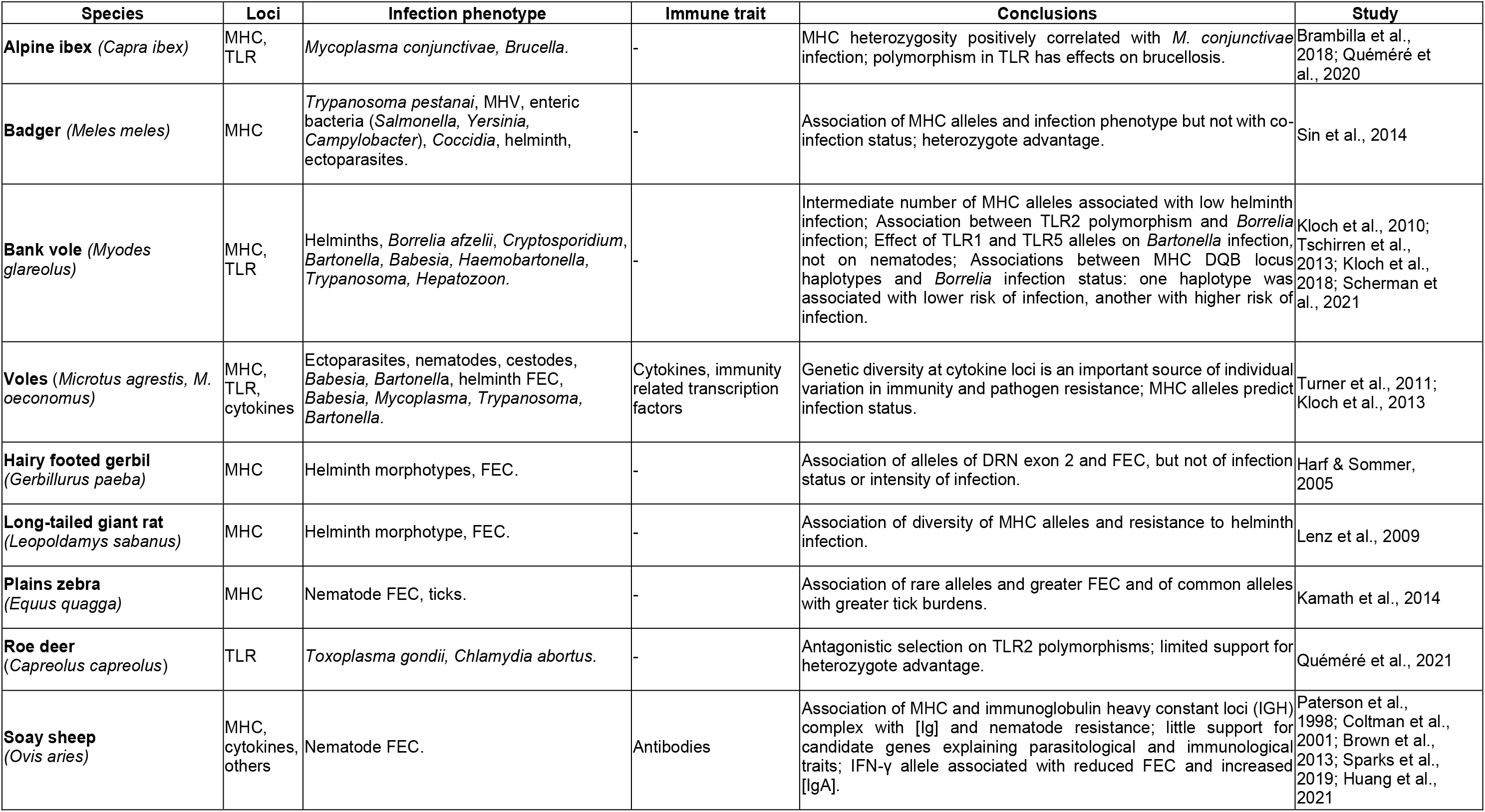
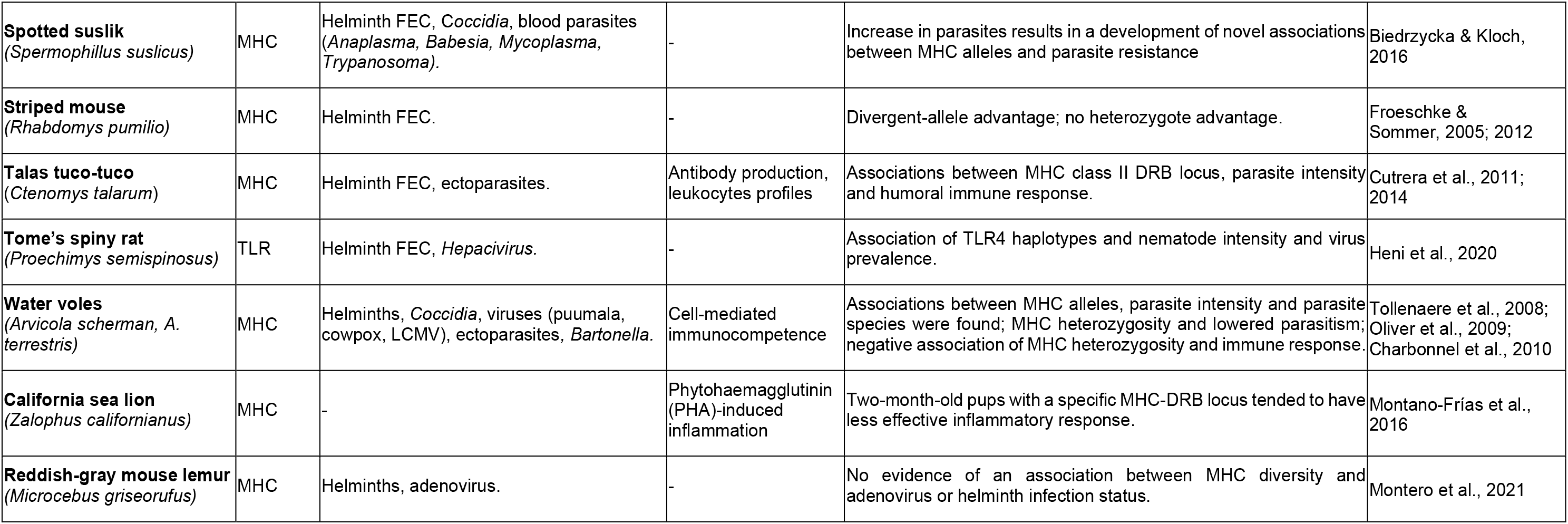
Summary of studies in wild mammals testing associations between genetic variation in immune-related genes and immune or infection phenotypes. MHC, Major Histocompatibility Complex; TLR, Toll Like Receptor; MHV, murine hepatitis virus; FEC, faecal egg counts; GI, gastrointestinal; LCMV, Lymphocytic choriomeningitis virus. These articles were searched for from the Web of Science using combinations of key words “immuno-genetic”, “genetic variation”, “immunity”, “infection” and “wild”.

Here we have investigated the effect of diversity in immune-related loci on a wide range of immune and infection phenotypes in UK wild house mice, *Mus musculus domesticus*. We first assessed the extent of genetic variation in 23 immune-related loci (coding for cytokines, TLRs or the MHC) in our study populations, finding variation in 14 of these loci among 435 mice. We then tested the effect of polymorphism in these 14 loci on 24 discrete immune phenotypes, encompassing both cellular and humoral components of the innate and the adaptive immune response. We also tested the effect of their polymorphism on infection phenotypes of 9 different micro- and macroparasites (mites, worms, viruses and a bacterium). To look for effects of the genotype on the phenotype we used a two-step model selection approach, firstly testing for the effect of all non-genetic factors (site, sex, age and body mass), then secondly adding each genetic factor to the model in turn to assess its effect on the relevant phenotype. Our study thus provides a unique and more complete picture linking polymorphism in 14 immune-related loci to 24 immune phenotypes and 9 infection phenotypes in wild rodents.

## 2 Methods

### 2.1 Mice

The sampling, processing, determination of infection state, and immune phenotyping of the mice has previously been reported (Abolins et al., 2017 and 2018). In brief, 460 mice (*Mus musculus domesticus*) were live trapped from 12 sites in the southern UK between March 2012 and April 2014. After being humanely killed, mice were sexed, weighed and measured from the tip of the snout to the base of the tail. Female mice that were later found to be pregnant had the mass of the foetuses subtracted from their final mass, and these values were used in all subsequent analyses. To estimate body condition, the Scaled Mass Index (SMI) was calculated as previously described (Peig & Green, 2009). Age was determined using eye lens mass as previously described (Rowe et al., 1985). The distribution of individuals by study site, sex, and age class is given in Supporting Information 1.

### 2.2 Immune measures

The immune state of the mice was assessed by measuring (i) 12 immune cell populations, (ii) the concentration of 3 immunoglobulins, and (iii) the concentration of 9 cytokines produced after *in vitro* stimulation of spleen cells with 4 different stimuli (anti-CD3/anti-CD28, CpG, LPS or PG), which together encompass cellular and humoral components of the innate and adaptive immune responses (as reported in Abolins et al., 2017 and 2018).

The 12 cell populations measured were flow cytometric counts of splenic NKp46+ Natural Killer (NK) cells; CD19+ B cells; CD11c+ Dendritic cells (DCs); CD8+ T cells; CD4+ T cells; CD25+ FoxP3+ Treg cells; F4/80+ Ly6G-macrophages; F4/80+ Ly6G-low monocytes; F4/80+ Ly6G-intermediate hyper-granulocytic myeloid cells; F4/80 variable Ly6G-High polymorphonuclear (PMN) cells; FSC-low PMNs neutrophils; and FSC-High PMNs Myeloid-Derived Suppressor Cells (MDSC) (as Abolins et al., 2017). Each of the 12 cell counts was then scaled to the mouse body mass, using the same formula as the one used for the calculation of the SMI (see above), but replacing mouse mass with the count for each cell type. The relevant scaling component was calculated separately for each of the 12 cell types. This results in the count of each immune cell for each mouse scaled to the length of the average mouse. Scaling the immune cells in this manner, rather than using a proportion of each cell type, allowed us to use count data in our modelling while still controlling for effects of mouse body size.

The immunoglobulins measured were the serum concentrations of IgG and IgE and the faecal concentration of IgA. The cytokines measured were IFN-γ, IL-1β, IL-4, IL-6, IL-10, IL-12p40, IL-12p70, IL-13, MIP-2α.

### 2.3 Infection phenotype

The number of fur mites, *Myocoptes musculinus*, was determined as previously described and classified into the categories: 0–10, 11–20, 21–30, 31–40, 41–50, 51–100, 101–200, 201–300, 301–400, 401–500, 501–1000, 1001–1500, 1501–2000 (Weldon *et al.*, 2015). The number of intestinal nematodes was determined by gut examination as previously described (Weldon et al., 2015). Current or prior infection with Corona, Mouse Hepatitis, Sendai, Minute, Noro and Parvo viruses and to *Mycoplasma pulmonis* was inferred from serological data, as previously described (Abolins et al., 2017).

### 2.4 SNP identification and genotyping

Our approach to identifying SNPs and then genotyping wild mice at these SNPs was as follows. First, we identified 23 loci coding for products of immunological interest, specifically 15 coding for cytokines (IL-1a, IL-1b, IL-2, IL-4, IL-6, IL-10, IL-2a, IL-2b, IL-13, IL-17a, IL-17F, IFN-γ, TNF, IL2-rg, CD40lg), 3 coding for TLRs (TLR4, 5 and 9), 5 for products of the murine major histocompatibility complex (MHC) H2 (H2-Aa, H2-Ab1, H2-Eb1, H2-K1, H2-D1). We also selected, as a control, a gene encoding a non-immunological product (Myo1a). Second, we PCR amplified fragments of these loci in a randomly selected sub-set of 25 wild mice from the different sites, and sequenced those amplicons to identify putative SNPs. Third, we selected 23 of these SNPs (in 21 immune-related loci: IL-1a, IL-1b [2 SNPs, IL-1b_U, IL-1b_N], IL-2, IL-4, IL-6, IL-10, IL-2a, IL-2b, IL-13, IL-17a [2 SNPs, IL-17a_U, IL-17a_N], IL-17F, IFN-γ, TNF, IL2-rg, CD40lg, TLR4, TLR5, TLR 9, H2-Aa, H2-Ab1, H2-Eb1) which we then KASP genotyped in our full set of wild mice (outsourced to LGC Genomics, UK). We found that 14 (IL-1a, IL-1b_U, IL-6, IL-10, IL-3, IL-17a_U, IL-17a_N, IL-17F, TNF, TLR5, TLR 9, H2-Aa, H2-Ab1, H2-Eb1) of these 23 SNPs were polymorphic in this full set of mice (Lazarou, 2019) and these are the data that are analysed here. The control locus Myo1a was also successfully amplified, sequenced and two SNPs (Myo1a_1, Myo1a_2) were found to be polymorphic in these mice. A summary table of the SNPs identified and variants found is provided in Supporting Information 2.

The PCR amplification of these fragments used the primer sequences shown in Supporting Information 3. The reactions consisted of 1 μL of genomic DNA, 4 μL (10 μM) of each primer, 5 μL (20 units) of Platinum Taq High fidelity Buffer (Life Technologies), 1 μL of dNTPs (10 mM each) (ThermoFisher Scientific), 2 μL of the manufacturer’s supplied MgSO4 (Life Technologies) and 32 μL of water (Sigma Aldrich), which were then cycled through the following conditions: denaturation at 94°C for 30s, annealing temperature 15s, elongation 68°C with time depending on product, calculated at a rate of 30s per kb, and a final elongation at 68°C for 10 minutes. Annealing temperatures were optimised using gradient PCR with C57/BL6 target DNA. The 25 wild mice used for SNP discovery were mouse numbers 17, 20, 37, 53, 64, 107, 112, 123, 137, 157, 244, 251, 265, 267, 282, 290, 309, 317, 327, 351, 354, 372, 379, 382 and 403, from Abolins et al. (2017) and Abolins et al. (2018) (SNPs identified are shown in Supporting Information 4 and variants detected are shown in Supporting Information 5). Successful PCR products were purified using the GeneJet PCR clean-up kit (Thermo Fisher Scientific), following the manufacturer’s instructions and single-read sequencing of the purified products was carried out by Eurofins Genomics, UK.

### 2.5 Statistical analyses

We tested all possible associations between individual SNPs and immune or infection phenotypes. We used generalized linear models (GLM) for continuous variables (immune measures and worm number) and ordinal logistic regression for categorial data (*i.e.* intensity of infection with mites and microparasites). Each immune or infection phenotype was used as the response variable in the model. Raw data were used when the immune parameter had a Gaussian distribution, while traits with a non-Gaussian distribution were log(X+1) transformed before analysis.

As our dataset comprised concentrations of 9 different cytokines produced *in vitro* after spleen cell stimulation with 4 different stimuli (*i.e.* 36 cytokine measures for each mouse), we performed a principal component analysis (PCA) on all mice for which all 36 cytokine measures were available (N = 172) to identify the main axes of variation, for which we used the R package “ade4” (Dray & Dufour 2007). We removed data for three mice that contributed abnormally to the PCs (*i.e.* contributing 8 % of the first principal component) possibly because of a severe inflammatory state. PC1 and PC2 represent 34% and 19% (total 53%) of the covariation among the cytokine measures. PC1 was mostly influenced by IL-1B, IL-10, IL-12p40, IL-12p70, IL-13, IL-4; PC2 was mostly influenced by IL-6, IFN-γ and MIP-2α (Supporting Information 6). For cytokine measures, we therefore used PC1 and PC2 as the response variable in the modelling approach below.

Our modelling approach had two steps. Firstly, for each immune or infection phenotype we built a base model to investigate the effect of potential non-genetic confounders, specifically site effect (nine sites, Figure 1), age (linear function or three classes, Supporting Information 1), sex (two classes: male and female) and body condition (SMI, continuous). We merged data from some closely located sites with small numbers of mice, giving us nine sites for analysis (Figure 1; Supporting Information 1). The three age classes were: 0-6 weeks (immature animals unlikely to be venturing far from their nest site); 7-12 weeks (young animals unlikely to have reproduced); >12 weeks (sexually mature adults). We tested the two-way interactions between site, age and sex. Secondly, we tested the effect of the SNPs in the immune-related genes. Depending on the variables retained in the base model, two-way interactions between SNP, site, sex, and age were also included. Using a model selection procedure based on the Akaike Information Criterion (AIC, Burnham & Anderson, 2002), we retained the model with the lowest AIC. When the difference in AIC between competing models was less than 2, we retained the model with the fewest parameters to satisfy parsimony rules (Burnham & Anderson, 2002). Residuals plots were checked to ensure the fit of the selected regression models. When a genetic term was selected in a model, the significance of its effect was additionally assessed by using deletion testing and the log-likelihood ratio test (McCullagh & Nelder, 1989).

**Figure 1.**
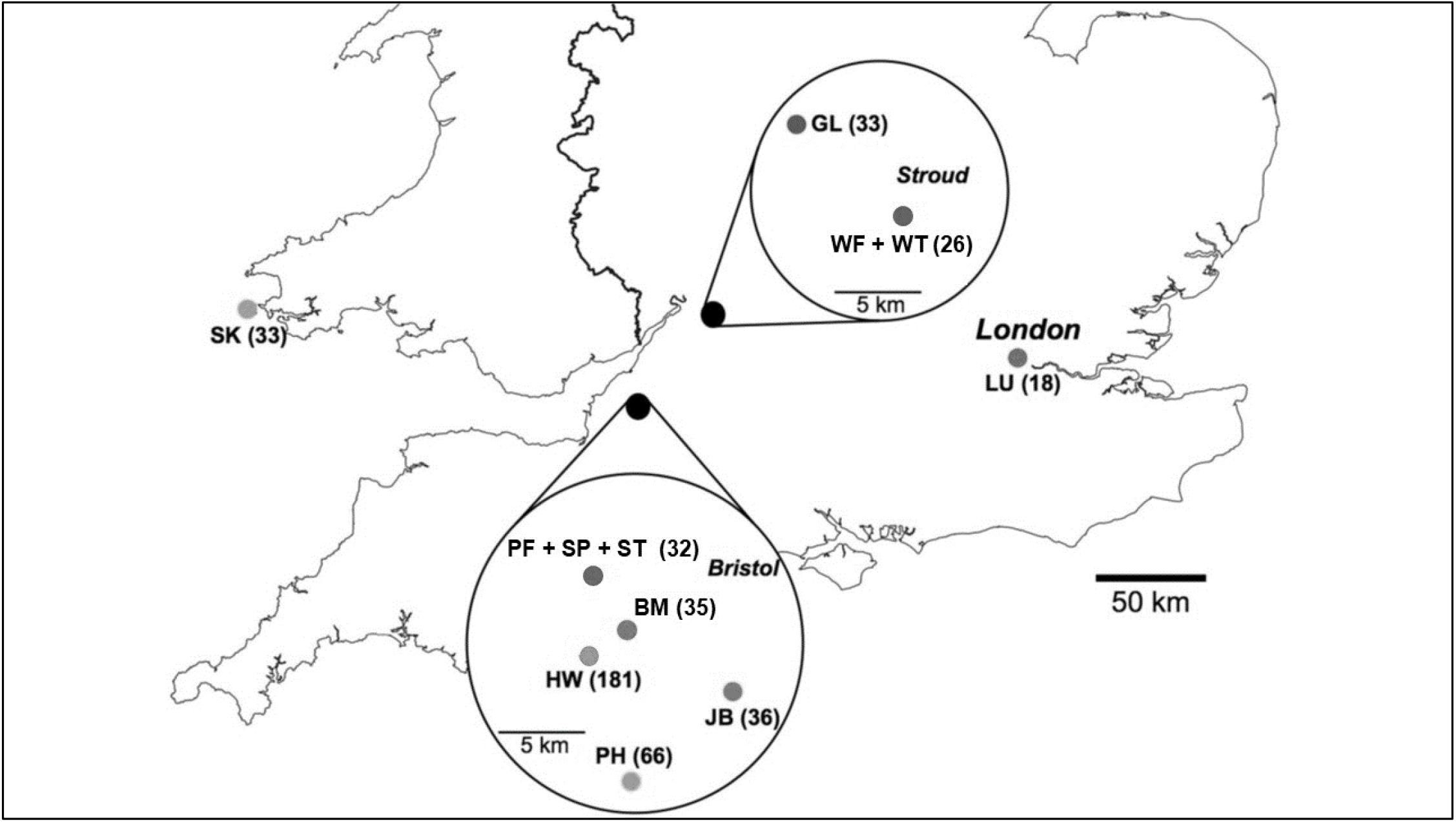
Wild mice sampled from across the southern UK. The 9 sampling sites are shown by letter designations, with the number of animals obtained at each site shown in parentheses. Figure adapted from Abolins et al., 2018.

For some of the significant results obtained from our model selection analysis we then plotted the raw, untransformed immune data against mouse genotype and analysed these untransformed data by ANOVA with sequential Bonferroni corrections. Multiple comparison tests were then performed using Tukey–Kramer HSD test.

All statistical modelling was performed using R version 4.0.5 (R Core Team, 2021).

## 3 Results

We genotyped 435 mice for 23 SNPs in 21 immune-related loci; 14 of these 23 SNPs were found to be polymorphic. We successfully genotyped 378 (87%) mice for all 14 SNPs; the remaining 57 (13%) mice were successfully genotyped for at least 11 SNPs. The minor allele frequencies and the observed and expected heterozygosity are shown in Supporting Information 7 and 8, respectively. The distribution of each SNP among sample sites is shown in Supporting Information 9.

There was extensive variation in the MHC loci within our wild mouse population. H2-Ab was the most diverse H-2 locus, with 3 different genotypes present at similar frequencies and with a heterozygosity of 0.33. For H2-Aa and H2-Eb, one genotype dominated (found in 85% and 70% of mice, respectively) and with lower heterozygosity (0.09 and 0.21, respectively).

For 4 of the cytokine coding loci (IL-1a, IL-1b_U, IL-17a_U and IL-17F) there was relatively abundant genetic diversity. Three IL-1a genotypes occurred in similar frequencies with an overall heterozygosity of 0.34. IL-1b_U, IL-17a_U and IL-17F each had one high frequency genotype (48 – 56 %) but with 2 other common genotypes; heterozygosity was 0.12 –0.28. In contrast, IL-6, IL-10, IL-13, IL-17a_N and TNF were each dominated by one very common genotype (89 – 95%), and had low heterozygosity (0.01 – 0.06). A similar situation pertains for the two TLR loci where one genotype dominated (95% and 96% for TLR5 and TLR9, respectively) with a heterozygosity of 0.04.

The degree of variation at different loci differs among sampling sites (Figure 1, Supporting Information 9). For example, mice from the island site SK (N=27) are invariant at all MHC SNPs, mice from sites BM, HW, JB, LU and WF+WT were polymorphic at 1 or 2 MHC SNPs, and mice from sites GL, PH and ST+SP+PH were polymorphic at all 3 MHC loci. For SNPs in the cytokine coding loci, IL-1a and IL-17F were polymorphic at all sampling sites; for IL-1b_U and IL-17a_U polymorphism was limited to only some sample sites (BM, HW, JB, LU, PH, SK, ST+SP+PH and, all except WF+WT, respectively). The IL-6, IL-10, TNF and TLR loci generally had low levels of polymorphism that was often restricted to a few sample sites (including just one site for TLR-9).

### 3.2 Genetic effects on immune phenotypes

We tested the effect of both non-genetic factors and variation within MHC, cytokine and TLR coding loci on the immune phenotype of wild mice. We first tested the effect of non-genetic factors, finding strong effects of both site and age on all immune parameters, and of body mass (SMI) on cellular components of the immune phenotype, as in Abolins et al. (2018) (Table 2). We then tested the effect of each SNP on the immune phenotype (Tables 2 and 3). Full details of the two-step model selection are reported in Supporting Information 10, effect estimates of all the models selected are shown in Supporting Information 11, and the p-values of the log-likelihood ratio test after deletion testing are reported in Supporting Information 12.

**Table 2.**
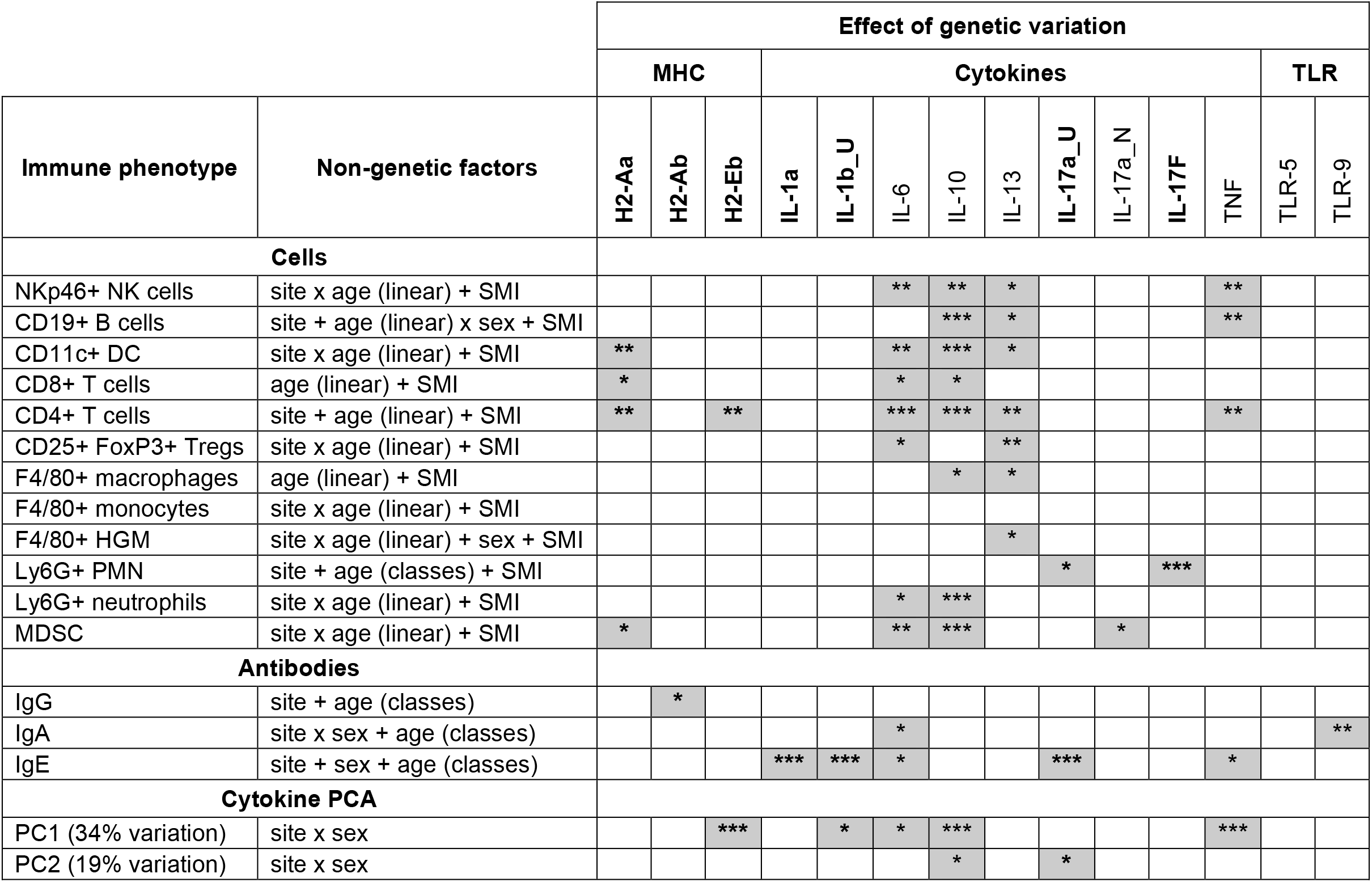
Effect of polymorphism in immune-related loci on immune phenotypes. Results of linear models where first the effect of different age functions (linear, classes), sex (female *vs.* male), site (9 sites) and Scaled Mass Index were tested to select the base model of non-genetic factors. Secondarily, the effect of each polymorphism was tested and shaded boxes indicate where genetic effects were selected in the model (Supporting Information10). The stars shows the significance of the selected genetic effects, determined by deletion testing and log-likelihood ratio test, as: *** for p<0.001; ** for p<0.01; * for p<0.05 and - for p>0.05 (Supporting Information 12). Loci where the minor allele frequency (MAF) is > 0.1 are shown in bold (Supporting Information 7); loci with a MFA < 0.1 are not bold and these results should be interpreted with more caution. DC is Dendritic Cells; F4/80+ HGM is F4/80+ hyper-granulocytic myeloid cells; Ly6G+ PMN is Ly6G+ polymorphonuclear cells; MDSC is Myeloid-Derived Suppressor Cells.

|  |  | Effect of genetic variation |  |  |  |  |  |  |  |  |  |  |  |  |  |
| --- | --- | --- | --- | --- | --- | --- | --- | --- | --- | --- | --- | --- | --- | --- | --- |
|  |  | MHC |  |  | Cytokines |  |  |  |  |  |  |  |  | TLR |  |
| Immune phenotype | Non-genetic factors | H2-Aa | H2-Ab | H2-Eb | IL-1a | IL-1b_U | IL-6 | IL-10 | IL-13 | IL-17a_U | IL-17a_N | IL-17F | TNF | TLR-5 | TLR-9 |
| Cells |  |  |  |  |  |  |  |  |  |  |  |  |  |  |  |
| NKp46+ NK cells | site x age (linear) + SMI |  |  |  |  |  | ** | ** | * |  |  |  | ** |  |  |
| CD19+ B cells | site + age (linear) x sex + SMI |  |  |  |  |  |  | *** | * |  |  |  | ** |  |  |
| CD11c+ DC | site x age (linear) + SMI | ** |  |  |  |  | ** | *** | * |  |  |  |  |  |  |
| CD8+ T cells | age (linear) + SMI | * |  |  |  |  | * | * |  |  |  |  |  |  |  |
| CD4+ T cells | site + age (linear) + SMI | ** |  | ** |  |  | *** | *** | ** |  |  |  | ** |  |  |
| CD25+ FoxP3+ Tregs | site x age (linear) + SMI |  |  |  |  |  | * |  | ** |  |  |  |  |  |  |
| F4/80+ macrophages | age (linear) + SMI |  |  |  |  |  |  | * | * |  |  |  |  |  |  |
| F4/80+ monocytes | site x age (linear) + SMI |  |  |  |  |  |  |  |  |  |  |  |  |  |  |
| F4/80+ HGM | site x age (linear) + sex + SMI |  |  |  |  |  |  |  | * |  |  |  |  |  |  |
| Ly6G+ PMN | site + age (classes) + SMI |  |  |  |  |  |  |  |  | * |  | *** |  |  |  |
| Ly6G+ neutrophils | site x age (linear) + SMI |  |  |  |  |  | * | *** |  |  |  |  |  |  |  |
| MDSC | site x age (linear) + SMI | * |  |  |  |  | ** | *** |  |  | * |  |  |  |  |
| Antibodies |  |  |  |  |  |  |  |  |  |  |  |  |  |  |  |
| IgG | site + age (classes) |  | * |  |  |  |  |  |  |  |  |  |  |  |  |
| IgA | site x sex + age (classes) |  |  |  |  |  | * |  |  |  |  |  |  |  | ** |
| IgE | site + sex + age (classes) |  |  |  | *** | *** | * |  |  | *** |  |  | * |  |  |
| Cytokine PCA |  |  |  |  |  |  |  |  |  |  |  |  |  |  |  |
| PC1 (34% variation) | site x sex |  |  | *** |  | * | * | *** |  |  |  |  | *** |  |  |
| PC2 (19% variation) | site x sex |  |  |  |  |  |  | * |  | * |  |  |  |  |  |

**Table 3.** The effect of different genotypes on immune phenotypes. The parameter estimates for genetic effects for loci where the minor allele frequency is > 0.1 are shown. Parameter estimates for other factors (age, site, SMI, *etc.*), interactions and other loci are in Supporting Information 11. In Variable the genotype in brackets is the genotype to which the comparison is made. Statistical significance is shown by - for p < 0.1, * for p < 0.05, ** for p < 0.01 and *** for p < 0.001. Degrees of freedom is df. DC is Dendritic Cells; MDSC is Myeloid-Derived Suppressor Cells; Ly6G+ PMN is Ly6G+ polymorphonuclear cells.

| Locus | Immune trait | Model selected | df | Variable | Parameter estimate ± SE | t value | p |
| --- | --- | --- | --- | --- | --- | --- | --- |
| H2-Aa | DCs | H2-Aa + site x age (linear) + SMI | 22 | intercept | 4.75 ± 0.33 | 14.53 | *** |
| | | | | AG (AA) | -0.47 ± 0.16 | -2.85 | ** |
| | | | | GG (AA) | -0.27 ± 0.18 | -1.51 | - |
| | CD4+ T cells | H2-Aa x site + age (linear) + SMI | 19 | intercept | 5.57 ± 0.19 | 29.02 | *** |
| | | | | AG (AA) | 0.27 ± 0.34 | 0.79 | - |
| | | | | GG (AA) | 0.16 ± 0.16 | 1.00 | - |
| | CD8+ T cells | H2-Aa + age (linear) + SMI | 6 | intercept | 5.72 ± 0.11 | 50.27 | *** |
| | | | | AG (AA) | -0.14 ± 0.09 | -1.55 | - |
| | | | | GG (AA) | -0.17 ± 0.07 | -2.44 | * |
| | MDSC | H2-Aa + site x age (linear) + SMI | 22 | intercept | 3.48 ± 0.34 | 10.23 | *** |
| | | | | AG (AA) | -0.39 ± 0.19 | -2.10 | * |
| | | | | GG (AA) | -0.07 ± 0.21 | -0.34 | - |
| H2-Ab | IgG | H2-Ab x age (classes) + site | 18 | intercept | 4.20 ± 0.09 | 47.41 | *** |
| | | | | AG (AA) | -0.25 ± 0.11 | -2.30 | * |
| | | | | GG (AA) | -0.35 ± 0.11 | -3.31 | ** |
| H2-Eb | CD4+ T cells | H2Eb x site + age (linear) + SMI | 18 | intercept | 5.32 ± 0.17 | 30.43 | *** |
| | | | | AG (AA) | 0.37 ± 0.20 | 1.85 | . |
| | | | | GG (AA) | 0.44 ± 0.13 | 3.44 | *** |
| | Cytokines PC1 | H2Eb + site x sex | 20 | intercept | -6.76 ± 1.75 | -3.87 | *** |
| | | | | AG (AA) | -1.75 ± 1.06 | -1.66 | . |
| | | | | GG (AA) | -0.04 ± 1.02 | -1.02 | - |
| IL-1a | IgE | IL1a x site + age (classes) + sex | 25 | intercept | 3.23 ± 0.28 | 11.74 | *** |
| | | | | TC (CC) | -1.08 ± 0.29 | -3.69 | *** |
| | | | | TT (CC) | -0.67 ± 0.25 | -2.66 | ** |
| IL-1b_U | Cytokines PC1 | IL1b_U + site x sex | 20 | intercept | -5.45 ± 1.68 | -3.24 | ** |
| | | | | GT (GG) | 0.33 ± 0.84 | 0.40 | - |
| | | | | TT (GG) | -1.78 ± 1.00 | -1.78 | . |
| | IgE | IL1b_U + site + sex + age (classes) | 15 | intercept | 2.05 ± 0.12 | 17.53 | *** |
| | | | | GT (GG) | 0.08 ± 0.07 | 1.15 | - |
| | | | | TT (GG) | 0.31 ± 0.09 | 3.47 | *** |
| IL-17a_U | Ly6G+ PMN | IL-17a_U x site + age (classes) + SMI | 21 | intercept | 3.44 ± 0.31 | 11.25 | *** |
|  |  |  |  | GA (AA) | -0.27 ± 0.27 | -1.00 | - |
|  |  |  |  | GG (AA) | 0.20 ± 0.37 | 0.56 | - |
|  | IgE | IL-17a_U x site + age (classes) + sex | 23 | intercept | 2.38 ± 0.09 | 27.72 | *** |
|  |  |  |  | GA (AA) | -0.39 ± 0.44 | -0.90 | - |
|  |  |  |  | GG (AA) | -0.96 ± 0.20 | -4.79 | *** |
|  | Cytokines PC2 | IL-17a_U + site x sex | 20 | intercept | -3.36 ± 1.06 | -3.17 | *** |
|  |  |  |  | GA (AA) | 1.27 ± 0.92 | 0.17 | - |
|  |  |  |  | GG (AA) | -0.98 ± 0.90 | -1.09 | - |
| IL-17F | Ly6G+ PMN | IL-17F x site + age (classes) + SMI | 28 | intercept | 2.55 ± 0.73 | 3.50 | *** |
|  |  |  |  | CT (CC) | 1.15 ± 0.75 | 1.52 | - |
|  |  |  |  | TT (CC) | 0.53 ± 0.76 | 0.70 | - |

#### 3.2.1 Antibody concentration

We found a strong effect of variation in MHC and cytokine coding loci on antibody concentration (Table 2; Figure 2). IgE concentration was strongly affected by IL-1a and IL-1b_U polymorphisms (p<0.001 for both), with a similar pattern for each. Specifically, mice with the homozygous TT genotype for both IL-1a and IL-1b_U had the highest concentrations of IgE, while those with the other homozygous genotype (CC for IL-1a and GG IL-1b_U) had the lowest IgE concentrations; heterozygous individuals had intermediate IgE concentrations (ANOVA for IL-1a F_2,406_= 10.94, p<0.0001; for IL-1b_U F_2,406_= 4.825, p=0.008; Figure 2A and B). This pattern is suggestive of an allele dosage effect. The concentration of IgE was also affected by the IL-17a_U polymorphism (Table 2), where heterozygous mice had lower concentrations of IgE than either homozygous genotype (ANOVA F_2,398_=3.999, p=0.019; Figure 2C). Finally, the MHC H2-Ab polymorphism showed an effect on IgG concentration (Table 2), where heterozygous mice had lower concentrations than the homozygous GG genotype (ANOVA F_2,391_= 5.831, p=0.003; Figure 2D). While these patterns are evident in the whole data set, effects of IL-1a or IL-17a_U polymorphisms on IgE concentration and of H2-Aa polymorphism on IgG concentration also varied among sites and among mouse age classes (Table 3, Supporting Information 11).

**Figure 2.**
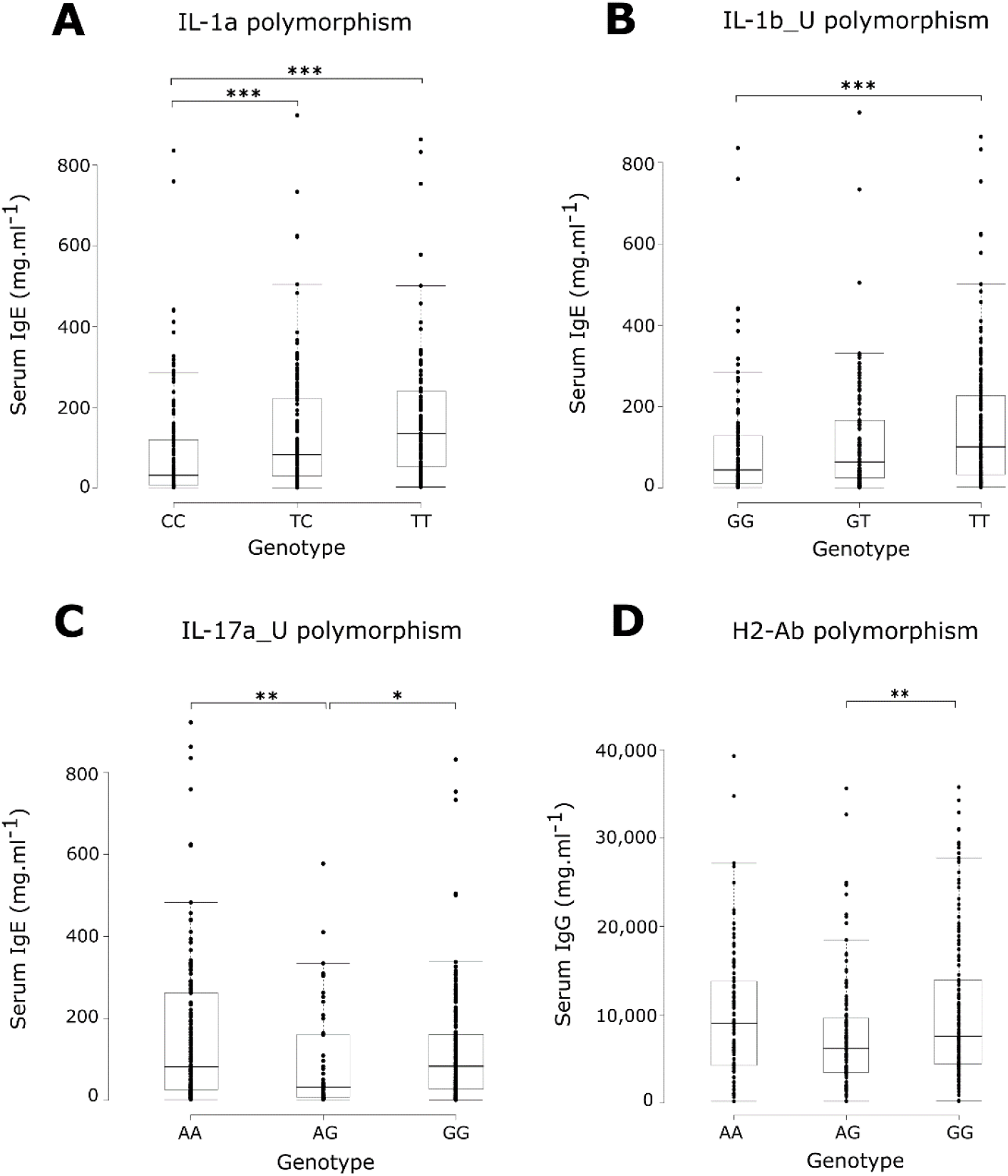
Ige and IgG concentration in mice of different IL-1a, IL-1b, IL-17a and H2-Ab genotypes. Asterisks indicate the p-values obtained using post-hoc Tukey–Kramer test, * for p < 0.05, ** for p < 0.01 and *** for p < 0.001. In the boxplots, midline is the median, with the upper and lower limits of the box being the third and first quartile (75th and 25th percentile) respectively; whiskers are 1.5 times the interquartile range; data points outside this range are shown individually.

#### 3.2.2 Cytokine production

We also found effects of polymorphism in cytokine coding loci on the PCA of cytokine responses (Table 2). Polymorphism in IL1-b_U affected PC1 (p<0.05), which mainly reflects cytokines IL-1B, IL-10, IL-12p40, IL12p70, IL-3 and IL-4. Specifically, heterozygous GT or homozygous GG mice had higher PC1 values than the most common genotype TT (Table 3). Polymorphism within MHC loci H2-Eb affected PC1 (p<0.001), with heterozygous AG mice having lower PC1 values than the rare AA genotype or the most common GG genotype (Table 3). Polymorphism in IL-17a_U affected cytokine PC2 (p<0.05), which mainly reflects cytokines IL-6, IFN-γ and MIP-2α; heterozygous mice had higher PC2 values than the homozygous genotypes (Table 3).

#### 3.2.3 Immune cells

We found some effects of MHC and cytokine loci polymorphisms on immune cell numbers (Table 2, all p>0.05). H2-Aa polymorphism affects the number of DCs, CD4^+^ and CD8^+^ T cells, and MDSC: mice with the rare AA genotype had more DCs, CD8^+^ T cells and MDSCs than either of the other genotypes (Table 3). In contrast, homozygous H-2Aa AA mice had the fewest number of CD4^+^ T cells and heterozygous mice had the most (Table 3). Similarly, the rare AA H2-Eb genotype was associated with fewer CD4^+^ T cells compared to the other genotypes (Table 3). Finally, there was a some effect of polymorphisms in IL-17F and IL-17a_U on the number of PMNs (Table 2). Mice heterozygous for IL-17F had more PMNs than either homozygous genotype (Table 3) whereas mice with the most common GG IL-17a_U genotype had more PMNs than the other rarer genotypes (Table 3). As above, some of these associations varied among sample sites (Supporting Information 11).

For some loci there was too little genetic variation to allow us to draw strong conclusions about their effect on immune phenotypes. However, we found a recurring tendency for effects of polymorphism in IL-6, IL-10, IL-13 and TNF coding loci on many different components of the immune phenotype (Table 2). This interesting trend would benefit from further study. We did not, however, find any striking effects of the two TLR loci, nor the IL-17a_N cytokine loci, on the immune phenotype.

### 3.3 Genetic effects on infection phenotypes

As above, we first tested the effect of non-genetic factors on infection phenotypes (Table 4). Again, we found strong site and age effects on virus positivity, mite and worm burden, and of SMI on mite and worm burden, as in Abolins et al. (2018) (Table 4). We then tested the effect of each SNP on infection phenotypes (Tables 4 and 5). Full details of the two-step model selection are reported in Supporting Information 10, effect estimates of all the models selected are shown in Supporting Information 11, and the p-values of the log-likelihood ratio test after deletion testing are reported in Supporting Information 12.

**Table 4.**
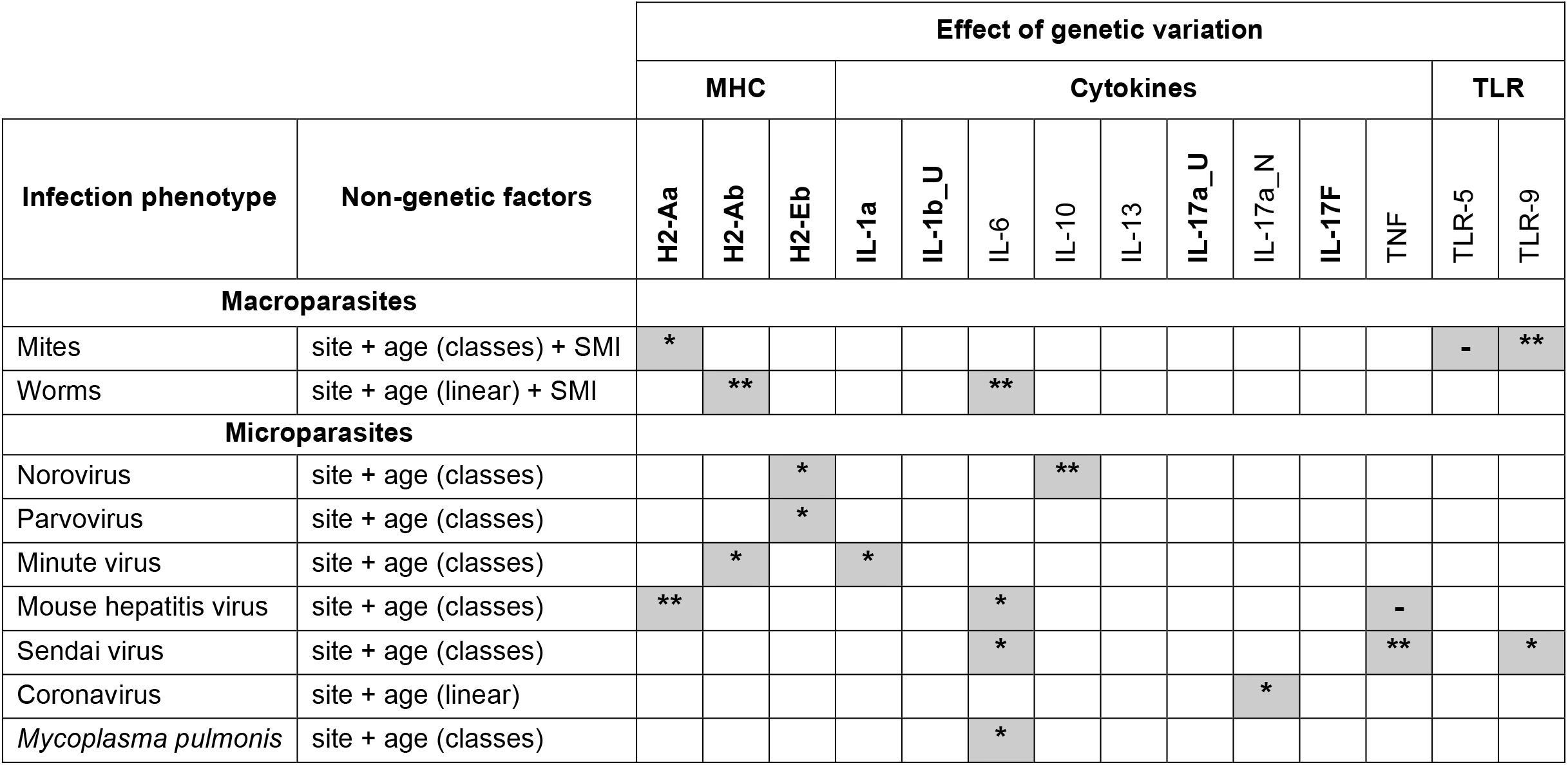
Effect of polymorphism in immune-related loci on infection phenotypes. Results of linear models where first the effect of different age functions (linear, classes), sex (female *vs.* male), site (9 sites) and Scaled Mass Index were tested to select the base model of non-genetic factors. Secondarily, the effect of each polymorphism was tested and shaded boxes indicate where genetic effects were selected in the model (Supporting Information 10). The stars shows the significance of the selected genetic effect, determined by deletion testing and the log-likelihood ratio test, as: *** for p<0.001; ** for p<0.01; * for p<0.05 and - for p>0.05 (Supporting Information 12). Loci where the minor allele frequency (MAF) is > 0.1 are shown in bold (Supporting Information 7); loci with a MFA < 0.1 are not bold and these results should be interpreted with more caution.

**Table 5.** The effect of different genotypes on infection phenotypes. The parameter estimates for genetic effects for loci where the minor allele frequency > 0.1 are shown. Parameter estimates for other factors (age, site, SMI, *etc.*), interactions and other loci are in Supporting Information 11. In Variable the genotype in brackets is the genotype to which the comparison is made. Statistical significance is represented by - for p < 0.1, * for p < 0.05, ** for p < 0.01 and *** for p < 0.001. Degrees of freedom is df.

| Locus | Parasite | Model | df | Variable | Parameter estimate ± SE | t-value | p |
| --- | --- | --- | --- | --- | --- | --- | --- |
| H2-Aa | Mites | H2-Aa + site + age (classes) + SMI | 21 | intercept (0\|50) | 0.33 ± 0.98 | 0.33 | - |
| | | | | intercept (50\|100) | 0.53 ± 0.98 | 0.54 | - |
| | | | | intercept (100\|200) | 1.11 ± 0.98 | 1.14 | - |
| | | | | intercept (200\|300) | 2.24 ± 0.98 | 2.28 | * |
| | | | | intercept (300\|400) | 2.62 ± 0.99 | 2.66 | ** |
| | | | | intercept (400\|500) | 2.75 ± 0.99 | 2.79 | ** |
| | | | | intercept (500\|1000) | 3.63 ± 0.99 | 3.66 | *** |
| | | | | intercept (1000\|1500) | 4.78 ± 1.01 | 4.75 | *** |
| | | | | AG (AA) | 1.91 ± 0.76 | 2.50 | * |
| | | | | GG (AA) | 1.80 ± 0.84 | 2.15 | * |
| | Mouse hepatitis virus | H2-Aa + site + age (classes) | 15 | intercept (0\|1) | -1.90 ± 1.88 | -1.01 | - |
| | | | | intercept (1\|2) | -0.02 ± 1.87 | -0.01 | - |
| | | | | intercept (2\|3) | 1.61 ± 1.88 | 0.86 | - |
| | | | | AG (AA) | -2.64 ± 1.97 | -1.34 | - |
| | | | | GG (AA) | -0.04 ± 1.84 | -0.02 | - |
| H2-Ab | Worms | H2-Ab x site + age (linear) + SMI | 25 | intercept | 0.69 ± 0.22 | 3.15 | ** |
| | | | | AG (AA) | -0.10 ± 0.24 | -0.42 | - |
| | | | | GG (AA) | -0.12 ± 0.41 | -0.29 | - |
| | Minute virus | H2-Ab + site + age (classes) | 15 | intercept (0\|1) | 1.59 ± 0.52 | 3.03 | ** |
| | | | | intercept (1\|2) | 2.99 ± 0.55 | 5.40 | *** |
| | | | | intercept (2\|3) | 3.82 ± 0.59 | 6.53 | *** |
| | | | | AG (AA) | -0.25 ± 0.45 | -0.55 | - |
| | | | | GG (AA) | 0.76 ± 0.45 | 1.69 | - |
| H2-Eb | Norovirus | H2-Eb x age (classes) + site | 19 | intercept (0\|1) | 1.90 ± 0.64 | 2.95 | ** |
| | | | | intercept (1\|2) | 4.41 ± 0.69 | 0.69 | *** |
| | | | | intercept (2\|3) | 6.17 ± 0.77 | 0.77 | *** |
| | | | | AG (AA) | -0.64 ± 0.57 | -1.13 | - |
| | | | | GG (AA) | 0.42 ± 0.46 | 0.92 | - |
| | Parvovirus | H2-Eb + site + age (classes) | 16 | intercept (0\|1) | -1.25 ± 0.51 | -2.43 | * |
| | | | | intercept (1\|2) | 0.18 ± 0.51 | 0.36 | - |
| | | | | intercept (2\|3) | 0.97 ± 0.51 | 1.92 | . |
|  |  |  |  | intercept (3\|4) | 3.13 ± 0.54 | 5.85 | *** |
|  |  |  |  | AG (AA) | 1.02 ± 0.40 | 2.51 | ** |
|  |  |  |  | GG (AA) | 0.56 ± 0.38 | 1.48 | - |
| IL-1a | Minute virus | IL1a + site + age (classes) | 15 | intercept (0\|1) | 2.56 ± 0.66 | 3.86 | *** |
|  |  |  |  | intercept (1\|2) | 3.94 ± 0.69 | 5.74 | *** |
|  |  |  |  | intercept (2\|3) | 4.73 ± 0.71 | 6.62 | *** |
|  |  |  |  | TC (CC) | 1.10 ± 0.46 | 2.37 | * |
|  |  |  |  | TT (CC) | 0.75 ± 0.52 | 1.43 | - |

We found a strong effect of MHC H2-Aa polymorphism on mite burden: mice with the rare homozygous AA genotype had fewer mites than those with the other more common genotypes (Tables 4 and 5, p<0.05). MHC H2-Ab polymorphism had an effect on worm burden (Table 4, p<0.01), with a general tendency for mice with the AA genotype to have higher worm burdens than the two other genotypes. However this general pattern obscures some local variation between sites (see all effect estimates of the H2-Ab x site interaction in Supporting Information 11); for instance H2-Ab heterozygous mice from site WF+WT have higher worm burdens than the other genotypes.

We also found effects of polymorphism in MHC loci on seropositivity for virus infections (Table 4). For example, mice with AA or AG H2-Ab genotypes were less likely to be seropositive for Minute virus compared to mice with the most common GG genotype (Table 5, p<0.05); mice that were heterozygous at the H2-Aa locus were less likely to be seropositive for Mouse Hepatitis virus (MHV) infection than either of the homozygous genotypes (Table 5, p<0.01). Finally, mice heterozygous at the H2-Eb locus were less likely to be seropositive for norovirus – but more likely to be seropositive for parvovirus – compared with the two homozygous genotypes (Table 5, p<0.05 for both).

We found only limited evidence of associations between polymorphisms in cytokine coding loci and infection phenotypes (Tables 4 and 5), though mice that were heterozygous at the IL-1a locus were more likely to be seropositive for Minute virus genotypes (p<0.05). Finally, we found a strong effect of IL-6 on both macroparasite (worms number, p<0.01) and microparasite infection markers (MHV, sendai virus and *M. pulmonis;* Table 4, p<0.05 for all). We also found an effect of the TLR-9 locus on mite infection phenotype (Table 4, p<0.01). However, for these loci the relative abundance of the different genotypes means that these results should be considered with caution, and would benefit from further study.

## 4 Discussion

We tested the effect of polymorphism in 14 immune-related loci on a wide range of immune and infection phenotypes in wild UK house mice, *Mus musculus domesticus*. These results bring new insights to the field of immunogenetics in three ways. First, this is a first study, of which we are aware, where the effect of a large number of immune-related loci (MHC, cytokines and TLR) on multiple immune and infection phenotypes have been studied in a wild mammal. Second, these effects have been characterized in a large sample of mice from 9 different sample sites, which is rare in a wild species where field constrains often limits sample sizes. Third, these results are of particular interest because they also consider infection phenotypes, which here are the result of natural processes of infection, co-infection and host immune responses occurring in these populations.

Most of the immune-related loci that we investigated were polymorphic, though there were differences in the representation of genotypes at loci among mice from different sites. Overall the observed genetic variation was most notable for the MHC and cytokine coding loci, but was much less extensive for the TLR coding loci, which may explain why we found comparatively fewer clear effects of polymorphism in TLR coding loci. We did, however, find strong effects of the MHC and cytokine coding loci on different components of the immune phenotype – principally on antibody concentration and on cytokine production – but also on infection phenotypes. The effects of the MHC loci on immune and infection state are consistent with previous results in natural populations (as Table 1). Interestingly, we found that polymorphism in the MHC H2-Ab locus affected both IgG concentration and intestinal worm burden and there is the possibility that this is causal. Associations between MHC loci and antibody response in wild populations have been described in other mammals such as Soay sheep (Huang et al., 2021; but also see in other species reported in Gaigher et al., 2019). We also found effects of the MHC H2-Aa locus on various populations of DCs, CD4+ and CD8+ T cells, and MDSCs as well as on ectoparasite burden.

We found major effects of polymorphisms in cytokine coding loci on antibody concentration. Specifically, the serum concentration of IgE was affected by polymorphism in IL-1a, IL1b_U, IL-6 IL-17a_U, and TNF, with different IL-1a, IL1b_U, IL-17a_U genotypes having quite strikingly different concentrations (Figure 2). Polymorphisms in IL-6 also affected the concentration of faecal IgA. The potential mechanism of these effects on antibody concentration are that these polymorphism affect (i) animals’ antibody production *per se*, (ii) animals’ resistance or susceptibility to a wide range of infections, which is manifest as different antibody concentrations, or (iii) a combination of these.

Among the cytokine-coding loci that we investigated we found a range of effects on cell populations, on the concentration of antibodies, and on cytokine PCs. Many cytokines are multifunctional and widely connected in the generation on an immune response (Fonseca dos Reis et al., 2021), such that we should not be surprised that genetic effects of cytokine-coding loci may have pleiotropic phenotypic effects. Five cytokine coding loci (IL-1a, IL-1b, IL-6, IL-17a and TNF) affected the concentration of IgE, and an overlapping set of 5 (IL-1b, IL-6, IL-10, IL-17a, and TNF) affected cytokine PCs (Table 2). Moreover, a further combination of 5 of these (IL-1a, IL-6, IL-10, IL-17a, and TNF) also affected measures of infection (Table 4). The associations between these genotypes and both immune and infection phenotypes might be the result of causative relationships, although further evidence would be need to substantiate this. Polymorphism in IL-10 appears to have pleiotropic immune and infection effects and this is consistent with IL-10 being known to play a central role in regulating immune responses to a wide range of micro- and macroparasites (Couper et al., 2008). Similarly, polymorphism in IL-6 also appears to be pleiotropic, which is also consist with IL-6’s role in integrating different aspects of an immune response, and where in humans suppression of IL-6 function increases the risk of serious infection (Rose-John et al., 2017). However, we need to be cautious in interpreting these data, since the measures of microparasite infection are serological, and so may be confounded with individuals’ antibody production *per se*, which we also found to be affected by polymorphism in these and other loci.

Our measures of viral infection was serological, and while this is an easy-to-use approach it is potentially problematic because it is itself an immunological measure. An advantage of serological diagnosis is that it shows evidence of historical infection, though in the mice studied here their median age is 6 – 7 weeks (maximum 20 – 39, for male and females, respectively; Abolins et al., 2017), meaning that in many cases these serological diagnoses are often likely of relatively recent viral infection. Our study could be improved by using specific viral diagnostic approaches, and indeed by considering a wider-still range of parasites but also of pathology.

The relationships that we have found do not provide strong support for the notion of heterozygote advantage; this is in line with the inconclusive evidence for this phenomenon in other wild mammals (Table 1). Rather, we found that some rare homozygous genotypes are less parasitized than more common homozygous or heterozygous genotypes; for example, mice with the rare AA MHC H2-Aa genotype had fewer mites than the other two genotypes. Parasites are thought to impose a significant selection pressure on their hosts and thus represent the major component of balancing selection influencing MHC polymorphism, which can result in patterns such as we have seen (Bryja et al., 2006).

While the present study focusses on genetic effects on measures of the immune response, previous analysis of this same data set (Abolins et al., 2018) has revealed how non-genetic factors including body condition, age, infection status, and season also affect immune state; this was confirmed in our analysis here. Taken together, these studies reveal the complexity of genetic and non-genetic factors affecting immune status and infection phenotypes in wild animals. Although it is difficult to compare the relative magnitude of the genetic and non-genetic effects, the genetic effects on antibody concentrations and cytokine production that we have observed appear to be especially significant.

For some of the immune-related loci (including IL-6, IL-10, IL-13, TNF, TLR) there was a substantial imbalance in the frequency of different genotypes that limited our analytical power. Previous studies have noted that effective sample sizes can limit what can be studied in wild populations and have suggested minimum sample sizes of at least 200 individuals (Gaigher et al., 2019). It is notable that even with our large sample size of more than 400 mice, the relative under-representation of some genotypes has constrained the power of our analysis. Previous genetic analyses of these same mice using putatively neutral loci showed genetic differentiation among mice from the different sites (Abolins et al., 2018), again consistent with differences in genotype frequencies in the immune-related loci that we have observed. Moreover, previous analyses of the immune distance among these mice found that mice within one sample site were on average more immunologically similar to each other, compared with mice from other sample sites (Abolins et al., 2018). In agreement with both of these analyses, in the present study we also found significant effects of sample site on immune and infection phenotypes. Genetic and immunological differences among mice at different sites is therefore a notable feature of the mouse populations that we have studied. This host genetic and immunological diversity therefore presents a heterogeneous environment in which parasites are selected and are evolving, and where hosts continue to evolve to respond to that threat.

## Supporting information

All supplemental files

## Data accessibility

All data will be deposited in Dryad.

## Acknowledgements

This work was supported by NERC (NE/I022892/1). We would like to thank the farm owners and London Underground for generous access to their properties. We also would like to thank Jane Hurst and Steve Paterson for helpful discussions.

## Author Contributions

Conceived and designed the experiments: EMR, MV. Performed the experiments: LL, EMR, MV. Analysed and interpreted the data: LC, LL, EMR, MV. Wrote the paper: LC, EMR, MV.

