## Supplementary material for "The genetics of immune and infection phenotypes in wild mice, *Mus musculus domesticus*": All supplemental files

#### Table of Contents:


| | |
|---|---|
| **1. Distribution of the wild mice by site, sex, and age class.** | Page 2 |
| **2. The immune-related loci successfully amplified and detection of SNPs.** | Page 3 |
| **3. Target loci.** | Page 4 |
| **4. SNPs identified in the pilot sample of 25 wild mice.** | Page 5 |
| **5. Variants detected in the pilot study of 25 wild mice for (A) exonic and (B) non-exonic regions.** | Page 6 |
| **6. Correlations of the 36 cytokine measures with the PCA axes 1 and 2 (PC1 and PC2, respectively), together accounting for 53% of the total variation.** | Page 10 |
| **7. The minor allele frequencies for the 14 immune-related SNPs.** | Page 11 |
| **8. Heterozygosity of the 14 immune-related SNPs.** | Page 12 |
| **9. The number of SNP genotypes at different sample sites.** | Page 13 |
| **10. Selection of the best explanatory models for immune and infection phenotype response variables.** | Page 14 |
| **11. Parameter estimates of the selected generalized linear models of immune and infection phenotype in wild mice.** | Page 25 |
| **12. P-values of the deletion and the log-likelihood ratio test, assessing the magnitude of the genetic effect retained in the model selection process.** | Page 33 |

Supplementary 1. Distribution of the wild mice by site, sex, and age class. The three age classes used were: 0-6 weeks, immature animals less likely to be venturing far from their nest site; 7-12 weeks, young animals that are less likely to have reproduced; >12 weeks, likely sexually active adults. Age was missing for four individuals (column “NA”). Site is as Figure 1 and Abolins et al., 2018.

| | | **Sex** | | **Age Class** | | | |
|---|---|---|---|---|---|---|---|
| **Site** | **N** | **Females** | **Males** | **Adults** | **Young** | **Immature** | **NA** |
| BM | 35 | 17 | 18 | 13 | 15 | 7 | 0 |
| GL | 33 | 19 | 14 | 14 | 8 | 11 | 0 |
| HW | 181 | 81 | 100 | 27 | 74 | 78 | 2 |
| JB | 36 | 17 | 19 | 8 | 12 | 16 | 0 |
| LU | 18 | 5 | 13 | 12 | 0 | 6 | 0 |
| PH | 66 | 22 | 44 | 22 | 21 | 23 | 0 |
| SK | 33 | 21 | 12 | 13 | 10 | 10 | 0 |
| PF+SP+ST | 32 | 13 | 19 | 5 | 12 | 13 | 2 |
| WF+WT | 26 | 16 | 10 | 10 | 5 | 11 | 0 |
| **Total** | **460** | **211** | **249** | **124** | **157** | **175** | **4** |

Supplementary 2. The immune-related loci successfully amplified and detection of SNPs.

| Locus | PCR amplification success | SNP names | Polymorphic (Yes/No) |
|---|---|---|---|
| IL-1a | Yes | IL-1a | Yes |
| IL-1b | Yes | IL-1b_U | Yes |
| | | IL-1b_N | No |
| IL-2 | Yes | IL-2 | No |
| IL-4 | Yes | IL-4 | No |
| IL-6 | Yes | IL-6 | Yes |
| IL-10 | Yes | IL-10 | Yes |
| IL-12a | Yes | IL-12a | No |
| IL-12b | Yes | IL-12b | No |
| IL13 | Yes | IL-13 | Yes |
| IL-17a | Yes | IL-17a_N | Yes |
| | | IL-17a_U | Yes |
| IL-17F | Yes | IL-17F | Yes |
| IFNγ | Yes | IFNγ | No |
| TNF | Yes | TNF | Yes |
| IL-2rg | Yes | IL-2rg | No |
| CD40LG | Yes | CD40LG | No |
| TLR-4 | Yes | TLR-4 | No |
| TLR-5 | Yes | TLR-5 | Yes |
| TLR-9 | Yes | TLR-9 | Yes |
| H2-Aa | Yes | H2-Aa | Yes |
| H2-Ab1 | Yes | H2-Ab1 | Yes |
| H2-Eb1 | Yes | H2-Eb1 | Yes |
| H2-K1 | Did not amplify | - | - |
| H2-D1 | Did not amplify | - | - |
| **23 loci** | **21 loci successfully amplified** | **23 SNPs found** | **14 SNPs are polymorphic** |

Supplementary 3. Target loci. The target regions for amplification of 24 loci, showing the linkage group (LG), exons, and the Start and End nucleotide limits of the targeted amplification. The sequence of the forward (F) and reverse (R) primer are shown, based on the sequence of C57/BL6 mice. Annealing is the temperature used in the PCR amplification; Size is the predicted product size in bp. For IL-12a, there are three splice variants and the region targeted recognises the final exon, which is as shown for the three variants. H2-K1 and H2-D1 failed to amplify using these primers and so are not considered further.

| Locus | LG | Exon | Start | End | Primer F | Primer R | Annealing | Size |
|---|---|---|---|---|---|---|---|---|
| IL-1a | 2 | 4 | 129306468 | 129306667 | CAGATCATGGGTTATGGACTGC | TCCTTCTATGATGCAAGCTATGG | 58.1-58.8 | 199 |
| IL-1b | 2 | 6-7 | 129364613 | 129366065 | TCAAAGCAATGTGCTGGTGC | AGGAGAACCAAGCAACGACA | 59.9-59.9 | 1452 |
| IL-2 | 3 | 4 | 37120725 | 37121193 | GGAGAGCTTTATTTCTTGAAAACAC | TCTGACAACACATTTGAGTGCC | 57.1-59.3 | 468 |
| IL-4 | 11 | 1-2 | 53618173 | 53618647 | CGTTGCTGTGAGGACGTTTG | AAACTTAATTGTCTCTCGTCACTG | 57.2-60.0 | 474 |
| IL-6 | 5 | 5 | 30019402 | 30019833 | GTCCTTCCTACCCCAATTTCCA | TCCAAGAAACCATCTGGCTAGG | 59.6-59.7 | 431 |
| IL-10 | 1 | 3-4 | 131021333 | 131022552 | ACTTGGGTTGCCAAGCCTTA | TATTAAAATCACTCTTCACCTGCTC | 57.1-59.9 | 1219 |
| IL-12a | 3 | 6,7,8 | 68697895 | 68698394 | TCTCTGAATCATAATGGCGAGACT | TTTTAAATAAGGGGTGACTGAGTGT | 58.3-59.4 | 499 |
| IL-12b | 11 | 6-7 | 44411101 | 44412635 | TGAAGGAGACAGAGGAGGGG | GAACACATGCCCACTTGCTG | 59.9-60.0 | 1534 |
| IL-13 | 11 | 1 | 53634494 | 53634693 | CTGGTCTTGTGTGATGTTGCTC | CAGCCTAGGCCAGCCCAC | 59.7-62.5 | 199 |
| IL-17a | 1 | 3 | 20733648 | 20734449 | CCTCTGTGATCTGGGAAGCTC | CAGAGTAGGGAGCTAAATTATCCA | 57.3-59.8 | 801 |
| IL-17F | 1 | 3 | 20777324 | 20777953 | TGTGTGCTTCTTCCTTGCCA | CAGACACTCAGGCTGCATCA | 60.0-60.1 | 629 |
| IFNγ | 10 | 2-3 | 118442463 | 118442793 | TCAAGTGGCATAGATGTGGAAGA | CAAACTTGGCAATACTCATGAATGC | 59.4-59.7 | 330 |
| TNF | 17 | 1 | 35201656 | 35201995 | TCATCCCTTTGGGGACCGAT | CTCAGCGAGGACAGCAAGG | 60.4-60.6 | 339 |
| IL-2rg | X | 8 | 101264439 | 101265001 | GGTAGAAAAAGGGAGGGAGAATCC | CTGCAGCCAGACTACAGTGA | 59.3-60.3 | 562 |
| CD40LG | X | 5 | 57223215 | 57224026 | ACAGTGGGCCAAGAAAGGAT | GGAGCCCAGGTCAACCATAA | 59.2-59.3 | 811 |
| TLR-4 | 4 | 3 | 66840072 | 66841071 | AGTGGCCCTACCAAGTCTCA | GCGGGGCACTCCTTCTTCTA | 60.1-61.6 | 999 |
| TLR-5 | 1 | 4 | 182973251 | 182973739 | CAGACGGCAGGATAGCCTTT | TGCAGAGGCTCGAGTTCATC | 59.8-59.8 | 488 |
| TLR-9 | 9 | 2 | 106224312 | 106225324 | ATGTGGCCAAAAGTCCCTCC | ATACCGTTGCCGCTGAAGTC | 60.2-60.7 | 1012 |
| H2-Aa | 17 | 2-3 | 34283580 | 34284533 | TGTTTCAGAACCGGCTCCTC | ACCACGTAGGCACCTATGGTA | 59.9-60.3 | 953 |
| H2-Ab1 | 17 | 3-4 | 34267339 | 34268002 | AGCCCAATGTCGTCATCTCC | TGTGACGGATGAAAAGGCCA | 59.8-59.8 | 663 |
| H2-Eb1 | 17 | 3 | 34314174 | 34314406 | GCCTACGGTGACTGTGTACC | GCTGGGATGCTCCACCTG | 60.1-60.1 | 232 |
| H2-K1 | 17 | 2-3 | 33999288 | 33999972 | GCGTTCCCGTTCTTCAGGTA | TGAGGTATTTCGTCACCGCC | 60.0-60.1 | 684 |
| H2-D1 | 17 | 2-3 | 35263380 | 35264078 | CCCACACTCGATGCGGTATT | GCGTTCCCGTTCTTCAGGTA | 60.0-60.1 | 698 |
| Myo1a | 10 | 8-10 | 127710010 | 127710993 | AGGAGTATGTTATCCCCAACCA | TGTGACATTCAGCACCGTGA | 58.2-59.8 | 983 |

Supplementary 4. SNPs identified in the pilot sample of 25 wild mice. Number shows the number of mice from which data were obtained; Exon and Intron sequence length in bp, and the number of SNPs identified in the Exons and Introns.

| Locus | Number | Exon (bp) | Exon SNPs | Intron (bp) | Intron SNPs |
|---|---|---|---|---|---|
| IL-1a | 23 | 157 | 1 | | - |
| IL-1b | 20 | 572 | 10 | 201 | 10 |
| IL-2 | 22 | 371 | - | | - |
| IL-4 | 22 | 162 | 1 | 191 | |
| IL-6 | 22 | 388 | 1 | | |
| IL-10 | 20 | 99 | - | 667 | 7 |
| IL-12a | 24 | 435 | - | | - |
| IL-12b | 17 | - | - | 783 | 1 |
| IL-13 | 22 | 138 | 2 | | - |
| IL-17a | 15 | 732 | 5 | | - |
| IL-17F | 21 | 564 | 3 | | - |
| IFNγ | 22 | 172 | - | 45 | - |
| TNF | 19 | 272 | 3 | | - |
| IL-2rg | 17 | 507 | - | | - |
| CD40LG | 17 | 636 | - | | - |
| TLR-4 | 21 | 744 | 5 | | - |
| TLR-5 | 21 | 443 | 3 | | - |
| TLR-9 | 21 | 828 | 1 | | - |
| H2-Aa | 13 | 209 | 11 | 290 | 26 |
| H2-Ab1 | 22 | 276 | 14 | 305 | 20 |
| H2-Eb1 | 23 | 187 | 7 | | - |
| H2-K1 | 21 | 252 | 2 | 539 | 10 |

Supplementary 5. Variants detected in the pilot study of 25 wild mice for (A) exonic and (B) non-exonic regions. Locus is the target locus; SNP shows the identified variant; Position is the position within the amplified fragment; Genomic is the location in the mouse genome. Number shows the number of animals as a fraction of the number for whom data were available. Variants in bold were used for KASP genotyping. Lab Mice indicates whether or not the SNP identified in wild mice is known in strains of laboratory mice, as recorded in informatics.jax.org in 2020 In (A) Change is the genomic feature and nature of the variant; S/NS indicated whether the variant is predicted to be code for a synonymous or non-synonymous substitution, respectively. For IL-1b, the SNP at position 129364687 is named IL-1b_U and that at position129365058 IL-1b_N; for IL-17a the SNP at position 20733797 is named IL-17a_N and that at position 20734091 is IL-17a_U; for Myo1a, the SNP at position 127710437 is named Myo1a_1 and that at position 546 is named Myo1a_2.

| **A.** | | | | | | | |
|---|---|---|---|---|---|---|---|
| **Locus** | **SNP** | **Position** | **Genomic** | **Number** | **Change** | **S/NS** | **Lab Mice** |
| **IL-1a** | **C/T** | **93** | **129306612** | **12/23** | **-** | **Syn** | Y |
| **IL-1b** | **A/C** | **21** | **129364687** | **12/20** | **UTR** | **-** | N |
| | G/A | 39 | 129364705 | 1/20 | UTR | - | Y |
| | C/A | 177 | 129364843 | 1/20 | UTR | - | N |
| | C/T | 252 | 129364918 | 1/20 | UTR | - | Y |
| | A/C | 306 | 129364972 | 1/20 | UTR | - | N |
| | A/C | 320 | 129364986 | 1/20 | UTR | - | N |
| | **C/A** | **392** | **129365058** | **1/20** | **Asp>Tyr** | **Non-syn** | N |
| | C/T | 432 | 129365098 | 1/20 | Phe>Phe | Syn | Y |
| | G/A | 435 | 129365101 | 1/20 | Val>Val | Syn | N |
| | T/C | 531 | 129365197 | 1/20 | Phe>Phe | Syn | Y |
| **IL-4** | **C/G** | **220** | **53618503** | **1/22** | **Gly>Ala** | **Non-syn** | N |
| **IL-6** | **C/T** | **92** | **30019545** | **4/22** | **Thr>Thr** | **Syn** | Y |
| **IL-13** | T/C | 21 | 53634563 | 2/22 | - | Syn | Y |
| | **C/T** | **27** | **53634569** | **2/23** | **-** | **Syn** | Y |
| **IL-17a** | **G/A** | **98** | **20733797** | **2/15** | **Glu>Lys** | **Non-syn** | N |
| | A/G | 165 | 20733864 | 2/15 | UTR | - | Y |
| | C/T | 218 | 20733917 | 6/9 | UTR | - | Y |
| | **G/A** | **392** | **20734091** | **7/15** | **UTR** | **-** | Y |
| | A/G | 498 | 20734197 | 3/10 | UTR | - | Y |
| **IL-17F** | A/T | 68 | 20777440 | 1/21 | UTR | - | N |
| | **T/C** | **174** | **20777546** | **11/19** | **UTR** | **-** | Y |
| | T/C | 419 | 20777791 | 1/19 | Lys>Lys | - | Y |
| **TNF** | **G/T** | **114** | **35201817** | **4/19** | **Ala>Ser** | **Non-syn** | N |
| | C/A | 227 | 35201930 | 5/19 | UTR | - | N |
| | G/A | 236 | 35201939 | 2/19 | UTR | - | N |
| **TLR4-** | C/T | 555 | 66840696 | 20/21 | Asn>Asn | Syn | N |
| | C/A | 580 | 68840711 | 1/21 | His>Asn□ | Non-syn | N |
| | A/G | 581 | 68840712 | 1/21 | His>Arg□ | Non-syn | N |
| | **A/C** | **609** | **68840750** | **20/21** | **Glu>Asp** | **Non-syn** | N |
| | T/C | 682 | 68840823 | 1/21 | Leu>Leu | Syn | N |
| **TLR-5** | T/C | 15 | 182973315 | 1/21 | Thr>Thr | Syn | N |
| | **G/A** | **255** | **182973555** | **1/22** | **Leu>Leu** | **Syn** | N |

| | | | | | | | |
|---|---|---|---|---|---|---|---|
| | A/G | 267 | 182973567 | 1/23 | Gly>Gly | Syn | N |
| **TLR-9** | **C/T** | **175** | **106224547** | **1/22** | **Arg>Cys** | **Non-syn** | N |
| **H2-Aa** | G/A | 36 | 34283683 | 2/11 | Tyr>His | Non-syn | N |
| | A/C | 40 | 34283687 | 1/11 | Arg>Arg | Syn | N |
| | G/A | 49 | 34283696 | 1/11 | Phe>Phe | Syn | N |
| | **G/A** | **51** | **34283698** | **2/11** | **Phe>Leu** | **Non-syn** | N |
| | C/T | 61 | 34283708 | 2/11 | Glu>Glu | Syn | N |
| | G/A | 70 | 34283717 | 1/11 | Gly>Gly | Syn | N |
| | T/C | 78 | 34283725 | 1/11 | Thr>Ala | Non-syn | N |
| | G/A | 88 | 34283735 | 3/11 | Ser>Ser | Syn | N |
| | T/A | 121 | 34283768 | 11/12 | Pro>Pro | Syn | N |
| | G/T | 145 | 34283792 | 4/12 | Ile>Ile | Syn | N |
| | A/G | 148 | 34283795 | 5/12 | Leu>Leu | Syn | N |
| | G/A | 178 | 34283825 | 2/13 | Ser>Ser | Syn | N |
| **H2-Ab1** | C/T | 13 | 34267403 | 5/22 | Cys>Cys | Syn | N |
| | A/G | 16 | 34267406 | 10/22 | Ser>Ser | Syn | N |
| | A/G | 34 | 34267424 | 5/22 | Pro>Pro | Syn | N |
| | C/A | 59 | 34267449 | 10/22 | Arg>Arg | Syn | N |
| | **G/A** | **79** | **34267459** | **10/23** | **Gln>Arg** | **Non-syn** | N |
| | C/T | 156 | 34267536 | 3/23 | Glu>Lys | Non-syn | N |
| | A/G | 159 | 34267539 | 7/23 | Met>Val | Non-syn | N |
| | T/G | 160 | 34267540 | 10/23 | Met>Arg | Non-syn | N |
| | A/G | 162 | 34267542 | 10/23 | Thr>Ala | Non-syn | N |
| | C/T | 181 | 34267561 | 1/23 | Val>Ala | Non-syn | N |
| | C/T | 184 | 34267564 | 10/23 | Tyr>Ser | - | N |
| | C/G | 196 | 34267576 | 2/23 | - | - | N |
| | C/T | 543 | 34267923 | 3/22 | - | - | N |
| | C/T | 553 | 34267933 | 1/22 | Ser>Ser | Syn | N |
| **H2-Eb1** | **G/A** | **21** | **34314245** | **3/23** | **Ser>Asn** | **Non-syn** | N |
| | A/G | 45 | 34314269 | 1/23 | Glu>Gly | Non-syn | N |
| | T/C | 64 | 34314288 | 1/23 | Asn>Asn | Syn | N |
| | A/G | 77 | 34314301 | 2/23 | Lys>Glu | Non-syn | N |
| | A/C | 133 | 34314357 | 11/23 | Thr>Thr | Syn | N |
| | A/G | 149 | 34314373 | 2/23 | Thr>Ala | Non-syn | N |
| | A/G | 161 | 34314385 | 2/24 | Ser>Gly | Non-syn | N |
| **Myo1a** | G/A | 308 | 127710379 | 1/21 | Ala>Ser | Syn | Y |
| | C/T | **366** | **127710437** | **2/21** | **Leu>Leu** | **Syn** | N |

| **B.** | | | | |
|---|---|---|---|---|
| **Locus** | **SNP** | **Position** | **Genomic** | **Number** |
| **IL-1b** | G/A | 606 | 129365272 | 1/20 |
| | T/G | 630 | 129365296 | 1/20 |
| | G/C | 644 | 129365310 | 1/20 |
| | C/T | 651 | 129365317 | 8/19 |
| | A/G | 667 | 129365333 | 8/19 |
| | G/A | 674 | 129365340 | 10/19 |
| | A/T | 695 | 129365361 | 8/19 |
| | A/G | 697 | 129365363 | 1/19 |
| | C/T | 739 | 129365405 | 8/19 |
| | C/A | 767 | 129365433 | 1/19 |

| | | | | |
|---|---|---|---|---|
| **IL-10** | T/C | 138 | 131021519 | 4/20 |
| | **T/C** | **139** | **131021520** | **4/20** |
| | C/T | 323 | 131021704 | 3/20 |
| | C/T | 383 | 131021764 | 1/20 |
| | G/A | 500 | 131021881 | 3/20 |
| | G/A | 540 | 131021921 | 1/20 |
| | T/G | 628 | 131022009 | 1/19 |
| **IL-12b** | **G/A** | **749** | **44411954** | **4/15** |
| **H2-Aa** | A/T | 214 | 34283861 | 1/13 |
| | G/C | 236 | 34283883 | 1/14 |
| | C/G | 240 | 34283887 | 2/13 |
| | C/T | 246 | 34283893 | 3/13 |
| | T/C | 264 | 34283905 | 3/11 |
| | T/C | 269 | 34283910 | 3/13 |
| | T/A | 279 | 34283920 | 4/13 |
| | T/C | 305 | 34283946 | 3/13 |
| | T/C | 307 | 34283948 | 3/14 |
| | C/T | 317 | 34283958 | 4/13 |
| | T/C | 327 | 34283968 | 3/13 |
| | G/A | 328 | 34283969 | 4/13 |
| | A/T | 329 | 34283970 | 1/13 |
| | T/C | 330 | 34283971 | 2/13 |
| | C/T | 337 | 34283978 | 3/13 |
| | C/T | 342 | 34283983 | 1/13 |
| | C/T | 349 | 34283990 | 2/13 |
| | T/C | 360 | 34284001 | 4/13 |
| | G/A | 361 | 34284002 | 4/14 |
| | C/T | 379 | 34284020 | 7/13 |
| | G/A | 380 | 34284021 | 2/13 |
| | C/T | 438 | 34284079 | 1/13 |
| | C/T | 442 | 34284083 | 2/13 |
| | C/T | 460 | 34284101 | 1/13 |
| | C/T | 463 | 34284104 | 3/13 |
| | G/A | 480 | 34284121 | 5/13 |
| **H2-Ab** | A/C | 246 | 34267626 | 10/22 |
| | G/A | 274 | 34267654 | 1/22 |
| | G/C | 282 | 34267662 | 5/22 |
| | T/A | 289 | 34267669 | 5/22 |
| | T/C | 291 | 34267671 | 5/22 |
| | T/C | 327 | 34267707 | 21/22 |
| | C/G | 340 | 34267720 | 4/22 |
| | A/G | 367 | 34267747 | 18/22 |
| | C/T | 381 | 34267761 | 2/22 |
| | T/C | 388 | 34267768 | 10/22 |
| | G/A | 405 | 34267785 | 10/22 |
| | A/G | 438 | 34267818 | 17/22 |
| | T/C | 449 | 34267829 | 7/22 |
| | G/A | 466 | 34267846 | 7/22 |
| | C/T | 473 | 34267853 | 18/22 |
| | A/T | 478 | 34267858 | 1/22 |

| | | | | |
|---|---|---|---|---|
| | G/A | 489 | 34267869 | 5/22 |
| | C/T | 517 | 34267897 | 8/22 |
| | C/T | 527 | 34267907 | 2/22 |
| **Myo1a** | C/T | 72 | 127710143 | 5/21 |
| | C/T | 254 | 127710325 | 16/20 |
| | C/T | 467 | 127710538 | 2/19 |
| | T/A | 504 | 127710575 | 2/19 |
| | **T/G** | **546** | **127710617** | **12/19** |
| | C/G | 551 | 127710622 | 1/19 |
| | C/T | 614 | 127710663 | 2/19 |
| | C/T | 636 | 127710685 | 2/19 |
| | C/T | 726 | 127710775 | 13/19 |
| | T/C | 775 | 127711824 | 14/16 |

Supplementary 6. Correlations of the 36 cytokine measures with the PCA axes 1 and 2 (PC1 and PC2, respectively), together accounting for 53% of the total variation. For each cytokine the shading shows, for all stimulants, whether the correlation is greater for PC1 or PC2.

| Cytokine | Stimulant | PC1 | PC2 |
|---|---|---|---|
| IL-1B | LPS | -0.70 | 0.30 |
| | CD3CD28 | -0.57 | 0.49 |
| | CPG | -0.82 | -0.14 |
| | PG | -0.71 | 0.08 |
| IL-6 | LPS | -0.52 | 0.72 |
| | CD3CD28 | -0.34 | 0.81 |
| | CPG | -0.36 | 0.75 |
| | PG | -0.49 | 0.65 |
| IL-10 | LPS | -0.60 | 0.36 |
| | CD3CD28 | -0.61 | 0.13 |
| | CPG | -0.50 | 0.47 |
| | PG | -0.50 | -0.04 |
| IL-12p40 | LPS | -0.28 | 0.10 |
| | CD3CD28 | -0.07 | 0.09 |
| | CPG | -0.44 | 0.41 |
| | PG | -0.63 | -0.36 |
| IL-12p70 | LPS | -0.77 | -0.45 |
| | CD3CD28 | -0.80 | -0.41 |
| | CPG | -0.87 | -0.35 |
| | PG | -0.78 | -0.46 |
| IL-13 | LPS | -0.81 | -0.43 |
| | CD3CD28 | -0.62 | -0.18 |
| | CPG | -0.82 | -0.45 |
| | PG | -0.83 | -0.41 |
| IFNγ | LPS | -0.26 | 0.30 |
| | CD3CD28 | -0.30 | 0.48 |
| | CPG | -0.32 | 0.60 |
| | PG | -0.22 | 0.31 |
| IL-4 | LPS | -0.73 | -0.41 |
| | CD3CD28 | -0.23 | 0.11 |
| | CPG | -0.78 | -0.41 |
| | PG | -0.71 | -0.36 |
| MIP-2a | LPS | -0.52 | 0.56 |
| | CD3CD28 | -0.25 | 0.64 |
| | CPG | -0.22 | 0.39 |
| | PG | -0.35 | 0.54 |

Supplementary 7. The minor allele frequencies for the 14 immune-related SNPs. Loci with overall minor allele frequencies greater than 0.1 are shaded. Myo1a_1 and Myo1a_2 are the control SNPs.

| | SNPs | | | | | | | | | | | | | | | |
|---|---|---|---|---|---|---|---|---|---|---|---|---|---|---|---|---|
| **Site** | **IL-1a** | **IL-1b_U** | **IL-6** | **IL-10** | **IL-13** | **IL-17a_N** | **IL-17a_U** | **IL-17F** | **TNF** | **TILR-5** | **TLR-9** | **H2-Aa** | **H2-Ab** | **H2-Eb** | **Myo1a_2** | **Myo1a_1** |
| **BM** | 0.23 | 0.06 | 0.26 | 0.03 | 0.24 | 0.00 | 0.01 | 0.27 | 0.03 | 0.29 | 0.00 | 0.03 | 0.30 | 0.00 | 0.44 | 0.00 |
| **GL** | 0.08 | 0.00 | 0.00 | 0.20 | 0.00 | 0.00 | 0.40 | 0.50 | 0.00 | 0.00 | 0.00 | 0.13 | 0.02 | 0.03 | 0.17 | 0.00 |
| **HW** | 0.47 | 0.47 | 0.00 | 0.00 | 0.00 | 0.00 | 0.00 | 0.01 | 0.00 | 0.00 | 0.00 | 0.00 | 0.40 | 0.40 | 0.13 | 0.00 |
| **JB** | 0.41 | 0.01 | 0.00 | 0.00 | 0.00 | 0.01 | 0.03 | 0.39 | 0.00 | 0.00 | 0.23 | 0.00 | 0.04 | 0.04 | 0.04 | 0.00 |
| **LU** | 0.03 | 0.03 | 0.00 | 0.00 | 0.07 | 0.00 | 0.14 | 0.17 | 0.46 | 0.00 | 0.00 | 0.00 | 0.07 | 0.00 | 0.03 | 0.00 |
| **PF** | 0.00 | 0.00 | 0.19 | 0.13 | 0.00 | 0.00 | 0.00 | 0.00 | 0.44 | 0.13 | 0.00 | 0.00 | 0.44 | 0.00 | 0.32 | 0.00 |
| **PH** | 0.00 | 0.13 | 0.00 | 0.00 | 0.00 | 0.00 | 0.41 | 0.16 | 0.00 | 0.00 | 0.00 | 0.28 | 0.23 | 0.04 | 0.44 | 0.00 |
| **SK** | 0.02 | 0.02 | 0.00 | 0.04 | 0.00 | 0.13 | 0.04 | 0.13 | 0.00 | 0.00 | 0.00 | 0.00 | 0.00 | 0.00 | 0.00 | 0.48 |
| **SP** | 0.43 | 0.00 | 0.00 | 0.00 | 0.00 | 0.00 | 0.00 | 0.00 | 0.00 | 0.00 | 0.00 | 0.00 | 0.00 | 0.00 | 0.00 | 0.00 |
| **ST** | 0.50 | 0.25 | 0.00 | 0.38 | 0.17 | 0.00 | 0.00 | 0.17 | 0.00 | 0.17 | 0.00 | 0.17 | 0.33 | 0.27 | 0.42 | 0.00 |
| **WF** | 0.15 | 0.00 | 0.19 | 0.00 | 0.00 | 0.00 | 0.00 | 0.42 | 0.00 | 0.00 | 0.00 | 0.27 | 0.33 | 0.00 | 0.17 | 0.00 |
| **WT** | 0.25 | 0.00 | 0.38 | 0.00 | 0.00 | 0.00 | 0.00 | 0.00 | 0.44 | 0.00 | 0.00 | 0.00 | 0.06 | 0.00 | 0.00 | 0.00 |
| **Total** | **0.47** | **0.39** | **0.07** | **0.09** | **0.03** | **0.06** | **0.45** | **0.34** | **0.04** | **0.03** | **0.02** | **0.11** | **0.44** | **0.20** | **0.33** | **0.03** |

Supplementary 8. Heterozygosity of the 14 immune-related SNPs. The observed (O) expected (E) heterozygosity for each SNP at each sampling site, and for all sample sites combined (All) is shown. Values in bold show where the observed and expected values differ by 0.05 or more. Myo1a_1 and Myo1a_2 are the control SNPs.

| Site | | SNPs | | | | | | | | | | | | | | | |
|---|---|---|---|---|---|---|---|---|---|---|---|---|---|---|---|---|---|
| | | **IL-1a** | **IL-1b_U** | **IL-6** | **IL-10** | **IL-13** | **IL-17a_N** | **IL-17a_U** | **IL-17f** | **TNF** | **TLR-5** | **TLR-9** | **H2-Aa** | **H2-Ab** | **H2-Eb** | **Myo1a_1** | **Myo1a_2** |
| **BM** | O | **0.35** | 0.11 | **0.38** | 0.06 | 0.37 | 0.00 | 0.03 | 0.40 | 0.06 | **0.41** | 0.00 | 0.06 | 0.42 | 0.00 | 0.00 | 0.49 |
| | E | **0.46** | 0.11 | **0.29** | 0.06 | 0.37 | 0.00 | 0.03 | 0.37 | 0.06 | **0.29** | 0.00 | 0.06 | 0.42 | 0.00 | 0.00 | 0.52 |
| **GL** | O | 0.15 | 0.00 | 0.00 | **0.32** | 0.00 | 0.00 | **0.48** | **0.50** | 0.00 | 0.00 | 0.00 | 0.23 | 0.03 | 0.06 | 0.00 | 0.28 |
| | E | 0.17 | 0.00 | 0.00 | **0.27** | 0.00 | 0.00 | **0.72** | **0.67** | 0.00 | 0.00 | 0.00 | 0.20 | 0.03 | 0.07 | 0.00 | 0.27 |
| **HW** | O | 0.50 | 0.50 | 0.00 | 0.00 | 0.00 | 0.00 | 0.01 | 0.02 | 0.00 | 0.00 | 0.00 | 0.00 | 0.48 | 0.48 | 0.00 | 0.22 |
| | E | 0.52 | 0.51 | 0.00 | 0.00 | 0.00 | 0.00 | 0.01 | 0.01 | 0.00 | 0.00 | 0.00 | 0.00 | 0.45 | 0.46 | 0.00 | 0.26 |
| **JB** | O | **0.49** | 0.03 | 0.00 | 0.00 | 0.00 | 0.03 | 0.06 | 0.47 | 0.00 | 0.00 | **0.35** | 0.00 | **0.08** | **0.08** | 0.00 | 0.08 |
| | E | **0.43** | 0.03 | 0.00 | 0.00 | 0.00 | 0.03 | 0.00 | 0.49 | 0.00 | 0.00 | **0.46** | 0.00 | **0.03** | **0.03** | 0.00 | 0.09 |
| **LU** | O | 0.06 | 0.06 | 0.00 | 0.00 | 0.12 | 0.00 | **0.24** | **0.28** | **0.50** | 0.00 | 0.00 | 0.00 | 0.12 | 0.00 | 0.00 | 0.06 |
| | E | 0.07 | 0.07 | 0.00 | 0.00 | 0.13 | 0.00 | **0.29** | **0.33** | **0.64** | 0.00 | 0.00 | 0.00 | 0.13 | 0.00 | 0.00 | 0.07 |
| **PF** | O | 0.00 | 0.00 | 0.30 | 0.22 | 0.00 | 0.00 | 0.00 | 0.00 | **0.49** | 0.22 | 0.00 | 0.00 | **0.49** | 0.00 | 0.00 | **0.43** |
| | E | 0.00 | 0.00 | 0.38 | 0.25 | 0.00 | 0.00 | 0.00 | 0.00 | **0.88** | 0.25 | 0.00 | 0.00 | **0.88** | 0.00 | 0.00 | **0.13** |
| **PH** | O | 0.00 | 0.23 | 0.00 | 0.00 | 0.00 | 0.00 | **0.48** | 0.26 | 0.00 | 0.00 | 0.00 | **0.41** | **0.35** | 0.08 | 0.00 | **0.49** |
| | E | 0.00 | 0.26 | 0.00 | 0.00 | 0.00 | 0.00 | **0.41** | 0.25 | 0.00 | 0.00 | 0.00 | **0.34** | **0.42** | 0.08 | 0.00 | **0.43** |
| **SK** | O | 0.04 | 0.04 | 0.00 | **0.07** | 0.00 | **0.23** | **0.07** | **0.23** | 0.00 | 0.00 | 0.00 | 0.00 | 0.00 | 0.00 | 0.50 | 0.00 |
| | E | 0.04 | 0.04 | 0.00 | **0.00** | 0.00 | **0.11** | **0.00** | **0.11** | 0.00 | 0.00 | 0.00 | 0.00 | 0.00 | 0.00 | 0.52 | 0.00 |
| **SF** | O | **0.49** | 0.00 | 0.00 | 0.00 | 0.00 | 0.00 | 0.00 | 0.00 | 0.00 | 0.00 | 0.00 | 0.00 | 0.00 | 0.00 | 0.00 | 0.00 |
| | E | **0.57** | 0.00 | 0.00 | 0.00 | 0.00 | 0.00 | 0.00 | 0.00 | 0.00 | 0.00 | 0.00 | 0.00 | 0.00 | 0.00 | 0.00 | 0.00 |
| **ST** | O | **0.50** | **0.38** | 0.00 | **0.47** | **0.28** | 0.00 | 0.00 | **0.28** | 0.00 | **0.28** | 0.00 | **0.28** | **0.44** | **0.40** | 0.00 | **0.49** |
| | E | **0.67** | **0.50** | 0.00 | **0.42** | **0.33** | 0.00 | 0.00 | **0.17** | 0.00 | **0.33** | 0.00 | **0.33** | **0.50** | **0.18** | 0.00 | **0.33** |
| **WF** | O | **0.25** | 0.00 | **0.31** | 0.00 | 0.00 | 0.00 | 0.00 | 0.49 | 0.00 | 0.00 | 0.00 | **0.39** | 0.44 | 0.00 | 0.00 | 0.28 |
| | E | **0.18** | 0.00 | **0.39** | 0.00 | 0.00 | 0.00 | 0.00 | 0.50 | 0.00 | 0.00 | 0.00 | **0.53** | 0.40 | 0.00 | 0.00 | 0.22 |
| **WT** | O | **0.38** | 0.00 | **0.47** | 0.00 | 0.00 | 0.00 | 0.00 | 0.00 | **0.49** | 0.00 | 0.00 | 0.00 | 0.12 | 0.00 | 0.00 | 0.00 |
| | E | **0.50** | 0.00 | **0.75** | 0.00 | 0.00 | 0.00 | 0.00 | 0.00 | **0.38** | 0.00 | 0.00 | 0.00 | 0.13 | 0.00 | 0.00 | 0.00 |
| **All** | O | **0.34** | **0.28** | **0.06** | 0.04 | 0.04 | **0.01** | **0.12** | **0.20** | 0.05 | 0.04 | 0.04 | **0.09** | **0.33** | **0.21** | 0.03 | **0.26** |
| | E | **0.49** | **0.47** | **0.12** | 0.08 | 0.05 | **0.10** | **0.49** | **0.44** | 0.07 | 0.05 | 0.03 | **0.20** | **0.49** | **0.32** | 0.05 | **0.44** |

Supplementary 9. The number of SNP genotypes at different sample sites. We genotyped 435 mice for 23 SNPs in 21 immune-related loci; 14 of these 23 SNPs were found to be polymorphic. We successfully genotyped 378 (87%) mice for all 14 SNPs; the remaining 57 (13%) mice were successfully genotyped for at least 11 SNPs.

| Locus | Genotype | Total | BM | GL | HW | JB | LU | PH | SK | ST+SP+PH | WF+WT |
|---|---|---|---|---|---|---|---|---|---|---|---|
| H2-Aa | A:A | **29** | 0 | 23 | 0 | 0 | 0 | 6 | 0 | 0 | 0 |
| | A:G | **38** | 2 | 6 | 0 | 0 | 0 | 18 | 0 | 4 | 8 |
| | G:G | **355** | 33 | 1 | 173 | 35 | 17 | 29 | 27 | 25 | 15 |
| H2-Ab | A:A | **114** | 16 | 0 | 29 | 33 | 15 | 1 | 0 | 6 | 14 |
| | A:G | **139** | 14 | 1 | 76 | 1 | 2 | 24 | 0 | 14 | 7 |
| | G:G | **163** | 3 | 29 | 63 | 1 | 0 | 32 | 25 | 8 | 2 |
| H2-Eb | A:A | **43** | 0 | 0 | 30 | 1 | 0 | 0 | 0 | 12 | 0 |
| | A:G | **89** | 0 | 2 | 79 | 1 | 0 | 5 | 0 | 2 | 0 |
| | G:G | **298** | 35 | 28 | 63 | 33 | 17 | 57 | 26 | 14 | 25 |
| IL-1a | C:C | **155** | 0 | 0 | 37 | 13 | 0 | 62 | 26 | 4 | 13 |
| | T:C | **146** | 16 | 5 | 90 | 15 | 1 | 0 | 1 | 12 | 6 |
| | T:T | **131** | 19 | 25 | 47 | 7 | 16 | 0 | 0 | 13 | 4 |
| IL-1b_U | G:G | **105** | 0 | 0 | 37 | 0 | 0 | 42 | 26 | 0 | 0 |
| | G:T | **117** | 4 | 0 | 89 | 1 | 1 | 15 | 1 | 6 | 0 |
| | T:T | **206** | 31 | 30 | 47 | 34 | 16 | 0 | 0 | 23 | 25 |
| IL-6 | C:C | **389** | 21 | 30 | 173 | 34 | 17 | 62 | 26 | 24 | 2 |
| | C:T | **28** | 10 | 0 | 0 | 0 | 0 | 0 | 0 | 5 | 13 |
| | T:T | **15** | 4 | 0 | 0 | 0 | 0 | 0 | 0 | 0 | 11 |
| IL-10 | C:C | **31** | 0 | 2 | 0 | 0 | 0 | 0 | 26 | 3 | 0 |
| | C:T | **17** | 2 | 8 | 0 | 0 | 0 | 0 | 0 | 7 | 0 |
| | T:T | **386** | 33 | 20 | 174 | 35 | 17 | 61 | 1 | 19 | 26 |
| IL-13 | C:C | **411** | 20 | 30 | 174 | 35 | 15 | 61 | 25 | 25 | 26 |
| | C:T | **19** | 13 | 0 | 0 | 0 | 2 | 0 | 0 | 4 | 0 |
| | T:T | **2** | 2 | 0 | 0 | 0 | 0 | 0 | 0 | 0 | 0 |
| IL-17a_N | A:A | **22** | 0 | 0 | 0 | 0 | 0 | 0 | 22 | 0 | 0 |
| | A:G | **4** | 0 | 0 | 0 | 1 | 0 | 0 | 3 | 0 | 0 |
| | G:G | **409** | 35 | 30 | 174 | 34 | 17 | 62 | 2 | 29 | 26 |
| IL-17a_U | A:A | **165** | 33 | 7 | 0 | 32 | 10 | 12 | 26 | 22 | 23 |
| | A:G | **53** | 1 | 21 | 1 | 0 | 6 | 24 | 0 | 0 | 0 |
| | G:G | **204** | 0 | 1 | 171 | 1 | 0 | 23 | 1 | 7 | 0 |
| IL-17F | C:C | **241** | 3 | 5 | 170 | 5 | 0 | 44 | 2 | 1 | 11 |
| | C:T | **87** | 13 | 20 | 1 | 17 | 7 | 15 | 3 | 2 | 9 |
| | T:T | **103** | 19 | 5 | 1 | 13 | 10 | 2 | 22 | 25 | 6 |
| TNF | G:G | **403** | 33 | 30 | 174 | 35 | 2 | 62 | 27 | 19 | 21 |
| | T:G | **24** | 2 | 0 | 0 | 0 | 11 | 0 | 0 | 8 | 3 |
| | T:T | **7** | 0 | 0 | 0 | 0 | 3 | 0 | 0 | 2 | 2 |
| TLR-5 | A:A | **6** | 5 | 0 | 0 | 0 | 0 | 0 | 0 | 1 | 0 |
| | A:G | **16** | 10 | 0 | 0 | 0 | 0 | 0 | 0 | 6 | 0 |
| | G:G | **413** | 20 | 30 | 174 | 35 | 17 | 62 | 27 | 22 | 26 |
| TLR-9 | C:C | **418** | 35 | 30 | 173 | 19 | 17 | 62 | 27 | 29 | 26 |
| | T:C | **16** | 0 | 0 | 0 | 16 | 0 | 0 | 0 | 0 | 0 |
| | T:T | **0** | 0 | 0 | 0 | 0 | 0 | 0 | 0 | 0 | 0 |

Supplementary 10. Selection of the best explanatory models for immune and infection phenotype. This shows the best-supported models for each phenotypic trait, specifically the model with the smallest AIC, and models with similar explanatory power, *i.e*. with a ΔAIC ≤ 2.0. This table therefore does not present the full range of the models tested for each phenotypic trait; see the Methods section for full details on model selection. df is degrees of freedom and is the number of terms in the model. The model with the lowest AIC is shown in italics. The model with very similar explanatory power but which included fewer terms, and so that which was selected, is shown in bold. AIC of the null model is given for comparison.

##### Step 1: selection of the base model for all phenotypic traits.

| **PHENOTYPE** (response variable) | **Models in ΔAIC < 2 and null model** | **df** | **AIC** | **ΔAIC** |
|---|---|---|---|---|
| NK cells | ***site x age (linear) + SMI*** | ***20*** | ***419.75*** | ***0.00*** |
| | site x age (linear) + sex + SMI | 21 | 420.23 | 0.49 |
| | null model | 2 | 610.63 | 190.89 |
| B cells | *age (classes) x sex + site + SMI* | *16* | *444.65* | *0.00* |
| | site x sex + age (linear) + SMI | 21 | 445.19 | 0.54 |
| | **age (linear) x sex + site + SMI** | **14** | **446.49** | **1.84** |
| | null model | 2 | 657.83 | 213.18 |
| Dendritic cells | ***site x age (linear) + SMI*** | ***20*** | ***498.39*** | ***0.00*** |
| | site x age (linear) + sex + SMI | 21 | 498.99 | 0.59 |
| | null model | 2 | 651.64 | 153.24 |
| CD8+ T cells | ***age (linear) + SMI*** | ***4*** | ***264.38*** | ***0.00*** |
| | age (linear) x sex + SMI | 6 | 266.03 | 1.64 |
| | sex + age (linear) + SMI | 5 | 266.30 | 1.91 |
| | null model | 2 | 426.18 | 161.79 |
| CD4+ T cells | ***site + age (linear) + SMI*** | ***12*** | ***203.18*** | ***0.00*** |
| | site + sex + age (linear) + SMI | 13 | 204.01 | 0.83 |
| | null model | 2 | 379.94 | 176.76 |
| Tregs cells | ***site x age (linear) + SMI*** | ***20*** | ***265.35*** | ***0.00*** |
| | site x age (linear) + sex + SMI | 21 | 267.18 | 1.83 |
| | null model | 2 | 454.14 | 188.79 |
| Macrophages | *sex + age (linear) + SMI* | *5* | *553.09* | *0.00* |
| | **age (linear) + SMI** | **4** | **553.50** | **0.41** |
| | age (linear) x sex + SMI | 6 | 554.90 | 1.81 |
| | null model | 2 | 692.78 | 139.68 |
| Monocytes | ***site x age (linear) + SMI*** | ***20*** | ***682.35*** | ***0.00*** |
| | site x age (linear) + sex + SMI | 21 | 683.43 | 1.08 |
| | null model | 2 | 844.04 | 161.69 |
| Hyper-granulocytic myeloid cell | ***site x age (linear) + sex + SMI*** | ***21*** | ***750.60*** | ***0.00*** |
| | null model | 2 | 941.53 | 190.92 |
| Polymorphonuclear cells | ***site + age (classes) + SMI*** | ***13*** | ***721.39*** | ***0.00*** |
| | site + sex + age (classes) + SMI | 14 | 722.36 | 0.97 |
| | null model | 2 | 878.16 | 156.77 |
| Neutrophils | ***site x age (linear) + SMI*** | ***20*** | ***341.23*** | ***0.00*** |
| | site x age (linear) + sex + SMI | 21 | 343.11 | 1.88 |
| | null model | 2 | 471.90 | 130.67 |
| Myeloid Derived Suppressor cells | ***site x age (linear) + SMI*** | ***20*** | ***359.52*** | ***0.00*** |
| | site x age (linear) + sex + SMI | 21 | 361.47 | 1.95 |
| | null model | 2 | 487.95 | 128.42 |
| IgG | ***site + age (classes)*** | ***12*** | ***367.54*** | ***0.00*** |
| | site + age (classes) + SMI | 13 | 368.32 | 0.77 |
| | site + sex + age (classes) | 13 | 369.47 | 1.92 |
| | null model | 2 | 465.80 | 98.25 |
| IgA | ***site x sex + age (classes)*** | ***21*** | ***456.84*** | ***0.00*** |
| | site x sex + age (classes) + SMI | 22 | 458.75 | 1.90 |
| | null model | 2 | 499.73 | 42.88 |
| IgE | *site + sex + age (classes) + SMI* | *14* | *551.61* | *0.00* |
| | **site + sex + age (classes)** | **13** | **552.29** | **0.67** |
| | null model | 2 | 768.98 | 217.36 |
| PC1 cytokines | ***site x sex*** | ***18*** | ***907.35*** | ***0.00*** |
| | site x sex + age (linear) | 19 | 908.94 | 1.59 |
| | site x sex + SMI | 19 | 909.16 | 1.81 |
| | null model | 2 | 921.86 | 14.51 |
| PC2 cytokines | *site x sex + age (linear)* | *19* | *805.80* | *0.00* |
| | site x sex + age (linear) + SMI | 20 | 806.52 | 0.71 |
| | site x sex + SMI | 19 | 806.70 | 0.89 |
| | **site x sex** | **18** | **807.06** | **1.26** |
| | null model | 2 | 824.72 | 18.92 |
| Mite number | ***site + age (classes) + SMI*** | ***19*** | ***1,431.33*** | ***0.00*** |
| | site + sex + age (classes) + SMI | 20 | 1,433.10 | 1.77 |
| | null model | 8 | 1,590.49 | 159.15 |
| Worm number | *age (linear) x sex + site + SMI* | *14* | *842.65* | *0.00* |
| | **site + age (linear) + SMI** | **12** | **842.95** | **0.30** |

| | | | | |
|---|---|---|---|---|
| | site + sex + age (linear) + SMI | 13 | 843.71 | 1.06 |
| | null model | 2 | 1,042.54 | 199.88 |
| Norovirus | ***site + age (classes)*** | ***13*** | ***656.51*** | ***0.00*** |
| | site + age (classes) + SMI | 14 | 658.45 | 1.94 |
| | site + sex + age (classes) | 14 | 658.51 | 1.99 |
| | null model | 3 | 764.29 | 107.78 |
| Parvovirus | ***site + age (classes)*** | ***14*** | ***1,056.09*** | ***0.00*** |
| | site + age (classes) + SMI | 15 | 1,057.68 | 1.59 |
| | site + sex + age (classes) | 15 | 1,057.86 | 1.77 |
| | null model | 4 | 1,146.53 | 90.44 |
| Minute virus | *site + sex + age (classes)* | *14* | *566.70* | *0.00* |
| | **site + age (classes)** | **13** | **568.21** | **1.51** |
| | site + sex + age (classes) + SMI | 15 | 568.69 | 1.99 |
| | null model | 3 | 639.64 | 72.94 |
| Mouse hepatitis virus | ***site + age (classes)*** | ***13*** | ***589.32*** | ***0.00*** |
| | site + sex + age (classes) | 14 | 589.59 | 0.27 |
| | age (classes) x sex + site | 16 | 590.23 | 0.91 |
| | site + age (classes) + SMI | 14 | 591.32 | 2.00 |
| | null model | 3 | 745.44 | 156.12 |
| Sendai virus | ***site + age (classes)*** | ***13*** | ***656.04*** | ***0.00*** |
| | site + sex + age (classes) | 14 | 656.40 | 0.37 |
| | site + age (classes) + SMI | 14 | 658.04 | 2.00 |
| | null model | 3 | 673.89 | 17.86 |
| Coronavirus | *site + age (classes) + SMI* | *13* | *296.71* | *0.00* |
| | site x sex + age (linear) + SMI | 21 | 296.78 | 0.08 |
| | site x sex + age (linear) | 20 | 296.98 | 0.27 |
| | site + age (classes) | 12 | 297.98 | 1.27 |
| | **site + age (linear)** | **11** | **298.39** | **1.69** |
| | site + sex + age (classes) + SMI | 14 | 298.41 | 1.70 |
| | site + age (linear) + SMI | 12 | 298.45 | 1.74 |
| | null model | 2 | 446.99 | 150.28 |
| *Mycoplasma pulmonis* | *site + age (classes) + SMI* | *14* | *640.22* | *0.00* |
| | **site + age (classes)** | **13** | **640.89** | **0.67** |
| | site + sex + age (classes) + SMI | 15 | 641.31 | 1.09 |
| | site + sex + age (classes) | 14 | 642.05 | 1.84 |
| | null model | 2 | 653.73 | 23.58 |

#### Step 2: Selection of the effect of the SNPs. *(here, the simplest "base model" selected in step 1 is always reported for comparison - even if its AIC is outside the ΔAIC ≤ 2.0).*

| PHENOTYPE | SNP | Models in ΔAIC < 2 | df | AIC | ΔAIC |
|---|---|---|---|---|---|
| NK cells | H2-Aa | *site x age (linear) + SMI + H2-Aa* | *22* | 340.94 | 0.00 |
| | | **site x age (linear) + SMI** | **20** | **341.21** | **0.27** |
| | H2-Ab | ***site x age (linear) + SMI*** | ***20*** | ***348.69*** | ***0.00*** |
| | | site x age (linear) + SMI + H2-Ab | 22 | 350.53 | 1.85 |
| | H2-Eb | ***site x age (linear) + SMI*** | ***20*** | ***358.16*** | ***0.00*** |
| | | site x age (linear) + SMI + H2-Eb | 22 | 358.18 | 0.02 |
| | IL-1a | ***site x age (linear) + SMI*** | ***20*** | ***357.09*** | ***0.00*** |
| | | site x age (linear) + SMI + IL-1a | 22 | 359.69 | 2.60 |
| | IL-1b_U | ***site x age (linear) + SMI*** | ***20*** | ***353.82*** | ***0.00*** |
| | | site x age (linear) + SMI + IL-1b_U | 22 | 357.64 | 3.81 |
| | IL-6 | ***site x age (linear) + SMI + IL-6*** | ***22*** | ***354.05*** | ***0.00*** |
| | | site x age (linear) + SMI | 20 | 359.99 | 5.95 |
| | IL-10 | ***site x age (linear) + SMI + IL-10*** | ***22*** | ***351.77*** | ***0.00*** |
| | | site x age (linear) + SMI | 20 | 360.07 | 8.30 |
| | IL-13 | ***site x age (linear) + SMI + IL-13*** | ***22*** | ***355.09*** | ***0.00*** |
| | | site x age (linear) + SMI | 20 | 358.83 | 3.74 |
| | IL-17a_N | ***site x age (linear) + SMI*** | ***20*** | ***360.07*** | ***0.00*** |
| | | site x age (linear) + SMI + IL-17a_N | 22 | 361.68 | 1.62 |
| | IL-17a_U | ***site x age (linear) + SMI*** | ***20*** | ***355.51*** | ***0.00*** |
| | IL-17_f | ***site x age (linear) + SMI*** | ***20*** | ***358.35*** | ***0.00*** |
| | TNF | ***site x age (linear) + SMI + TNF*** | ***22*** | ***354.52*** | ***0.00*** |
| | | *site x age (linear) + SMI* | 20 | 360.07 | 5.55 |
| | TLR-5 | *site x age (linear) + SMI + TLR-5* | *22* | *358.84* | *0.00* |
| | | **site x age (linear) + SMI** | **20** | **360.07** | **1.23** |
| | TLR-9 | ***site x age (linear) + SMI*** | ***20*** | ***360.05*** | ***0.00*** |
| | | site x age (linear) + SMI + TLR-9 | 21 | 361.94 | 1.89 |
| B cells | H2-Aa | *age (linear) x sex + site + SMI + H2-Aa* | *16* | *381.09* | *0.00* |
| | | **age (linear) x sex + site + SMI** | **14** | **381.98** | **0.89** |
| | H2-Ab | *age (linear) x sex + site + SMI + H2-Ab* | *16* | *378.91* | *0.00* |
| | | **age (linear) x sex + site + SMI** | **14** | **379.24** | **0.33** |
| | H2-Eb | ***age (linear) x sex + site + SMI*** | ***14*** | ***387.18*** | ***0.00*** |
| | IL-1a | ***age (linear) x sex + site + SMI*** | ***14*** | ***382.53*** | ***0.00*** |

| | | | | | |
|---|---|---|---|---|---|
| | IL-1b_U | ***age (linear) x sex + site + SMI*** | ***14*** | ***384.99*** | ***0.00*** |
| | IL-6 | ***age (linear) x sex + site + SMI + IL-6*** | ***16*** | ***370.28*** | ***0.00*** |
| | | age (linear) x sex + site + SMI | 14 | 389.59 | 19.30 |
| | IL-10 | ***age (linear) x sex + site + SMI + IL-10*** | ***16*** | ***377.57*** | ***0.00*** |
| | | age (linear) x sex + site + SMI | 14 | 390.76 | 13.18 |
| | IL-13 | ***age (linear) x sex + site + SMI + IL-13*** | ***16*** | ***385.41*** | ***0.00*** |
| | | age (linear) x sex + site + SMI | 14 | 388.32 | 2.91 |
| | IL-17a_N | ***age (linear) x sex + site + SMI*** | ***14*** | ***390.76*** | ***0.00*** |
| | IL-17a_U | ***age (linear) x sex + site + SMI*** | ***14*** | ***381.91*** | ***0.00*** |
| | IL-17f | ***age (linear) x sex + site + SMI*** | ***14*** | ***389.32*** | ***0.00*** |
| | TNF | ***age (linear) x sex + site + SMI + TNF*** | ***16*** | ***384.37*** | ***0.00*** |
| | | age (linear) x sex + site + SMI | 14 | 390.76 | 6.39 |
| | TLR-5 | ***age (linear) x sex + site + SMI*** | ***14*** | ***390.76*** | ***0.00*** |
| | TLR-9 | ***age (linear) x sex + site + SMI*** | ***14*** | ***390.68*** | ***0.00*** |
| | | age (linear) x sex + site + SMI + TLR-9 | 15 | 392.37 | 1.70 |
| **DC** | H2-Aa | ***site x age (linear) + SMI + H2Aa*** | ***22*** | ***429.30*** | ***0.00*** |
| | | site x age (linear) + SMI | 20 | 434.92 | 5.62 |
| | H2-Ab | ***site x age (linear) + SMI*** | ***20*** | ***438.38*** | ***0.00*** |
| | H2-Eb | ***site x age (linear) + SMI*** | ***20*** | ***443.47*** | ***0.00*** |
| | | site x age (linear) + SMI + H2Eb | 22 | 444.03 | 0.56 |
| | IL-1a | ***site x age (linear) + SMI*** | ***20*** | ***448.09*** | ***0.00*** |
| | | site x age (linear) + SMI + IL1a | 22 | 450.96 | 2.86 |
| | IL-1b_U | ***site x age (linear) + SMI*** | ***20*** | ***444.21*** | ***0.00*** |
| | IL-6 | ***site x age (linear) + SMI + IL6*** | ***22*** | ***442.41*** | ***0.00*** |
| | | site x age (linear) + SMI | 20 | 447.96 | 5.55 |
| | IL-10 | ***site x age (linear) + SMI + IL10*** | ***22*** | ***429.60*** | ***0.00*** |
| | | site x age (linear) + SMI | 20 | 450.55 | 20.95 |
| | IL-13 | ***site x age (linear) + SMI + IL13*** | ***22*** | ***445.41*** | ***0.00*** |
| | | site x age (linear) + SMI | 20 | 448.42 | 3.01 |
| | IL-17a_N | ***site x age (linear) + SMI*** | ***20*** | ***450.55*** | ***0.00*** |
| | IL-17a_U | ***site x age (linear) + SMI*** | ***20*** | ***435.38*** | ***0.00*** |
| | IL-17f | ***site x age (linear) + SMI*** | ***20*** | ***448.54*** | ***0.00*** |
| | | site x age (linear) + SMI + IL17f | 22 | 448.97 | 0.44 |
| | TNF | ***site x age (linear) + SMI*** | ***20*** | ***450.55*** | ***0.00*** |
| | | site x age (linear) + SMI + TNF | 22 | 451.08 | 0.53 |
| | TLR-5 | ***site x age (linear) + SMI*** | ***20*** | ***450.55*** | ***0.00*** |
| | TLR-9 | ***site x age (linear) + SMI*** | ***20*** | ***449.25*** | ***0.00*** |
| | | site x age (linear) + SMI + TLR9 | 21 | 451.16 | 1.91 |
| **Tregs** | H2-Aa | ***site x age (linear) + SMI*** | ***20*** | ***248.08*** | ***0.00*** |
| | H2-Ab | *site x age (linear) + SMI + H2Ab* | *22* | *249.67* | *0.00* |
| | | **site x age (linear) + SMI** | **20** | **250.23** | **0.56** |
| | H2-Eb | *site x age (linear) + SMI + H2Eb* | *22* | *255.57* | *0.00* |
| | | **site x age (linear) + SMI** | **20** | **256.44** | **0.87** |
| | IL-1a | ***site x age (linear) + SMI*** | ***20*** | ***254.97*** | ***0.00*** |
| | | site x age (linear) + SMI + IL1a | 22 | 255.87 | 0.90 |
| | IL-1b_U | ***site x age (linear) + SMI*** | ***20*** | ***249.09*** | ***0.00*** |
| | IL-6 | ***site x age (linear) + SMI + IL6*** | ***22*** | ***251.89*** | ***0.00*** |
| | | site x age (linear) + SMI | 20 | 254.79 | 2.89 |
| | IL-10 | ***site x age (linear) + SMI*** | ***20*** | ***257.75*** | ***0.00*** |
| | | site x age (linear) + SMI + IL10 | 22 | 258.55 | 0.81 |
| | IL-13 | ***site x age (linear) + SMI + IL13*** | ***22*** | ***249.92*** | ***0.00*** |
| | | site x age (linear) + SMI | 20 | 255.82 | 5.89 |
| | IL-17a_N | ***site x age (linear) + SMI*** | ***20*** | ***257.75*** | ***0.00*** |
| | IL-17a_U | ***site x age (linear) + SMI*** | ***20*** | ***253.96*** | ***0.00*** |
| | IL-17f | ***site x age (linear) + SMI*** | ***20*** | ***256.46*** | ***0.00*** |
| | TNF | ***site x age (linear) + SMI*** | ***22*** | ***256.92*** | ***0.00*** |
| | | site x age (linear) + SMI + TNF | 20 | 257.75 | 0.83 |
| | TLR-5 | ***site x age (linear) + SMI*** | ***20*** | ***257.75*** | ***0.00*** |
| | TLR-9 | ***site x age (linear) + SMI*** | ***20*** | ***257.75*** | ***0.00*** |
| | | site x age (linear) + SMI + TLR9 | 21 | 259.59 | 1.85 |
| **CD4+ T cells** | H2-Aa | ***site x H2Aa + age (linear) + SMI*** | ***19*** | ***138.77*** | ***0.00*** |
| | | site + age (linear) + SMI | 12 | 145.49 | 6.72 |
| | H2-Ab | ***site + age (linear) + SMI*** | ***12*** | ***148.12*** | ***0.00*** |
| | | site + age (linear) + SMI + H2Ab | 14 | 148.89 | 0.76 |
| | H2-Eb | ***site x H2Eb + age (linear) + SMI*** | ***18*** | ***146.15*** | ***0.00*** |
| | | site + age (linear) + SMI | 12 | 155.03 | 8.87 |
| | IL-1a | ***site + age (linear) + SMI*** | ***12*** | ***153.93*** | ***0.00*** |
| | IL-1b_U | ***site + age (linear) + SMI*** | ***12*** | ***159.02*** | ***0.00*** |
| | IL-6 | ***site x IL6 + age (linear) + SMI*** | ***17*** | ***143.53*** | ***0.00*** |
| | | site + age (linear) + SMI | 12 | 157.45 | 13.92 |
| | IL-10 | ***site x IL10 + age (linear) + SMI*** | ***18*** | ***136.93*** | ***0.00*** |
| | | site + age (linear) + SMI | 12 | 156.29 | 19.36 |

| | | | | | |
|---|---|---|---|---|---|
| | IL-13 | ***age (linear) x IL13 + site + SMI*** | ***15*** | ***147.39*** | ***0.00*** |
| | | site + age (linear) + SMI | 12 | 155.72 | 8.33 |
| | IL-17a_N | ***site + age (linear) + SMI*** | ***12*** | ***156.29*** | ***0.00*** |
| | | site + age (linear) + SMI + IL17a_N | 14 | 157.51 | 1.22 |
| | IL-17a_U | *site x IL17a_U + age (linear) + SMI* | *21* | *154.62* | *0.00* |
| | | **site + age (linear) + SMI** | **12** | **154.91** | **0.29** |
| | IL-17f | ***site + age (linear) + SMI*** | ***12*** | ***154.68*** | ***0.00*** |
| | | site + age (linear) + SMI + IL17f | 14 | 156.66 | 1.98 |
| | TNF | ***site + age (linear) + SMI + TNF*** | ***14*** | ***150.41*** | ***0.00*** |
| | | site + age (linear) + SMI | 12 | 156.29 | 5.88 |
| | TLR-5 | ***site + age (linear) + SMI*** | ***12*** | ***156.29*** | ***0.00*** |
| | TLR-9 | ***site + age (linear) + SMI*** | ***12*** | ***156.29*** | ***0.00*** |
| | | site + age (linear) + SMI + TLR9 | 13 | 158.12 | 1.83 |
| **CD8+ T cells** | H2-Aa | ***age (linear) + SMI + H2Aa*** | ***6*** | ***224.13*** | ***0.00*** |
| | | age (linear) x H2Aa + SMI | 8 | 224.35 | 0.22 |
| | | age (linear) + SMI | 4 | 226.15 | 2.01 |
| | H2-Ab | ***age (linear) + SMI*** | ***4*** | ***222.55*** | ***0.00*** |
| | H2-Eb | ***age (linear) + SMI*** | ***4*** | ***230.84*** | ***0.00*** |
| | | age (linear) + SMI + H2Eb | 6 | 231.11 | 0.27 |
| | IL-1a | ***age (linear) + SMI*** | ***4*** | ***229.19*** | ***0.00*** |
| | | age (linear) + SMI + IL1a | 6 | 229.22 | 0.03 |
| | | age (linear) x IL1a + SMI | 8 | 231.14 | 1.94 |
| | IL-1b_U | ***age (linear) + SMI*** | ***4*** | ***229.01*** | ***0.00*** |
| | | age (linear) + SMI + IL1b_U | 6 | 230.54 | 1.53 |
| | IL-6 | ***age (linear) + SMI + IL6*** | ***6*** | ***228.52*** | ***0.00*** |
| | | age (linear) + SMI | 4 | 232.34 | 3.82 |
| | IL-10 | ***age (linear) + SMI + IL10*** | ***6*** | ***228.57*** | ***0.00*** |
| | | age (linear) x IL10 + SMI | 8 | 229.61 | 1.04 |
| | | age (linear) + SMI | 4 | 231.79 | 3.23 |
| | IL-13 | ***age (linear) + SMI*** | ***4*** | ***231.93*** | ***0.00*** |
| | | age (linear) + SMI + IL13 | 6 | 233.17 | 1.23 |
| | | age (linear) x IL13 + SMI | 7 | 233.02 | 1.09 |
| | IL-17a_N | ***age (linear) + SMI*** | ***4*** | ***231.79*** | ***0.00*** |
| | | age (linear) + SMI + IL17a_N | 6 | 233.63 | 1.84 |
| | IL-17a_U | age (linear) + SMI + IL17a_U | 6 | 227.31 | 0.00 |
| | | ***age (linear) + SMI*** | ***4*** | ***227.91*** | ***0.60*** |
| | IL-17f | ***age (linear) + SMI*** | ***4*** | ***231.04*** | ***0.00*** |
| | | age (linear) + SMI + IL17f | 6 | 232.26 | 1.22 |
| | TNF | ***age (linear) + SMI*** | ***4*** | ***231.79*** | ***0.00*** |
| | TLR-5 | ***age (linear) + SMI*** | ***4*** | ***231.79*** | ***0.00*** |
| | | age (linear) + SMI + TLR5 | 6 | 232.88 | 1.09 |
| | | age (linear) x TLR5 + SMI | 8 | 233.60 | 1.81 |
| | TLR-9 | ***age (linear) + SMI*** | ***4*** | ***231.79*** | ***0.00*** |
| | | age (linear) x TLR9 + SMI | 6 | 232.90 | 1.11 |
| | | age (linear) + SMI + TLR9 | 5 | 233.75 | 1.95 |
| **Macrophages** | H2-Aa | ***age (linear) + SMI*** | ***4*** | ***493.19*** | ***0.00*** |
| | | age (linear) + SMI + H2Aa | 6 | 494.86 | 1.67 |
| | H2-Ab | ***age (linear) + SMI*** | ***4*** | ***498.40*** | ***0.00*** |
| | H2-Eb | *age (linear) + SMI + H2Eb* | *6* | *506.97* | *0.00* |
| | | **age (linear) + SMI** | **4** | **507.35** | **0.38** |
| | IL-1a | ***age (linear) + SMI*** | ***4*** | ***508.59*** | ***0.00*** |
| | IL-1b_U | ***age (linear) + SMI*** | ***4*** | ***507.72*** | ***0.00*** |
| | IL-6 | *age (linear) + SMI + IL6* | *6* | *511.79* | *0.00* |
| | | **age (linear) + SMI** | **4** | **511.81** | **0.02** |
| | | age (linear) x IL6 + SMI | 8 | 511.98 | 0.20 |
| | IL-10 | ***age (linear) + IL10 + SMI*** | ***6*** | ***505.39*** | ***0.00*** |
| | | age (linear) + SMI | 4 | 513.21 | 7.83 |
| | IL-13 | ***age (linear) x IL13 + SMI*** | ***7*** | ***506.23*** | ***0.00*** |
| | | age (linear) + SMI | 4 | 510.61 | 4.38 |
| | IL-17a_N | ***age (linear) + SMI*** | ***4*** | ***513.21*** | ***0.00*** |
| | IL-17a_U | *age (linear) + SMI + IL17a_U* | *6* | *498.39* | *0.00* |
| | | **age (linear) + SMI** | **4** | **499.08** | **0.68** |
| | IL-17f | ***age (linear) + SMI*** | ***4*** | ***510.83*** | ***0.00*** |
| | TNF | ***age (linear) + SMI*** | ***4*** | ***513.21*** | ***0.00*** |
| | TLR-5 | *age (linear) + SMI + TLR5* | *6* | *512.81* | *0.00* |
| | | **age (linear) + SMI** | **4** | **513.21** | **0.40** |
| | | age (linear) x TLR5 + SMI | 8 | 513.34 | 0.53 |
| | TLR-9 | *age (linear) x TLR9 + SMI* | *6* | *512.06* | *0.00* |
| | | age (linear) + TLR9 + SMI | 5 | 512.45 | 0.39 |
| | | **age (linear) + SMI** | **4** | **512.49** | **0.43** |
| **Monocytes** | H2-Aa | ***site x age (linear) + SMI*** | ***20*** | ***608.78*** | ***0.00*** |
| | | site x age (linear) + SMI + H2Aa | 22 | 609.83 | 1.05 |
| | H2-Ab | ***site x age (linear) + SMI*** | ***20*** | ***615.07*** | ***0.00*** |
| | H2-Eb | ***site x age (linear) + SMI*** | ***20*** | ***626.84*** | ***0.00*** |
| | IL-1a | ***site x age (linear) + SMI*** | ***20*** | ***627.03*** | ***0.00*** |
| | IL-1b_U | ***site x age (linear) + SMI*** | ***20*** | ***621.56*** | ***0.00*** |
| | IL-6 | ***site x age (linear) + SMI*** | ***20*** | ***629.56*** | ***0.00*** |

| | | | | | |
|---|---|---|---|---|---|
| | | site x age (linear) + SMI + IL6 | 22 | 631.38 | 1.82 |
| | IL-10 | ***site x age (linear) + SMI*** | ***20*** | ***631.81*** | ***0.00*** |
| | IL-13 | ***site x age (linear) + SMI*** | ***20*** | ***626.95*** | ***0.00*** |
| | | site x age (linear) + SMI + IL13 | 22 | 627.53 | 0.58 |
| | IL-17a_N | ***site x age (linear) + SMI*** | ***20*** | ***631.81*** | ***0.00*** |
| | IL-17a_U | *site x age (linear) + SMI + IL17a_U* | *22* | *613.08* | *0.00* |
| | | **site x age (linear) + SMI** | **20** | **615.01** | **1.93** |
| | IL-17f | ***site x age (linear) + SMI*** | ***20*** | ***628.98*** | ***0.00*** |
| | TNF | ***site x age (linear) + SMI*** | ***20*** | ***631.81*** | ***0.00*** |
| | TLR-5 | ***site x age (linear) + SMI*** | ***20*** | ***631.81*** | ***0.00*** |
| | TLR-9 | ***site x age (linear) + SMI*** | ***20*** | ***630.79*** | ***0.00*** |
| | | **site x age (linear) + SMI + TLR9** | 21 | 632.73 | 1.94 |
| **HGM** | H2-Aa | ***site x age (linear) + sex + SMI*** | ***21*** | ***666.55*** | ***0.00*** |
| | | site x age (linear) + sex + SMI + H2Aa | 23 | 667.70 | 1.16 |
| | H2-Ab | ***site x age (linear) + sex + SMI*** | ***21*** | ***666.43*** | ***0.00*** |
| | | site x age (linear) + sex + SMI + H2Ab | 23 | 668.07 | 1.64 |
| | H2-Eb | ***site x age (linear) + sex + SMI*** | ***21*** | ***682.63*** | ***0.00*** |
| | IL-1a | ***site x age (linear) + sex + SMI*** | ***21*** | ***686.99*** | ***0.00*** |
| | | site x age (linear) + sex + SMI + IL1a | 23 | 687.19 | 0.20 |
| | IL-1b_U | ***site x age (linear) + sex + SMI*** | ***21*** | ***678.78*** | ***0.00*** |
| | IL-6 | ***site x age (linear) + sex + SMI*** | ***21*** | ***686.43*** | ***0.00*** |
| | | site x age (linear) + sex + SMI + IL6 | 23 | 687.45 | 1.02 |
| | IL-10 | ***site x age (linear) + sex + SMI*** | ***21*** | ***689.76*** | ***0.00*** |
| | IL-13 | ***site x age (linear) + sex + SMI + IL13*** | ***23*** | ***642.46*** | ***0.00*** |
| | | site x age (linear) + sex x IL13 + SMI | 24 | 644.24 | 1.78 |
| | | site x age (linear) + sex + SMI | 21 | 645.94 | 3.48 |
| | IL-17a_N | ***site x age (linear) + sex + SMI*** | ***21*** | ***689.76*** | ***0.00*** |
| | IL-17a_U | ***site x age (linear) + sex + SMI*** | ***21*** | ***666.89*** | ***0.00*** |
| | IL-17f | ***site x age (linear) + sex + SMI*** | ***21*** | ***684.65*** | ***0.00*** |
| | | site x age (linear) + sex + SMI + IL17f | 23 | 686.13 | 1.49 |
| | TNF | ***site x age (linear) + sex + SMI*** | ***21*** | ***689.76*** | ***0.00*** |
| | | site x age (linear) + sex + SMI + TNF | 23 | 691.73 | 1.97 |
| | TLR-5 | ***site x age (linear) + sex + SMI*** | ***21*** | ***689.76*** | ***0.00*** |
| | TLR-9 | ***site x age (linear) + sex + SMI*** | ***21*** | ***688.54*** | ***0.00*** |
| | | site x age (linear) + sex + SMI + TLR9 | 22 | 690.54 | 2.00 |
| **PMN** | H2-Aa | ***site + age (classes) + SMI*** | ***13*** | ***661.97*** | ***0.00*** |
| | H2-Ab | ***site + age (classes) + SMI*** | ***13*** | ***666.89*** | ***0.00*** |
| | | site x H2Ab + age (classes) + SMI | 25 | 667.04 | 0.15 |
| | H2-Eb | ***site + age (classes) + SMI*** | ***13*** | ***677.47*** | ***0.00*** |
| | IL-1a | ***site + age (classes) + SMI*** | ***13*** | ***675.54*** | ***0.00*** |
| | | site + age (classes) x IL1a + SMI | 19 | 677.29 | 1.75 |
| | IL-1b_U | ***site + age (classes) + SMI*** | ***13*** | ***669.41*** | ***0.00*** |
| | | site + age (classes) x IL1b_U + SMI | 19 | 670.92 | 1.50 |
| | IL-6 | ***site + age (classes) + SMI*** | ***13*** | ***679.37*** | ***0.00*** |
| | | site + age (classes) + IL6 + SMI | 15 | 681.26 | 1.89 |
| | IL-10 | ***site + age (classes) + SMI*** | ***13*** | ***683.00*** | ***0.00*** |
| | IL-13 | ***site + age (classes) + SMI*** | ***13*** | ***676.41*** | ***0.00*** |
| | IL-17a_N | ***site + age (classes) + SMI*** | ***13*** | ***683.00*** | ***0.00*** |
| | | site + age (classes) + SMI + IL17a_N | 15 | 683.95 | 0.95 |
| | IL-17a_U | ***site x IL17a_U + age (classes) + SMI*** | ***21*** | ***656.38*** | ***0.00*** |
| | | site + age (classes) + SMI | 13 | 659.09 | 2.71 |
| | IL-17f | ***site x IL17f+ age (classes) + SMI*** | 28 | 669.56 | 0.00 |
| | | site + age (classes) + SMI | 13 | 678.98 | 9.43 |
| | TNF | ***site + age (classes) + SMI*** | ***13*** | ***683.00*** | ***0.00*** |
| | TLR-5 | ***site + age (classes) + SMI*** | ***13*** | ***683.00*** | ***0.00*** |
| | TLR-9 | ***site + age (classes) + SMI*** | ***13*** | ***681.80*** | ***0.00*** |
| | | site + age (classes) + SMI + TLR9 | 14 | 683.54 | 1.74 |
| **Neutrophils** | H2-Aa | *site x age (linear) + SMI + H2Aa* | *22* | *314.29* | *0.00* |
| | | **site x age (linear) + SMI** | **20** | **315.63** | **1.33** |
| | H2-Ab | ***site x age (linear) + SMI*** | ***20*** | ***321.71*** | ***0.00*** |
| | H2-Eb | ***site x age (linear) + SMI*** | ***20*** | ***325.09*** | ***0.00*** |
| | | site x age (linear) + SMI + H2Eb | 22 | 326.93 | 1.84 |
| | IL-1a | ***site x age (linear) + SMI*** | ***20*** | ***317.93*** | ***0.00*** |
| | | site x age (linear) + SMI + IL1a | 22 | 319.56 | 1.63 |
| | IL-1b_U | ***site x age (linear) + SMI*** | ***20*** | ***317.15*** | ***0.00*** |
| | | site x age (linear) + SMI + IL1b_U | 22 | 317.25 | 0.11 |
| | IL-6 | ***site x age (linear) + SMI + IL6*** | ***22*** | ***319.99*** | ***0.00*** |
| | | site x age (linear) + SMI | 20 | 324.04 | 4.05 |
| | IL-10 | ***site x age (linear) + SMI + IL10*** | ***22*** | ***313.55*** | ***0.00*** |
| | | site x age (linear) + SMI | 20 | 326.35 | 12.81 |
| | IL-13 | ***site x age (linear) + SMI*** | ***20*** | ***323.20*** | ***0.00*** |
| | IL-17a_N | ***site x age (linear) + SMI*** | ***20*** | ***326.35*** | ***0.00*** |
| | | site x age (linear) + SMI + IL17a_N | 22 | 328.13 | 1.77 |
| | IL-17a_U | ***site x age (linear) + SMI*** | ***20*** | ***314.61*** | ***0.00*** |
| | | site x age (linear) + SMI + IL17a_U | 22 | 315.02 | 0.41 |
| | IL-17f | ***site x age (linear) + SMI*** | ***20*** | ***323.89*** | ***0.00*** |
| | TNF | ***site x age (linear) + SMI*** | ***20*** | ***326.35*** | ***0.00*** |
| | TLR-5 | ***site x age (linear) + SMI*** | ***20*** | ***326.35*** | ***0.00*** |

| | | | | | |
|---|---|---|---|---|---|
| | TLR-9 | ***site x age (linear) + SMI*** | ***20*** | ***326.11*** | ***0.00*** |
| | | site x age (linear) + SMI + TLR9 | 21 | 328.08 | 1.98 |
| **MDSC** | H2-Aa | ***site x age (linear) + SMI + H2Aa*** | ***22*** | ***294.27*** | ***0.00*** |
| | | site x age (linear) + SMI | 20 | 297.63 | 3.36 |
| | H2-Ab | ***site x age (linear) + SMI*** | ***20*** | ***306.38*** | ***0.00*** |
| | H2-Eb | ***site x age (linear) + SMI*** | ***20*** | ***307.94*** | ***0.00*** |
| | IL-1a | ***site x age (linear) + SMI*** | ***20*** | ***303.98*** | ***0.00*** |
| | IL-1b_U | ***site x age (linear) + SMI*** | ***20*** | ***303.92*** | ***0.00*** |
| | IL-6 | ***site x age (linear) + SMI + IL6*** | ***22*** | ***303.75*** | ***0.00*** |
| | | site x age (linear) + SMI | 20 | 311.39 | 7.64 |
| | IL-10 | ***site x age (linear) + SMI + IL10*** | ***22*** | ***302.59*** | ***0.00*** |
| | | site x age (linear) + SMI | 20 | 312.46 | 9.88 |
| | IL-13 | ***site x age (linear) + SMI*** | ***20*** | ***308.94*** | ***0.00*** |
| | IL-17a_N | ***site x age (linear) + SMI + IL17a_N*** | ***22*** | ***308.18*** | ***0.00*** |
| | | site x age (linear) + SMI | 20 | 312.46 | 4.28 |
| | IL-17a_U | ***site x age (linear) + SMI*** | ***20*** | ***301.34*** | ***0.00*** |
| | | site x age (linear) + SMI + IL17a_U | 22 | 302.36 | 1.02 |
| | IL-17f | ***site x age (linear) + SMI*** | ***20*** | ***311.49*** | ***0.00*** |
| | TNF | ***site x age (linear) + SMI*** | ***20*** | ***312.46*** | ***0.00*** |
| | TLR-5 | ***site x age (linear) + SMI*** | ***20*** | ***312.46*** | ***0.00*** |
| | | site x age (linear) + SMI + TLR5 | 22 | 313.98 | 1.52 |
| | TLR-9 | ***site x age (linear) + SMI*** | ***20*** | ***311.80*** | ***0.00*** |
| | | site x age (linear) + SMI + TLR9 | 21 | 313.63 | 1.84 |

| PHENOTYPE | SNP | Models in ΔAIC < 2 | df | AIC | ΔAIC |
|---|---|---|---|---|---|
| **IgA** | H2-Aa | ***site x sex + age (classes)*** | ***21*** | ***420.88*** | ***0.00*** |
| | H2-Ab | ***site x sex + age (classes)*** | ***21*** | ***436.12*** | ***0.00*** |
| | H2-Eb | ***site x sex + age (classes)*** | ***21*** | ***442.72*** | ***0.00*** |
| | | site x sex + age (classes) + H2Eb | 23 | 442.94 | 0.22 |
| | IL-1a | *site x sex + age (classes) + IL1a* | *23* | *442.52* | *0.00* |
| | | **site x sex + age (classes)** | **21** | **443.12** | **0.59** |
| | IL-1b_U | *site x sex + age (classes) + IL1b_U* | *23* | *439.29* | *0.00* |
| | | **site x sex + age (classes)** | **21** | **440.06** | **0.77** |
| | IL-6 | ***site x sex + age (classes) x IL6*** | ***27*** | ***438.74*** | ***0.00*** |
| | | site x sex + age (classes) | 21 | 441.24 | 2.49 |
| | IL-10 | ***site x sex + age (classes)*** | ***21*** | ***443.91*** | ***0.00*** |
| | IL-13 | ***site x sex + age (classes)*** | ***21*** | ***443.20*** | ***0.00*** |
| | IL-17a_N | ***site x sex + age (classes)*** | ***21*** | ***444.21*** | ***0.00*** |
| | | site x sex + age (classes) + IL17a_N | 23 | 446.04 | 1.83 |
| | IL-17a_U | ***site x sex + age (classes)*** | ***21*** | ***433.99*** | ***0.00*** |
| | | site x sex + age (classes) + IL17a_U | 23 | 435.91 | 1.92 |
| | IL-17_f | ***site x sex + age (classes)*** | ***21*** | ***442.15*** | ***0.00*** |
| | | site x sex + age (classes) + IL17f | 23 | 443.81 | 1.66 |
| | TNF | ***site x sex + age (classes)*** | ***21*** | ***443.91*** | ***0.00*** |
| | | site x sex + age (classes) + TNF | 23 | 445.12 | 1.21 |
| | TLR-5 | ***site x sex + age (classes)*** | ***21*** | ***444.21*** | ***0.00*** |
| | | site x sex + age (classes) + TLR5 | 23 | 446.16 | 1.95 |
| | TLR-9 | ***site x sex + age (classes) x TLR9*** | ***24*** | ***437.05*** | ***0.00*** |
| | | site x sex + age (classes) | 21 | 443.84 | 6.78 |
| **IgG** | H2-Aa | *site + age (classes) x H2Aa* | *18* | *357.23* | *0.00* |
| | | **site + age (classes)** | **12** | **357.39** | **0.16** |
| | | site + age (classes) + H2Aa | 14 | 358.35 | 1.12 |
| | H2-Ab | ***site + age (classes) x H2Ab*** | ***18*** | ***341.79*** | ***0.00*** |
| | | site + age (classes) | 12 | 348.83 | 7.04 |
| | H2-Eb | ***site + age (classes)*** | ***12*** | ***359.27*** | ***0.00*** |
| | | site + age (classes) + H2Eb | 14 | 359.94 | 0.68 |
| | IL-1a | ***site + age (classes)*** | ***12*** | ***358.40*** | ***0.00*** |
| | IL-1b_U | ***site + age (classes)*** | ***12*** | ***358.80*** | ***0.00*** |
| | IL-6 | ***site + age (classes)*** | ***12*** | ***360.85*** | ***0.00*** |
| | | site + age (classes) + IL6 | 14 | 362.06 | 1.21 |
| | IL-10 | ***site + age (classes)*** | ***12*** | ***360.44*** | ***0.00*** |
| | IL-13 | ***site + age (classes)*** | ***12*** | ***361.81*** | ***0.00*** |
| | IL-17a_N | ***site + age (classes)*** | ***12*** | ***362.35*** | ***0.00*** |
| | IL-17a_U | ***site + age (classes)*** | ***12*** | ***319.60*** | ***0.00*** |
| | IL-17_f | ***site + age (classes)*** | ***12*** | ***357.07*** | ***0.00*** |
| | | site + age (classes) + IL17f | 14 | 357.81 | 0.73 |
| | TNF | ***site + age (classes)*** | ***12*** | ***362.35*** | ***0.00*** |
| | TLR-5 | ***site + age (classes)*** | ***12*** | ***362.35*** | ***0.00*** |
| | TLR-9 | *site + age (classes) x TLR9* | *15* | *361.64* | *0.00* |
| | | **site + age (classes)** | **12** | **362.52** | **0.88** |
| **IgE** | H2-Aa | *site + sex + age (classes) + H2Aa* | *15* | *502.83* | *0.00* |

| | | | | | |
|---|---|---|---|---|---|
| | | site + sex x H2Aa + age (classes) | 17 | 504.30 | 1.47 |
| | | **site + sex + age (classes)** | **13** | **504.37** | **1.54** |
| | | site x H2Aa + sex + age (classes) | 20 | 504.51 | 1.68 |
| | H2-Ab | *site x H2Ab + sex + age (classes)* | *26* | *512.52* | *0.00* |
| | | **site + sex + age (classes)** | **13** | **514.33** | **1.82** |
| | H2-Eb | *site x H2Eb + sex + age (classes)* | *21* | *525.96* | *0.00* |
| | | **site + sex + age (classes)** | **13** | **527.35** | **1.39** |
| | IL-1a | ***site x IL1a + sex + age (classes)*** | ***25*** | ***493.46*** | ***0.00*** |
| | | site + sex + age (classes) | 13 | 514.01 | 20.55 |
| | IL-1b_U | ***site + sex + age (classes) + IL1b_U*** | ***15*** | ***500.47*** | ***0.00*** |
| | | site x IL1b_U + sex + age (classes) | 21 | 502.05 | 1.58 |
| | | site + sex + age (classes) | 13 | 510.77 | 10.30 |
| | IL-6 | ***site x IL6 + age (classes) + sex*** | ***18*** | ***525.42*** | ***0.00*** |
| | | site + sex + age (classes) | 13 | 528.20 | 2.78 |
| | IL-10 | ***site + sex + age (classes)*** | ***13*** | ***528.99*** | ***0.00*** |
| | | site + sex + age (classes) + IL10 | 15 | 529.71 | 0.72 |
| | IL-13 | ***site + sex + age (classes)*** | ***13*** | ***526.85*** | ***0.00*** |
| | IL-17a_N | ***site + sex + age (classes)*** | ***13*** | ***529.45*** | ***0.00*** |
| | | site + sex + age (classes) x IL17a_N | 16 | 531.07 | 1.63 |
| | IL-17a_U | ***IL-17a_U x site + age (classes) + sex*** | ***23*** | ***489.20*** | ***0.00*** |
| | | site + sex + age (classes) | 13 | 502.88 | 13.68 |
| | IL-17_f | ***site + sex + age (classes)*** | ***13*** | ***528.64*** | ***0.00*** |
| | TNF | ***site x TNF + sex + age (classes)*** | ***19*** | ***525.59*** | ***0.00*** |
| | | site + sex + age (classes) | 13 | 529.45 | 3.86 |
| | TLR-5 | ***site + sex + age (classes)*** | ***13*** | ***529.45*** | ***0.00*** |
| | TLR-9 | ***site + sex + age (classes)*** | ***13*** | ***517.93*** | ***0.00*** |
| | | site + sex + age (classes) + TLR9 | 14 | 518.59 | 0.66 |
| | | site + sex x TLR9 + age (classes) | 15 | 519.75 | 1.82 |
| **PC1 cytokines** | H2-Aa | ***site x sex*** | ***18*** | ***907.35*** | ***0.00*** |
| | | site x sex + H2Aa | 20 | 908.51 | 1.16 |
| | H2-Ab | ***site x sex*** | ***18*** | ***907.35*** | ***0.00*** |
| | H2-Eb | ***site x sex + H2Eb*** | ***20*** | ***905.12*** | ***0.00*** |
| | | site x sex | 18 | 907.35 | 2.23 |
| | IL-1a | ***site x sex*** | ***18*** | ***907.35*** | ***0.00*** |
| | IL-1b_U | ***site x sex + IL1b_U*** | ***20*** | ***904.33*** | ***0.00*** |
| | | site x sex | 18 | 907.35 | 3.02 |
| | IL-6 | ***site x sex + IL6*** | ***20*** | ***903.36*** | ***0.00*** |
| | | site x sex | 18 | 907.35 | 3.99 |
| | IL-10 | ***site x sex + IL10*** | ***19*** | ***884.07*** | ***0.00*** |
| | | site x sex | 18 | 907.35 | 23.28 |
| | IL-13 | ***site x sex*** | ***18*** | ***907.35*** | ***0.00*** |
| | IL-17a_N | ***site x sex*** | ***18*** | ***907.35*** | ***0.00*** |
| | IL-17a_U | ***site x sex*** | ***18*** | ***907.35*** | ***0.00*** |
| | IL-17_f | ***site x sex*** | ***18*** | ***907.35*** | ***0.00*** |
| | | site x sex + IL17f | 20 | 907.78 | 0.43 |
| | TNF | ***site x sex + TNF*** | ***20*** | ***896.30*** | ***0.00*** |
| | | site x sex | 18 | 907.35 | 11.05 |
| | TLR-5 | ***site x sex*** | ***18*** | ***907.35*** | ***0.00*** |
| | TLR-9 | ***site x sex*** | ***18*** | ***907.35*** | ***0.00*** |
| | | site x sex + TLR9 | 19 | 909.19 | 1.84 |
| **PC2 cytokines** | H2-Aa | ***site x sex*** | ***18*** | ***807.06*** | ***0.00*** |
| | H2-Ab | ***site x sex*** | ***18*** | ***807.06*** | ***0.00*** |
| | H2-Eb | ***site x sex*** | ***18*** | ***807.06*** | ***0.00*** |
| | IL-1a | ***site x sex*** | ***18*** | ***807.06*** | ***0.00*** |
| | IL-1b_U | ***site x sex*** | ***18*** | ***807.06*** | ***0.00*** |
| | IL-6 | ***site x sex*** | ***18*** | ***807.06*** | ***0.00*** |
| | | site x sex + IL6 | 20 | 808.58 | 1.52 |
| | IL-10 | ***site x sex + IL10*** | ***19*** | ***797.37*** | ***0.00*** |
| | | site x sex | 18 | 807.06 | 9.69 |
| | IL-13 | ***site x sex*** | ***18*** | ***807.06*** | ***0.00*** |
| | IL-17a_N | ***site x sex*** | ***18*** | ***807.06*** | ***0.00*** |
| | IL-17a_U | ***site x sex + IL17a_U*** | ***20*** | ***804.80*** | ***0.00*** |
| | | site x sex | 18 | 807.06 | 2.27 |
| | IL-17_f | ***site x sex*** | ***18*** | ***807.06*** | ***0.00*** |
| | TNF | ***site x sex*** | ***18*** | ***807.06*** | ***0.00*** |
| | | site x sex + TNF | 20 | 808.29 | 1.23 |
| | TLR-5 | ***site x sex*** | ***18*** | ***807.06*** | ***0.00*** |
| | TLR-9 | ***site x sex*** | ***18*** | ***807.06*** | ***0.00*** |
| | | site x sex + TLR9 | 19 | 808.40 | 1.33 |

| PHENOTYPE | SNP | Models in ∆AIC < 2 | df | AIC | ∆AIC |
|---|---|---|---|---|---|
| **Worms** | H2-Aa | ***site + age (linear) + SMI*** | ***12*** | ***793.95*** | ***0.00*** |

| | | | | | |
|---|---|---|---|---|---|
| | | site + age (linear) + SMI + H2Aa | 14 | 795.12 | 1.17 |
| | H2-Ab | ***site x H2Ab + age (linear) + SMI*** | ***25*** | ***770.85*** | ***0.00*** |
| | | site + age (linear) + SMI | 12 | 774.77 | 3.92 |
| | H2-Eb | ***site + age (linear) + SMI*** | ***12*** | ***793.39*** | ***0.00*** |
| | IL-1a | ***site + age (linear) + SMI*** | ***12*** | ***792.95*** | ***0.00*** |
| | IL-1b_U | ***site + age (linear) + SMI*** | ***12*** | ***793.16*** | ***0.00*** |
| | IL-6 | *site x IL6 + age (linear) + SMI* | *17* | *790.96* | *0.00* |
| | | **site + age (linear) + SMI + IL6** | **14** | **791.65** | **0.70** |
| | | site + age (linear) + SMI | 12 | 798.82 | 7.87 |
| | IL-10 | ***site + age (linear) + SMI*** | ***12*** | ***803.30*** | ***0.00*** |
| | | site + age (linear) + SMI + IL10 | 14 | 804.15 | 0.85 |
| | IL-13 | ***site + age (linear) + SMI*** | ***12*** | ***800.17*** | ***0.00*** |
| | | site + age (linear) + SMI + IL13 | 14 | 801.27 | 1.10 |
| | IL-17a_N | ***site + age (linear) + SMI*** | ***12*** | ***803.30*** | ***0.00*** |
| | IL-17a_U | ***site + age (linear) + SMI*** | ***12*** | ***781.32*** | ***0.00*** |
| | | site x IL17a_U + age (linear) + SMI | 16 | 781.73 | 0.41 |
| | | site + age (linear) + SMI + IL17a_U | 14 | 781.96 | 0.64 |
| | IL-17_f | ***site + age (linear) + SMI*** | ***12*** | ***796.31*** | ***0.00*** |
| | | site + age (linear) + SMI + IL17f | 14 | 797.41 | 1.10 |
| | | site x IL17f + age (linear) + SMI | 16 | 797.44 | 1.13 |
| | TNF | ***site + age (linear) + SMI*** | ***12*** | ***802.28*** | ***0.00*** |
| | TLR-5 | *site x TLR5 + age (linear) + SMI* | *16* | *801.48* | *0.00* |
| | | **site + age (linear) + SMI** | **12** | **803.30** | **1.83** |
| | TLR-9 | ***site + age (linear) + SMI*** | ***12*** | ***801.61*** | ***0.00*** |
| | | site + age (linear) + SMI + TLR9 | 13 | 803.23 | 1.62 |
| **Mites** | H2-Aa | ***site + age (classes) + SMI + H2Aa*** | ***21*** | ***1320.93*** | ***0.00*** |
| | | site x H2Aa + age (classes) + SMI | 26 | 1321.55 | 0.62 |
| | | site + age (classes) + SMI | 19 | 1323.21 | 2.28 |
| | H2-Ab | ***site + age (classes) + SMI*** | ***19*** | ***1302.12*** | ***0.00*** |
| | H2-Eb | *site + age (classes) + SMI + H2Eb* | *21* | *1345.69* | *0.00* |
| | | site x H2Eb + age (classes) + SMI | 27 | 1346.03 | 0.34 |
| | | **site + age (classes) + SMI** | **19** | **1346.83** | **1.14** |
| | IL-1a | ***site + age (classes) + SMI*** | ***19*** | ***1348.51*** | ***0.00*** |
| | IL-1b_U | ***site + age (classes) + SMI*** | ***19*** | ***1346.72*** | ***0.00*** |
| | IL-6 | ***site + age (classes) + SMI*** | ***19*** | ***1350.86*** | ***0.00*** |
| | IL-10 | ***site + age (classes) + SMI*** | ***19*** | ***1360.02*** | ***0.00*** |
| | | site + age (classes) + SMI + IL10 | 21 | 1361.39 | 1.37 |
| | IL-13 | ***site + age (classes) + SMI*** | ***19*** | ***1354.10*** | ***0.00*** |
| | IL-17a_N | ***site + age (classes) + SMI*** | ***19*** | ***1360.02*** | ***0.00*** |
| | IL-17a_U | ***site + age (classes) + SMI*** | ***19*** | ***1316.97*** | ***0.00*** |
| | IL-17_f | ***site + age (classes) + SMI*** | ***19*** | ***1344.85*** | ***0.00*** |
| | TNF | ***site + age (classes) + SMI*** | ***19*** | ***1358.76*** | ***0.00*** |
| | | site + age (classes) + SMI + TNF | 21 | 1360.50 | 1.74 |
| | TLR-5 | *site x TLR5 + age (classes) + SMI* | *23* | *1356.77* | *0.00* |
| | | **site + age (classes) + SMI + TLR5** | **21** | **1357.60** | **0.82** |
| | | site + age (classes) + SMI | 19 | 1360.02 | 3.25 |
| | TLR-9 | ***site + age (classes) + SMI + TLR9*** | ***20*** | ***1346.77*** | ***0.00*** |
| | | site + age (classes) + SMI | 19 | 1356.73 | 9.96 |
| **Norovirus** | H2-Aa | ***site + age (classes)*** | ***13*** | ***601.58*** | ***0.00*** |
| | | site + age (classes) + H2Aa | 15 | 602.19 | 0.61 |
| | H2-Ab | ***site + age (classes)*** | ***13*** | ***584.63*** | ***0.00*** |
| | | site + age (classes) + H2Ab | 15 | 585.56 | 0.93 |
| | H2-Eb | ***site + age (classes) x H2Eb*** | ***19*** | ***606.32*** | ***0.00*** |
| | | site + age (classes) | 13 | 609.40 | 3.08 |
| | IL-1a | *site + age (classes) + IL1a* | *15* | *609.70* | *0.00* |
| | | **site + age (classes)** | **13** | **610.18** | **0.49** |
| | IL-1b_U | ***site + age (classes)*** | ***13*** | ***608.25*** | ***0.00*** |
| | IL-6 | *site + age (classes) + IL6* | *15* | *612.54* | *0.00* |
| | | **site + age (classes)** | **13** | **613.46** | **0.92** |
| | IL-10 | ***site + age (classes) x IL10*** | ***19*** | ***609.82*** | ***0.00*** |
| | | site + age (classes) | 13 | 616.60 | 6.78 |
| | IL-13 | ***site + age (classes)*** | ***13*** | ***612.43*** | ***0.00*** |
| | IL-17a_N | ***site + age (classes)*** | ***13*** | ***616.60*** | ***0.00*** |
| | IL-17a_U | *site + age (classes) x IL17a_U* | *19* | *597.66* | *0.00* |
| | | **site + age (classes)** | **13** | **598.80** | **1.13** |
| | IL-17_f | *site + age (classes) x IL17f* | *19* | *609.05* | *0.00* |
| | | **site + age (classes)** | **13** | **609.65** | **0.60** |
| | TNF | ***site + age (classes)*** | ***13*** | ***614.71*** | ***0.00*** |
| | TLR-5 | *site + age (classes) x TLR5* | *19* | *615.99* | *0.00* |
| | | **site + age (classes)** | **13** | **616.60** | **0.61** |
| | TLR-9 | ***site + age (classes)*** | ***13*** | ***615.56*** | ***0.00*** |
| | | site + age (classes) + TLR9 | 14 | 615.89 | 0.33 |
| **Parvovirus** | H2-Aa | ***site + age (classes)*** | ***29*** | ***975.10*** | ***0.00*** |
| | H2-Ab | ***site + age (classes)*** | ***29*** | ***949.88*** | ***0.00*** |
| | | site + age (classes) + H2Ab | 31 | 950.78 | 0.90 |
| | H2-Eb | ***site + age (classes) + H2Eb*** | ***31*** | ***974.47*** | ***0.00*** |

| | | | | | |
|---|---|---|---|---|---|
| | | site + age (classes) | 29 | 979.15 | 4.68 |
| | IL-1a | ***site + age (classes)*** | ***29*** | ***978.62*** | ***0.00*** |
| | | site + age (classes) + IL1a | 31 | 980.19 | 1.57 |
| | IL-1b_U | ***site + age (classes)*** | ***29*** | ***976.85*** | ***0.00*** |
| | IL-6 | ***site + age (classes)*** | ***29*** | ***978.66*** | ***0.00*** |
| | IL-10 | ***site + age (classes)*** | ***29*** | ***985.69*** | ***0.00*** |
| | | site + age (classes) + IL10 | 31 | 986.85 | 1.16 |
| | IL-13 | ***site + age (classes)*** | ***29*** | ***981.21*** | ***0.00*** |
| | IL-17a_N | ***site + age (classes)*** | ***29*** | ***985.69*** | ***0.00*** |
| | IL-17a_U | ***site + age (classes)*** | ***29*** | ***957.04*** | ***0.00*** |
| | IL-17_f | ***site + age (classes)*** | ***29*** | ***977.45*** | ***0.00*** |
| | TNF | ***site + age (classes)*** | ***29*** | ***983.63*** | ***0.00*** |
| | TLR-5 | ***site + age (classes)*** | ***29*** | ***985.69*** | ***0.00*** |
| | TLR-9 | ***site + age (classes)*** | ***29*** | ***984.37*** | ***0.00*** |
| | | site + age (classes) + TLR9 | 30 | 986.22 | 1.85 |
| | H2-Aa | ***site + age (classes)*** | ***21*** | ***516.68*** | ***0.00*** |
| | H2-Ab | ***site + age (classes) + H2Ab*** | ***23*** | ***499.97*** | ***0.00*** |
| | | site + age (classes) | 21 | 503.56 | 3.59 |
| | H2-Eb | ***site + age (classes)*** | ***21*** | ***529.67*** | ***0.00*** |
| | | site + age (classes) + H2Eb | 23 | 530.93 | 1.26 |
| | IL-1a | ***site + age (classes) + IL1a*** | ***23*** | ***519.96*** | ***0.00*** |
| | | site + age (classes) | 21 | 522.30 | 2.34 |
| | IL-1b_U | ***site + age (classes)*** | ***21*** | ***523.85*** | ***0.00*** |
| | IL-6 | ***site + age (classes)*** | ***21*** | ***530.89*** | ***0.00*** |
| | | site + age (classes) + IL6 | 23 | 532.83 | 1.94 |
| **Minute virus** | IL-10 | ***site + age (classes)*** | ***21*** | ***531.55*** | ***0.00*** |
| | IL-13 | ***site + age (classes)*** | ***21*** | ***524.86*** | ***0.00*** |
| | IL-17a_N | *site + age (classes) + IL17a_N* | *23* | *531.33* | *0.00* |
| | | **site + age (classes)** | **21** | **531.69** | **0.36** |
| | IL-17a_U | ***site + age (classes)*** | ***21*** | ***515.54*** | ***0.00*** |
| | IL-17_f | ***site + age (classes)*** | ***21*** | ***529.23*** | ***0.00*** |
| | | site + age (classes) + IL17f | 23 | 530.76 | 1.53 |
| | TNF | ***site + age (classes)*** | ***21*** | ***531.43*** | ***0.00*** |
| | TLR-5 | ***site + age (classes)*** | ***21*** | ***530.49*** | ***0.00*** |
| | TLR-9 | ***site + age (classes)*** | ***21*** | ***531.24*** | ***0.00*** |
| | | site + age (classes) + TLR9 | 22 | 532.82 | 1.59 |
| | H2-Aa | ***site + age (classes) + H2Aa*** | ***30*** | ***533.03*** | ***0.00*** |
| | | site + age (classes) | 28 | 537.54 | 4.51 |
| | H2-Ab | ***site + age (classes)*** | ***28*** | ***511.05*** | ***0.00*** |
| | H2-Eb | ***site + age (classes)*** | ***28*** | ***540.87*** | ***0.00*** |
| | IL-1a | ***site + age (classes)*** | ***28*** | ***543.20*** | ***0.00*** |
| | IL-1b_U | ***site + age (classes)*** | ***28*** | ***529.30*** | ***0.00*** |
| | IL-6 | ***site + age (classes) + IL6*** | ***30*** | ***524.80*** | ***0.00*** |
| | | site + age (classes) | 28 | 532.89 | 8.09 |
| | IL-10 | site + age (classes) | 28 | 545.01 | 0.00 |
| | IL-13 | ***site + age (classes)*** | ***28*** | ***538.28*** | ***0.00*** |
| **MHV** | IL-17a_N | ***site + age (classes)*** | ***28*** | ***545.01*** | ***0.00*** |
| | IL-17a_U | ***site + age (classes)*** | ***28*** | ***527.07*** | ***0.00*** |
| | IL-17_f | *site + age (classes) + IL17a_U* | *29* | *543.39* | *0.00* |
| | | **site + age (classes)** | **28** | **543.91** | **0.52** |
| | TNF | ***site + age (classes) + TNF*** | ***30*** | ***539.72*** | ***0.00*** |
| | | site + age (classes) | 28 | 543.55 | 3.83 |
| | TLR-5 | ***site + age (classes)*** | ***28*** | ***545.01*** | ***0.00*** |
| | TLR-9 | *site + age (classes) + TLR9* | *29* | *543.39* | *0.00* |
| | | **site + age (classes)** | **28** | **543.91** | **0.52** |
| | H2-Aa | ***site + age (classes)*** | ***13*** | ***608.79*** | ***0.00*** |
| | | site x H2Aa + age (classes) | 20 | 610.64 | 1.84 |
| | H2-Ab | *site + age (classes) + H2Ab* | *15* | *590.76* | *0.00* |
| | | **site + age (classes)** | **13** | **591.13** | **0.37** |
| | H2-Eb | *site + age (classes) + H2Eb* | *15* | *611.44* | *0.00* |
| | | **site + age (classes)** | **13** | **612.70** | **1.26** |
| | IL-1a | ***site + age (classes)*** | ***13*** | ***614.31*** | ***0.00*** |
| | IL-1b_U | ***site + age (classes)*** | ***13*** | ***613.72*** | ***0.00*** |
| | IL-6 | *site + age (classes) x IL6* | *19* | *612.71* | *0.00* |
| **Sendai virus** | | **site + age (classes) + IL6** | **15** | **614.13** | **1.41** |
| | | site + age (classes) | 13 | 616.17 | 3.46 |
| | IL-10 | ***site + age (classes)*** | ***13*** | ***619.47*** | ***0.00*** |
| | IL-13 | ***site + age (classes)*** | ***13*** | ***614.48*** | ***0.00*** |
| | IL-17a_N | ***site + age (classes)*** | ***13*** | ***619.47*** | ***0.00*** |
| | IL-17a_U | ***site + age (classes)*** | ***13*** | ***601.91*** | ***0.00*** |
| | IL-17_f | ***site + age (classes)*** | ***13*** | ***613.75*** | ***0.00*** |
| | | site + age (classes) + IL17f | 15 | 614.50 | 0.76 |
| | TNF | ***site x TNF + age (classes)*** | ***20*** | ***608.94*** | ***0.00*** |

| | | | | | |
|---|---|---|---|---|---|
| | | site + age (classes) | 13 | 617.46 | 8.52 |
| | TLR-5 | ***site + age (classes)*** | ***13*** | ***619.47*** | ***0.00*** |
| | | site + age (classes) x TLR5 | 19 | 620.13 | 0.65 |
| | | site + age (classes) + TLR5 | 15 | 621.14 | 1.67 |
| | TLR-9 | ***site + age (classes) x TLR9*** | ***16*** | ***614.44*** | ***0.00*** |
| | | site + age (classes) | 13 | 616.80 | 2.36 |
| **Coronavirus** | H2-Aa | ***site + age (linear)*** | ***11*** | ***279.27*** | ***0.00*** |
| | | site + age (linear) + H2Aa | 13 | 279.31 | 0.05 |
| | | site + IL17a_N x age (linear) | 15 | 280.84 | 1.58 |
| | H2-Ab | ***site + age (linear)*** | ***11*** | ***275.74*** | ***0.00*** |
| | H2-Eb | ***site + age (linear)*** | ***11*** | ***270.09*** | ***0.00*** |
| | | site + age (linear) + H2Eb | 13 | 271.45 | 1.36 |
| | IL-1a | *site + age (linear) + IL1a* | *13* | *279.22* | *0.00* |
| | | **site + age (linear)** | **11** | **279.74** | **0.52** |
| | IL-1b_U | ***site + age (linear)*** | ***11*** | ***279.64*** | ***0.00*** |
| | | site + age (linear) + IL1b_U | 13 | 281.25 | 1.61 |
| | IL-6 | ***site + age (linear)*** | ***11*** | ***279.58*** | ***0.00*** |
| | | site + age (linear) + IL6 | 13 | 281.28 | 1.70 |
| | IL-10 | *site x IL10 + age (linear)* | *17* | *279.16* | *0.00* |
| | | **site + age (linear)** | **11** | **279.93** | **0.77** |
| | IL-13 | ***site + age (linear)*** | ***11*** | ***279.38*** | ***0.00*** |
| | | site + age (linear) + IL13 | 13 | 279.38 | 0.00 |
| | IL-17a_N | ***site + IL17a_N x age (linear)*** | ***15*** | ***277.04*** | ***0.00*** |
| | | site + age (linear) | 11 | 279.93 | 2.89 |
| | IL-17a_U | ***site + age (linear)*** | ***11*** | ***274.86*** | ***0.00*** |
| | | site + age (linear) + IL17a_U | 13 | 276.35 | 1.49 |
| | IL-17_f | ***site + age (linear)*** | ***11*** | ***279.10*** | ***0.00*** |
| | | site + age (linear) + IL17f | 13 | 280.54 | 1.44 |
| | | site + IL17a_N x age (linear) | 15 | 280.62 | 1.52 |
| | TNF | ***site + age (linear)*** | ***11*** | ***278.42*** | ***0.00*** |
| | | site + age (linear) + TNF | 13 | 280.04 | 1.62 |
| | TLR-5 | ***site + age (linear)*** | ***11*** | ***279.93*** | ***0.00*** |
| | TLR-9 | ***site + age (linear)*** | ***11*** | ***279.79*** | ***0.00*** |
| | | site + age (linear) + TLR9 | 12 | 280.15 | 0.36 |
| ***M. pulmonis*** | H2-Aa | *site x H2Aa + age (classes)* | *19* | *586.00* | *0.00* |
| | | **site + age (classes)** | **12** | **586.56** | **0.56** |
| | H2-Ab | ***site + age (classes)*** | ***12*** | ***567.20*** | ***0.00*** |
| | H2-Eb | *site + age (classes) + H2Eb* | *14* | *587.91* | *0.00* |
| | | **site + age (classes)** | **12** | **589.62** | **1.71** |
| | IL-1a | ***site + age (classes)*** | ***12*** | ***591.03*** | ***0.00*** |
| | | site + age (classes) + IL1a | 14 | 592.10 | 1.07 |
| | IL-1b_U | ***site + age (classes)*** | ***12*** | ***589.22*** | ***0.00*** |
| | IL-6 | ***site + age (classes) + IL6*** | ***14*** | ***587.82*** | ***0.00*** |
| | | site + age (classes) | 12 | 591.12 | 3.30 |
| | IL-10 | ***site + age (classes)*** | ***12*** | ***595.98*** | ***0.00*** |
| | | site + age (classes) x IL10 | 18 | 597.51 | 1.53 |
| | IL-13 | ***site + age (classes)*** | ***15*** | ***595.06*** | ***0.00*** |
| | IL-17a_N | ***site + age (classes)*** | ***12*** | ***595.98*** | ***0.00*** |
| | | site + age (classes) + IL17a_N | 14 | 596.82 | 0.83 |
| | IL-17a_U | ***site + age (classes)*** | ***12*** | ***586.34*** | ***0.00*** |
| | | site + age (classes) + IL17a_U | 14 | 588.08 | 1.74 |
| | IL-17_f | ***site + age (classes)*** | ***12*** | ***587.47*** | ***0.00*** |
| | | site + age (classes) + IL17f | 14 | 588.32 | 0.85 |
| | | site x IL17f + age (classes) | 28 | 589.20 | 1.73 |
| | TNF | ***site + age (classes)*** | ***12*** | ***596.74*** | ***0.00*** |
| | | site + age (classes) + TNF | 14 | 597.38 | 0.64 |
| | TLR-5 | ***site + age (classes)*** | ***12*** | ***595.98*** | ***0.00*** |
| | | site x TLR5 + age (classes) | 18 | 596.74 | 0.75 |
| | | site + age (classes) x TLR5 | 16 | 597.43 | 1.45 |
| | TLR-9 | ***site + age (classes)*** | ***12*** | ***593.66*** | ***0.00*** |
| | | site x TLR9 + age (classes) | 15 | 594.30 | 0.65 |
| | | site + age (classes) x TLR9 | 13 | 595.64 | 1.98 |

Supplementary 11. Parameter estimates of the selected generalized linear models of immune and infection phenotype in wild mice. Estimates are presented ± standard error (SE). Statistical significance is represented by * for $p < 0.05$, ** for $p < 0.01$ and *** for $p < 0.001$. The potential fixed effects tested were the 9 different sites (here siteST stands for PF+SP+ST and siteW stands for WF+WT) ; age (linear function as "age_l" or three classes as "age_cl" ("IM" for immatures, "YO" for young, "AD" for adults); body condition ("SMI"); sex (males "M" vs females "F") and the loci of interest. Interactions are represented with " x ". The parameter in brackets is the one to which the comparison is made.

#### Antibodies

**IgA ~ site x sex + age (classes) x IL6**

| | Estimate | ±SE | t value | p |
|---|---|---|---|---|
| (Intercept) | 2.25 | 0.15 | 15.46 | *** |
| siteGL | 0.13 | 0.18 | 0.71 | - |
| siteHW | 0.13 | 0.16 | 0.80 | - |
| siteJB | -0.13 | 0.19 | -0.66 | - |
| siteLU | 0.35 | 0.36 | 0.95 | - |
| sitePH | -0.14 | 0.18 | -0.79 | - |
| siteSK | -0.30 | 0.21 | -1.44 | - |
| siteST | -0.29 | 0.21 | -1.42 | - |
| siteW | -0.73 | 0.29 | -2.57 | * |
| sex(M) | 0.00 | 0.18 | -0.01 | - |
| age_cl(IM) | -0.28 | 0.08 | -3.63 | *** |
| age_cl(YO) | 0.09 | 0.08 | 1.04 | - |
| IL6(CT) | -0.15 | 0.23 | -0.66 | - |
| IL6(TT) | -1.52 | 0.55 | -2.75 | ** |
| siteGL x sex(M) | -0.52 | 0.25 | -2.04 | * |
| siteHW x sex(M) | 0.01 | 0.20 | 0.04 | - |
| siteJB x sex(M) | -0.02 | 0.27 | -0.06 | - |
| siteLU x sex(M) | -0.49 | 0.41 | -1.19 | - |
| sitePH x sex(M) | -0.02 | 0.22 | -0.11 | - |
| siteSK x sex(M) | 0.63 | 0.32 | 1.98 | * |
| siteST x sex(M) | 0.22 | 0.26 | 0.82 | - |
| siteW x sex(M) | 0.93 | 0.34 | 2.77 | ** |
| age_cl(IM) x IL6(CT) | 0.44 | 0.27 | 1.64 | - |
| age_cl(YO) x IL6(CT) | -0.07 | 0.27 | -0.26 | - |
| age_cl(IM) x IL6(TT) | 1.74 | 0.73 | 2.38 | * |
| age_cl(YO) x IL6(TT) | 1.33 | 0.62 | 2.17 | * |

**IgA ~ site x sex + age (classes) x TLR9**

| | Estimate | ±SE | t value | p |
|---|---|---|---|---|
| (Intercept) | 2.18 | 0.13 | 16.87 | *** |
| siteGL | 0.16 | 0.17 | 0.95 | - |
| siteHW | 0.14 | 0.15 | 0.93 | - |
| siteJB | 0.00 | 0.19 | -0.02 | - |
| siteLU | 0.41 | 0.36 | 1.15 | - |
| sitePH | -0.14 | 0.17 | -0.81 | - |
| siteSK | -0.27 | 0.20 | -1.38 | - |
| siteST | -0.26 | 0.20 | -1.33 | - |
| siteW | -0.93 | 0.23 | -4.02 | *** |
| sex(M) | -0.03 | 0.17 | -0.15 | - |
| age_cl(IM) | -0.20 | 0.08 | -2.63 | ** |
| age_cl(YO) | 0.11 | 0.08 | 1.37 | - |
| Tlr9(TC) | 0.59 | 0.50 | 1.20 | - |
| siteGL x sex(M) | -0.49 | 0.25 | -1.96 | - |
| siteHW x sex(M) | 0.04 | 0.20 | 0.20 | - |
| siteJB x sex(M) | 0.15 | 0.26 | 0.59 | - |
| siteLU x sex(M) | -0.51 | 0.40 | -1.27 | - |
| sitePH x sex(M) | 0.03 | 0.22 | 0.13 | - |
| siteSK x sex(M) | 0.61 | 0.31 | 1.97 | - |
| siteST x sex(M) | 0.22 | 0.26 | 0.87 | - |
| siteW x sex(M) | 1.12 | 0.31 | 3.58 | *** |
| age_cl(IM) x Tlr9(TC) | -1.43 | 0.54 | -2.64 | ** |
| age_cl(YO) x Tlr9(TC) | -0.76 | 0.54 | -1.40 | - |

**IgG ~ H2Ab x age (classes) + site**

| | Estimate | ±SE | t value | p |
|---|---|---|---|---|
| (Intercept) | 4.20 | 0.09 | 47.41 | *** |
| age_cl(IM) | -0.51 | 0.10 | -5.34 | *** |
| age_cl(YO) | -0.35 | 0.10 | -3.60 | *** |
| H2Ab(AG) | -0.25 | 0.11 | -2.30 | * |
| H2Ab(GG) | -0.35 | 0.11 | -3.31 | ** |
| siteGL | -0.15 | 0.10 | -1.41 | - |
| siteHW | 0.02 | 0.08 | 0.28 | - |
| siteJB | -0.21 | 0.09 | -2.18 | * |
| siteLU | -0.16 | 0.15 | -1.05 | - |
| sitePH | 0.16 | 0.09 | 1.81 | . |
| siteSK | 0.44 | 0.11 | 4.00 | *** |
| siteST | -0.10 | 0.10 | -0.99 | - |
| siteWF+WT | -0.05 | 0.11 | -0.44 | - |
| age_cl(IM) x H2Ab(AG) | 0.16 | 0.13 | 1.21 | - |
| age_cl(YO) x H2Ab(AG) | 0.15 | 0.13 | 1.11 | - |
| age_cl(IM) x H2Ab(GG) | 0.36 | 0.12 | 2.96 | ** |
| age_cl(YO) x H2Ab(GG) | 0.38 | 0.12 | 3.07 | ** |

**IgE ~ IL-1a x site + age (classes) + sex**

| | Estimate | ±SE | t value | p |
|---|---|---|---|---|
| (Intercept) | 3.23 | 0.28 | 11.74 | *** |
| siteGL | -0.37 | 0.13 | -2.80 | ** |
| siteHW | -1.02 | 0.28 | -3.65 | *** |
| siteJB | -1.08 | 0.30 | -3.62 | *** |
| siteLU | -0.52 | 0.19 | -2.79 | ** |
| sitePH | -1.72 | 0.28 | -6.22 | *** |
| siteSK | -0.83 | 0.28 | -2.92 | ** |
| siteST | -1.59 | 0.41 | -3.91 | *** |
| siteWF+WT | -1.07 | 0.24 | -4.50 | *** |
| IL1a(TC) | -1.08 | 0.29 | -3.69 | *** |
| IL1a(TT) | -0.67 | 0.25 | -2.66 | ** |
| age_cl(IM) | -0.62 | -0.06 | 10.76 | *** |
| age_cl(YO) | -0.30 | 0.06 | -5.08 | *** |
| sex(M) | -0.12 | 0.04 | -2.75 | ** |
| siteGL x IL1a(TC) | 0.61 | 0.26 | 2.34 | * |
| siteHW x IL1a(TC) | 1.22 | 0.30 | 4.01 | *** |
| siteJB x IL1a(TC) | 1.22 | 0.34 | 3.63 | *** |
| siteLU x IL1a(TC) | 0.02 | 0.48 | 0.05 | - |
| siteSK x IL1a(TC) | 1.29 | 0.53 | 2.44 | * |
| siteST x IL1a(TC) | 1.85 | 0.44 | 4.22 | *** |
| siteWF+WT x IL1a(TC) | 0.65 | 0.33 | 2.00 | * |
| siteHW x IL1a(TT) | 1.08 | 0.27 | 3.98 | *** |
| siteJB x IL1a(TT) | 0.71 | 0.32 | 2.19 | * |
| siteST x IL1a(TT) | 1.55 | 0.41 | 3.77 | *** |

**IgE ~ IL-17a_U x site + age (classes) + sex**

| | Estimate | ±SE | t value | p |
|---|---|---|---|---|
| (Intercept) | 2.38 | 0.09 | 27.72 | *** |
| siteGL | -0.37 | 0.18 | -2.03 | * |
| siteHW | 0.97 | 0.22 | 4.44 | *** |
| siteJB | -0.12 | 0.11 | -1.11 | - |
| siteLU | -0.53 | 0.18 | -2.89 | ** |
| sitePH | -0.78 | 0.15 | -5.26 | *** |
| siteSK | 0.02 | 0.11 | 0.19 | - |
| siteST | 0.24 | 0.12 | 1.98 | * |
| siteWF+WT | -0.43 | 0.12 | -3.49 | *** |
| IL17a_U(GA) | -0.39 | 0.44 | -0.90 | - |
| IL17a_U(GG) | -0.96 | 0.20 | -4.79 | *** |
| age_cl(IM) | -0.60 | 0.06 | -10.30 | *** |
| age_cl(YO) | -0.32 | 0.06 | -5.22 | *** |
| sex(M) | -0.11 | 0.05 | -2.48 | * |
| siteGL x IL17a_U(GA) | 0.68 | 0.48 | 1.41 | - |
| siteHW x IL17a_U(GA) | -0.25 | 0.65 | -0.39 | - |
| siteLU x IL17a_U(GA) | 1.07 | 0.56 | 1.92 | . |
| sitePH x IL17a_U(GA) | 0.34 | 0.47 | 0.72 | - |
| siteGL x IL17a_U(GG) | 1.60 | 0.51 | 3.15 | ** |
| siteJB x IL17a_U(GG) | 0.78 | 0.48 | 1.61 | - |
| sitePH x IL17a_U(GG) | 0.79 | 0.25 | 3.09 | ** |
| siteSK x IL17a_U(GG) | 1.19 | 0.49 | 2.44 | * |

**IgE ~ IL-1b_U + site + sex + age (classes)**

| | Estimate | ±SE | t value | p |
|---|---|---|---|---|
| (Intercept) | 2.09 | 0.12 | 17.53 | *** |
| siteGL | -0.18 | 0.11 | -1.59 | - |
| siteHW | 0.17 | 0.09 | 1.85 | . |
| siteJB | -0.17 | 0.11 | -1.55 | - |
| siteLU | -0.37 | 0.17 | -2.18 | * |
| sitePH | -0.57 | 0.12 | -4.73 | *** |
| siteSK | 0.32 | 0.14 | 2.27 | * |
| siteST | 0.06 | 0.11 | 0.48 | - |
| siteWF+WT | -0.45 | 0.12 | -3.75 | *** |
| sex(M) | -0.13 | 0.05 | -2.81 | ** |
| age_cl(IM) | -0.61 | 0.06 | -10.44 | *** |
| age_cl(YO) | -0.30 | 0.06 | -4.94 | *** |
| IL1b_U(GT) | 0.08 | 0.07 | 1.15 | - |
| IL1b_U(TT) | 0.31 | 0.09 | 3.47 | *** |

| IgE ~ site x IL6 + age (classes) + sex | | | | |
|---|---|---|---|---|
| | Estimate | ±SE | t value | p |
| (Intercept) | 2.44 | 0.11 | 23.18 | *** |
| siteGL | -0.24 | 0.13 | -1.86 | . |
| siteHW | -0.08 | 0.11 | -0.76 | |
| siteJB | -0.24 | 0.13 | -1.93 | . |
| siteLU | -0.44 | 0.18 | -2.43 | * |
| sitePH | -0.96 | 0.11 | -8.41 | *** |
| siteSK | -0.04 | 0.13 | -0.29 | |
| siteST | -0.08 | 0.14 | -0.58 | |
| siteW | -0.71 | 0.33 | -2.12 | * |
| IL6(CT) | -0.06 | 0.17 | -0.34 | |
| IL6(TT) | -0.82 | 0.28 | -2.96 | ** |
| age_cl(IM) | -0.57 | 0.06 | -9.55 | *** |
| age_cl(YO) | -0.28 | 0.06 | -4.58 | *** |
| sex(M) | -0.10 | 0.05 | -2.25 | * |
| siteST x IL6(CT) | 0.14 | 0.28 | 0.51 | |
| siteW x IL6(CT) | 0.04 | 0.38 | 0.11 | |
| siteW x IL6(TT) | 1.18 | 0.45 | 2.60 | ** |

| IgE ~ site x Tnf + age (classes) + sex | | | | |
|---|---|---|---|---|
| | Estimate | ±SE | t value | p |
| (Intercept) | 2.38 | 0.09 | 27.19 | *** |
| siteGL | -0.17 | 0.11 | -1.48 | - |
| siteHW | 0.00 | 0.09 | -0.05 | - |
| siteJB | -0.17 | 0.11 | -1.54 | - |
| siteLU | -0.38 | 0.33 | -1.15 | - |
| sitePH | -0.89 | 0.10 | -9.03 | *** |
| siteSK | 0.02 | 0.12 | 0.17 | - |
| siteST | -0.12 | 0.13 | -0.85 | - |
| siteW | -0.38 | 0.13 | -2.91 | ** |
| Tnf(TG) | -0.31 | 0.33 | -0.95 | - |
| Tnf(TT) | -0.26 | 0.34 | -0.77 | - |
| age_cl(IM) | -0.59 | 0.06 | -9.98 | *** |
| age_cl(YO) | -0.31 | 0.06 | -5.06 | *** |
| sex(M) | -0.10 | 0.05 | -2.10 | * |
| siteLU x Tnf(TG) | 0.30 | 0.49 | 0.62 | - |
| siteST x Tnf(TG) | 0.72 | 0.38 | 1.89 | - |
| siteW x Tnf(TG) | -0.57 | 0.43 | -1.33 | - |
| siteST x Tnf(TT) | 0.35 | 0.47 | 0.75 | - |

#### Cytokines

| PC1 cytokines ~ site x sex + H2Eb | | | | |
|---|---|---|---|---|
| | Estimate | ±SE | t value | p |
| (Intercept) | -6.76 | 1.75 | -3.87 | *** |
| siteGL | 3.90 | 1.69 | 2.31 | * |
| siteHW | 8.64 | 1.62 | 5.34 | *** |
| siteJB | 8.30 | 1.77 | 4.69 | *** |
| siteLU | 1.12 | 2.48 | 0.45 | |
| sitePH | 9.46 | 1.94 | 4.88 | *** |
| siteSK | 6.62 | 1.86 | 3.56 | *** |
| siteST | 8.65 | 2.41 | 3.58 | *** |
| siteW | 6.83 | 2.01 | 3.40 | *** |
| sex(M) | 6.40 | 1.77 | 3.62 | *** |
| H2Eb(AG) | -1.75 | 1.06 | -1.66 | - |
| H2Eb(GG) | -0.04 | 1.02 | -0.04 | - |
| siteGL x sex(M) | -4.51 | 2.26 | -2.00 | * |
| siteHW x sex(M) | -7.10 | 1.95 | -3.63 | *** |
| siteJB x sex(M) | -7.96 | 2.22 | -3.59 | *** |
| sitePH x sex(M) | -7.99 | 2.36 | -3.39 | *** |
| siteSK x sex(M) | -5.06 | 2.66 | -1.90 | - |
| siteST x sex(M) | -7.10 | 2.78 | -2.55 | * |
| siteW x sex(M) | -9.74 | 2.92 | -3.34 | ** |

| PC1 cytokines ~ site x sex + IL1b_U | | | | |
|---|---|---|---|---|
| | Estimate | ±SE | t value | p |
| (Intercept) | -5.45 | 1.68 | -3.24 | ** |
| siteGL | 4.32 | 1.69 | 2.55 | * |
| siteHW | 6.85 | 1.61 | 4.27 | *** |
| siteJB | 8.73 | 1.77 | 4.92 | *** |
| siteLU | 1.36 | 2.48 | 0.55 | - |
| sitePH | 7.37 | 2.03 | 3.62 | *** |
| siteSK | 5.27 | 2.06 | 2.55 | * |
| siteST | 9.10 | 2.32 | 3.93 | *** |
| siteW | 7.25 | 2.01 | 3.61 | *** |
| sex(M) | 6.58 | 1.77 | 3.73 | *** |
| IL1b_U(GT) | 0.33 | 0.84 | 0.40 | - |
| IL1b_U(TT) | -1.78 | 1.00 | -1.78 | - |
| siteGL x sex(M) | -4.89 | 2.25 | -2.17 | * |
| siteHW x sex(M) | -7.34 | 1.96 | -3.75 | *** |
| siteJB x sex(M) | -8.27 | 2.21 | -3.73 | *** |
| sitePH x sex(M) | -7.63 | 2.36 | -3.24 | ** |
| siteSK x sex(M) | -5.24 | 2.66 | -1.97 | - |
| siteST x sex(M) | -7.82 | 2.79 | -2.80 | ** |
| siteW x sex(M) | -9.92 | 2.91 | -3.41 | *** |

| PC1 cytokines ~ site x sex + IL6 | | | | |
|---|---|---|---|---|
| | Estimate | ±SE | t value | p |
| (Intercept) | -8.87 | 1.66 | -5.36 | *** |
| siteGL | 5.96 | 1.89 | 3.15 | ** |
| siteHW | 9.82 | 1.77 | 5.54 | *** |
| siteJB | 10.37 | 1.96 | 5.29 | *** |
| siteLU | 2.81 | 2.56 | 1.10 | - |
| sitePH | 10.95 | 2.10 | 5.22 | *** |
| siteSK | 8.68 | 2.04 | 4.26 | *** |
| siteST | 10.74 | 2.46 | 4.36 | *** |
| siteW | 4.71 | 3.26 | 1.45 | - |
| sex(M) | 6.77 | 1.90 | 3.56 | *** |
| IL6(CT) | 3.44 | 1.44 | 2.38 | * |
| IL6(TT) | 4.18 | 2.56 | 1.63 | - |
| siteGL x sex(M) | -5.07 | 2.36 | -2.15 | * |
| siteHW x sex(M) | -7.34 | 2.07 | -3.54 | *** |
| siteJB x sex(M) | -8.45 | 2.32 | -3.64 | *** |
| sitePH x sex(M) | -7.91 | 2.45 | -3.23 | ** |
| siteSK x sex(M) | -5.42 | 2.75 | -1.98 | * |
| siteST x sex(M) | -8.33 | 2.94 | -2.84 | ** |
| siteW x sex(M) | -8.21 | 4.36 | -1.89 | - |

| PC1 cytokines ~ site x sex + IL10 | | | | |
|---|---|---|---|---|
| | Estimate | ±SE | t value | p |
| (Intercept) | -6.81 | 1.34 | -5.09 | *** |
| siteGL | 6.42 | 1.67 | 3.84 | *** |
| siteHW | 7.76 | 1.47 | 5.30 | *** |
| siteJB | 8.31 | 1.67 | 4.98 | *** |
| siteLU | 1.12 | 2.34 | 0.48 | - |
| sitePH | 8.89 | 1.81 | 4.91 | *** |
| siteSK | 6.62 | 1.75 | 3.78 | *** |
| siteST | 8.68 | 2.19 | 3.97 | *** |
| siteW | 6.83 | 1.89 | 3.61 | *** |
| sex(M) | 6.40 | 1.67 | 3.83 | *** |
| IL10(CT) | -6.05 | 1.22 | -4.94 | *** |
| siteGL x sex(M) | -5.87 | 2.14 | -2.74 | ** |
| siteHW x sex(M) | -6.97 | 1.84 | -3.78 | *** |
| siteJB x sex(M) | -8.08 | 2.09 | -3.86 | *** |
| sitePH x sex(M) | -7.54 | 2.22 | -3.40 | *** |
| siteSK x sex(M) | -5.06 | 2.51 | -2.01 | * |
| siteST x sex(M) | -5.59 | 2.64 | -2.12 | * |
| siteW x sex(M) | -9.74 | 2.75 | -3.54 | *** |

| PC1 ~ site x sex + Tnf | | | | |
|---|---|---|---|---|
| | Estimate | ±SE | t value | p |
| (Intercept) | -6.81 | 1.38 | -4.92 | *** |
| siteGL | 3.90 | 1.65 | 2.37 | * |
| siteHW | 7.76 | 1.51 | 5.12 | *** |
| siteJB | 8.30 | 1.72 | 4.82 | *** |
| siteLU | 0.21 | 2.69 | 0.08 | - |
| sitePH | 8.89 | 1.87 | 4.75 | *** |
| siteSK | 6.62 | 1.81 | 3.66 | *** |
| siteST | 7.90 | 2.48 | 3.19 | ** |
| siteW | 6.83 | 1.96 | 3.49 | *** |
| sex(M) | 6.14 | 1.76 | 3.49 | *** |
| TnfT x G | 1.17 | 1.53 | 0.77 | - |
| Tnf(TT) | -13.87 | 3.79 | -3.66 | *** |
| siteGL x sex(M) | -4.44 | 2.22 | -2.00 | * |
| siteHW x sex(M) | -6.71 | 1.93 | -3.47 | *** |
| siteJB x sex(M) | -7.82 | 2.19 | -3.57 | *** |
| sitePH x sex(M) | -7.28 | 2.32 | -3.14 | ** |
| siteSK x sex(M) | -4.80 | 2.62 | -1.83 | - |
| siteST x sex(M) | -6.50 | 2.82 | -2.30 | * |
| siteW x sex(M) | -4.85 | 3.13 | -1.55 | - |

| PC2 cytokines ~ site x sex + IL10 | | | | |
|---|---|---|---|---|
| | Estimate | ±SE | t value | p |
| (Intercept) | -3.36 | 1.04 | -3.24 | ** |
| siteGL | 3.16 | 1.30 | 2.43 | * |
| siteHW | 3.75 | 1.14 | 3.29 | ** |
| siteJB | 2.95 | 1.30 | 2.27 | * |
| siteLU | -0.94 | 1.82 | -0.52 | - |
| sitePH | 2.43 | 1.41 | 1.72 | - |
| siteSK | 5.96 | 1.36 | 4.38 | *** |
| siteST | 2.04 | 1.70 | 1.20 | - |
| siteW | 5.18 | 1.47 | 3.52 | *** |
| sex(M) | 2.24 | 1.30 | 1.73 | - |
| IL10(CT) | -3.13 | 0.95 | -3.29 | ** |
| siteGL x sex(M) | -1.09 | 1.66 | -0.65 | - |
| siteHW x sex(M) | -3.00 | 1.43 | -2.10 | * |
| siteJB x sex(M) | 0.49 | 1.63 | 0.30 | - |
| sitePH x sex(M) | -0.56 | 1.72 | -0.32 | - |
| siteSK x sex(M) | -3.91 | 1.95 | -2.01 | * |
| siteST x sex(M) | -0.83 | 2.05 | -0.40 | - |
| siteW x sex(M) | -6.20 | 2.14 | -2.90 | ** |

| PC2 cytokines ~ site x sex + IL17a_U | | | | |
|---|---|---|---|---|
| | Estimate | ±SE | t value | p |
| (Intercept) | -3.36 | 1.06 | -3.17 | ** |
| siteGL | 0.88 | 1.50 | 0.59 | - |
| siteHW | 4.73 | 1.47 | 3.22 | ** |
| siteJB | 2.95 | 1.32 | 2.23 | * |
| siteLU | -2.21 | 2.07 | -1.07 | - |
| sitePH | 2.12 | 1.58 | 1.34 | - |
| siteSK | 5.96 | 1.39 | 4.30 | *** |
| siteST | 2.36 | 1.76 | 1.35 | - |
| siteW | 5.18 | 1.50 | 3.45 | *** |
| sex(M) | 2.24 | 1.32 | 1.70 | - |
| IL17a_U(GA) | 1.27 | 0.92 | 1.37 | - |
| IL17a_U(GG) | -0.98 | 0.90 | -1.09 | - |
| siteGL x sex(M) | -0.35 | 1.70 | -0.21 | - |
| siteHW x sex(M) | -3.00 | 1.46 | -2.06 | * |
| siteJB x sex(M) | 0.55 | 1.66 | 0.33 | - |
| sitePH x sex(M) | -0.28 | 1.76 | -0.16 | - |
| siteSK x sex(M) | -3.91 | 1.99 | -1.97 | - |
| siteST x sex(M) | -1.69 | 2.08 | -0.81 | - |
| siteW x sex(M) | -6.20 | 2.18 | -2.85 | ** |

### Cells

| NK cells ~ site x age (linear) + SMI + IL6 | | | | |
|---|---|---|---|---|
| | Estimate | ±SE | t value | p |
| (Intercept) | 4.13 | 0.22 | 19.18 | *** |
| siteGL | 0.36 | 0.24 | 1.52 | - |
| siteHW | 0.12 | 0.19 | 0.63 | - |
| siteJB | 0.77 | 0.22 | 3.52 | *** |
| siteLU | 1.75 | 0.72 | 2.43 | * |
| sitePH | -0.05 | 0.20 | -0.26 | - |
| siteSK | 0.53 | 0.24 | 2.20 | * |
| siteST | 0.15 | 0.22 | 0.67 | - |
| siteW | 0.59 | 0.25 | 2.31 | * |
| age_l | -0.01 | 0.01 | -0.63 | - |
| SMI | 0.10 | 0.01 | 11.35 | *** |
| IL6(CT) | -0.17 | 0.14 | -1.25 | - |
| IL6(TT) | 0.35 | 0.19 | 1.81 | - |
| siteGL x age_l | -0.02 | 0.02 | -1.30 | - |
| siteHW x age_l | -0.01 | 0.01 | -1.16 | - |
| siteJB x age_l | -0.03 | 0.02 | -2.24 | * |
| siteLU x age_l | -0.09 | 0.03 | -2.80 | ** |
| sitePH x age_l | -0.02 | 0.01 | -1.33 | - |
| siteSK x age_l | -0.04 | 0.02 | -2.38 | * |
| siteST x age_l | -0.01 | 0.02 | -0.35 | - |
| siteW x age_l | -0.06 | 0.02 | -2.96 | ** |

| NK cells ~ site x age (linear) + SMI + IL10 | | | | |
|---|---|---|---|---|
| | Estimate | ±SE | t value | p |
| (Intercept) | 3.97 | 0.27 | 14.57 | *** |
| siteGL | 0.41 | 0.23 | 1.79 | . |
| siteHW | -0.02 | 0.18 | -0.10 | |
| siteJB | 0.63 | 0.21 | 3.04 | ** |
| siteLU | 1.62 | 0.72 | 2.26 | * |
| sitePH | -0.20 | 0.19 | -1.03 | |
| siteSK | 0.67 | 0.28 | 2.38 | * |
| siteST | 0.07 | 0.22 | 0.31 | |
| siteW | 0.59 | 0.24 | 2.42 | * |
| age_l | -0.02 | 0.01 | -1.37 | |
| SMI | 0.10 | 0.01 | 11.85 | *** |
| IL10(CT) | -0.15 | 0.20 | -0.74 | |
| IL10(TT) | 0.27 | 0.18 | 1.53 | |
| siteGL x age_l | -0.02 | 0.02 | -0.99 | |
| siteHW x age_l | -0.01 | 0.01 | -0.53 | |
| siteJB x age_l | -0.03 | 0.02 | -1.74 | . |
| siteLU x age_l | -0.08 | 0.03 | -2.57 | * |
| sitePH x age_l | -0.01 | 0.01 | -0.66 | |
| siteSK x age_l | -0.03 | 0.02 | -1.99 | * |
| siteST x age_l | 0.01 | 0.02 | 0.29 | |
| siteW x age_l | -0.07 | 0.02 | -3.41 | *** |

| NK cells ~ site x age (linear) + SMI + Tnf | | | | |
|---|---|---|---|---|
| | Estimate | ±SE | t value | p |
| (Intercept) | 4.16 | 0.21 | 19.78 | *** |
| siteGL | 0.28 | 0.23 | 1.22 | - |
| siteHW | 0.04 | 0.18 | 0.21 | - |
| siteJB | 0.68 | 0.21 | 3.31 | ** |
| siteLU | 1.97 | 0.74 | 2.68 | ** |
| sitePH | -0.14 | 0.19 | -0.76 | - |
| siteSK | 0.47 | 0.23 | 2.05 | * |
| siteST | 0.27 | 0.23 | 1.18 | - |
| siteW | 0.74 | 0.24 | 3.03 | ** |
| age_l | -0.01 | 0.01 | -1.15 | - |
| SMI | 0.10 | 0.01 | 12.01 | *** |
| Tnf(TG) | -0.34 | 0.13 | -2.59 | * |
| Tnf(TT) | -0.50 | 0.23 | -2.21 | * |
| siteGL x age_l | -0.02 | 0.02 | -0.97 | - |
| siteHW x age_l | -0.01 | 0.01 | -0.75 | - |
| siteJB x age_l | -0.03 | 0.02 | -1.89 | - |
| siteLU x age_l | -0.09 | 0.03 | -2.61 | ** |
| sitePH x age_l | -0.01 | 0.01 | -0.86 | - |
| siteSK x age_l | -0.04 | 0.02 | -2.16 | * |
| siteST x age_l | -0.01 | 0.02 | -0.65 | - |
| siteW x age_l | -0.07 | 0.02 | -3.62 | *** |

| NK cells ~ age (linear) x site + SMI + IL13 | | | | |
|---|---|---|---|---|
| | Estimate | ±SE | t value | p |
| (Intercept) | 3.92 | 0.24 | 16.50 | *** |
| age_l | 0.00 | 0.01 | -0.33 | - |
| siteGL | 0.57 | 0.25 | 2.25 | * |
| siteHW | 0.33 | 0.21 | 1.58 | - |
| siteJB | 0.98 | 0.23 | 4.21 | *** |
| siteLU | 2.07 | 0.74 | 2.80 | ** |
| sitePH | 0.16 | 0.22 | 0.73 | - |
| siteSK | 0.79 | 0.26 | 3.08 | ** |
| siteST | 0.29 | 0.24 | 1.19 | - |
| siteW | 0.93 | 0.27 | 3.51 | *** |
| SMI | 0.10 | 0.01 | 11.17 | *** |
| IL13 (TC) | 0.33 | 0.16 | 2.08 | * |
| IL13 (TT) | 0.86 | 0.43 | 2.02 | * |
| age_l x siteGL | -0.02 | 0.02 | -1.47 | - |
| age_l x siteHW | -0.02 | 0.01 | -1.37 | - |
| age_l x siteJB | -0.04 | 0.02 | -2.42 | * |
| age_l x siteLU | -0.10 | 0.03 | -3.02 | ** |
| age_l x sitePH | -0.02 | 0.01 | -1.59 | - |
| age_l x siteSK | -0.05 | 0.02 | -2.74 | ** |
| age_l x siteST | -0.01 | 0.02 | -0.52 | - |
| age_l x siteW | -0.08 | 0.02 | -3.84 | *** |

| B cells ~ age (linear) x sex + site + SMI + IL10 | | | | |
|---|---|---|---|---|
| | Estimate | ±SE | t value | p |
| (Intercept) | 5.37 | 0.23 | 22.96 | *** |
| age_l | -0.03 | 0.00 | -7.83 | *** |
| sex (M) | -0.21 | 0.08 | -2.71 | ** |
| siteGL | 0.35 | 0.13 | 2.69 | ** |
| siteHW | 0.13 | 0.11 | 1.26 | - |
| siteJB | 0.17 | 0.12 | 1.40 | - |
| siteLU | -0.27 | 0.16 | -1.67 | - |
| sitePH | -0.15 | 0.11 | -1.31 | - |
| siteSK | 0.40 | 0.21 | 1.86 | - |
| siteST | 0.39 | 0.14 | 2.79 | ** |
| siteW | 0.01 | 0.13 | 0.06 | - |
| SMI | 0.10 | 0.01 | 12.84 | *** |
| IL10(CT) | -0.16 | 0.20 | -0.79 | - |
| IL10(TT) | 0.34 | 0.18 | 1.90 | - |
| age_l x se x M | 0.01 | 0.01 | 1.43 | - |

| B cells ~ age (linear) x sex + site + SMI + Tnf | | | | |
|---|---|---|---|---|
| | Estimate | ±SE | t value | p |
| (Intercept) | 5.70 | 0.16 | 36.50 | *** |
| age_l | -0.03 | 0.00 | -7.80 | *** |
| sex (M) | -0.21 | 0.08 | -2.69 | ** |
| siteGL | 0.19 | 0.13 | 1.50 | - |
| siteHW | 0.14 | 0.11 | 1.30 | - |
| siteJB | 0.18 | 0.13 | 1.43 | - |
| siteLU | 0.05 | 0.19 | 0.26 | - |
| sitePH | -0.15 | 0.12 | -1.31 | - |
| siteSK | 0.08 | 0.13 | 0.60 | - |
| siteST | 0.40 | 0.14 | 2.82 | ** |
| siteW | 0.09 | 0.13 | 0.70 | - |
| SMI | 0.10 | 0.01 | 12.71 | *** |
| Tnf(TG) | -0.40 | 0.13 | -3.09 | ** |
| Tnf(TT) | -0.32 | 0.19 | -1.66 | - |
| age_l x sex (M) | 0.01 | 0.01 | 1.66 | - |

| B cells ~ age (linear) x sex + site + SMI + IL13 | | | | |
|---|---|---|---|---|
| | Estimate | ±SE | t value | p |
| (Intercept) | 5.56 | 0.17 | 32.38 | *** |
| age_l | -0.03 | 0.00 | -7.42 | *** |
| sexM | -0.21 | 0.08 | -2.64 | ** |
| siteGL | 0.36 | 0.14 | 2.52 | * |
| siteHW | 0.31 | 0.13 | 2.47 | * |
| siteJB | 0.35 | 0.14 | 2.47 | * |
| siteLU | -0.12 | 0.17 | -0.69 | - |
| sitePH | 0.02 | 0.14 | 0.18 | - |
| siteSK | 0.22 | 0.15 | 1.49 | - |
| siteST | 0.38 | 0.15 | 2.59 | ** |
| siteW | 0.19 | 0.15 | 1.26 | - |
| SMI | 0.10 | 0.01 | 12.08 | *** |
| IL13 (TC) | 0.24 | 0.15 | 1.56 | - |
| IL13 (TT) | 1.00 | 0.44 | 2.26 | * |
| age_l x sex(M) | 0.01 | 0.01 | 1.30 | - |

| DC ~ H2Aa + site x age (linear) + SMI | | | | |
|---|---|---|---|---|
| | Estimate | ±SE | t value | p |
| (Intercept) | 4.75 | 0.33 | 14.53 | *** |
| siteGL | -0.04 | 0.32 | -0.14 | - |
| siteHW | -0.06 | 0.24 | -0.25 | - |
| siteJB | 0.52 | 0.26 | 1.98 | * |
| siteLU | 0.53 | 0.51 | 1.06 | - |
| sitePH | 0.06 | 0.25 | 0.24 | - |
| siteSK | 0.49 | 0.29 | 1.70 | . |
| siteST | -0.06 | 0.28 | -0.22 | |
| siteWF+WT | 0.64 | 0.30 | 2.14 | * |
| age_l | -0.02 | 0.01 | -1.24 | - |
| SMI | 0.11 | 0.01 | 10.38 | *** |
| H2Aa(AG) | -0.47 | 0.16 | -2.85 | ** |
| H2Aa(GG) | -0.27 | 0.18 | -1.51 | - |
| siteGL x age_l | 0.00 | 0.02 | 0.06 | - |
| siteHW x age_l | -0.01 | 0.02 | -0.78 | - |
| siteJB x age_l | -0.05 | 0.02 | -2.84 | ** |
| siteLU x age_l | -0.03 | 0.02 | -1.04 | - |
| sitePH x age_l | -0.01 | 0.02 | -0.98 | - |
| siteSK x age_l | -0.05 | 0.02 | -2.33 | * |
| siteST x age_l | 0.01 | 0.02 | 0.47 | - |
| siteWF+WT x age_l | -0.06 | 0.02 | -2.80 | ** |

| DC ~ site x age (linear) + SMI + IL6 | | | | |
|---|---|---|---|---|
| | Estimate | ±SE | t value | p |
| (Intercept) | 4.44 | 0.29 | 15.39 | *** |
| siteGL | 0.15 | 0.30 | 0.51 | - |
| siteHW | 0.03 | 0.25 | 0.11 | - |
| siteJB | 0.65 | 0.28 | 2.33 | * |
| siteLU | 0.60 | 0.51 | 1.18 | - |
| sitePH | 0.06 | 0.26 | 0.21 | - |
| siteSK | 0.59 | 0.31 | 1.93 | - |
| siteST | 0.07 | 0.28 | 0.25 | - |
| siteW | 0.42 | 0.30 | 1.41 | - |
| age_l | -0.01 | 0.01 | -0.81 | - |
| SMI | 0.10 | 0.01 | 10.12 | *** |
| IL6(CT) | -0.19 | 0.17 | -1.10 | - |
| IL6(TT) | 0.35 | 0.22 | 1.61 | - |
| siteGL x age_l | 0.00 | 0.02 | -0.04 | - |
| siteHW x age_l | -0.02 | 0.02 | -1.12 | - |
| siteJB x age_l | -0.06 | 0.02 | -3.28 | ** |
| siteLU x age_l | -0.03 | 0.02 | -1.26 | - |
| sitePH x age_l | -0.02 | 0.01 | -1.29 | - |
| siteSK x age_l | -0.05 | 0.02 | -2.63 | ** |
| siteST x age_l | 0.00 | 0.02 | 0.12 | - |
| siteW x age_l | -0.04 | 0.02 | -2.04 | * |

| DC ~ site x age (linear) + SMI + IL10 | | | | |
|---|---|---|---|---|
| | Estimate | ±SE | t value | p |
| (Intercept) | 3.90 | 0.35 | 11.18 | *** |
| siteGL | 0.46 | 0.29 | 1.59 | - |
| siteHW | -0.04 | 0.23 | -0.19 | - |
| siteJB | 0.53 | 0.26 | 2.08 | * |
| siteLU | 0.54 | 0.49 | 1.10 | - |
| sitePH | -0.02 | 0.24 | -0.09 | - |
| siteSK | 1.06 | 0.35 | 3.06 | ** |
| siteST | 0.21 | 0.28 | 0.76 | - |
| siteW | 0.49 | 0.28 | 1.77 | - |
| age_l | -0.02 | 0.01 | -1.27 | - |
| SMI | 0.10 | 0.01 | 10.68 | *** |
| IL10(CT) | -0.10 | 0.25 | -0.39 | - |
| IL10(TT) | 0.60 | 0.22 | 2.72 | ** |
| siteGL x age_l | -0.01 | 0.02 | -0.36 | - |
| siteHW x age_l | -0.01 | 0.01 | -0.79 | - |
| siteJB x age_l | -0.05 | 0.02 | -2.91 | ** |
| siteLU x age_l | -0.03 | 0.02 | -1.06 | - |
| sitePH x age_l | -0.01 | 0.01 | -0.94 | - |
| siteSK x age_l | -0.04 | 0.02 | -2.36 | * |
| siteST x age_l | 0.00 | 0.02 | 0.22 | - |
| siteW x age_l | -0.05 | 0.02 | -2.36 | * |

| DC ~ age (linear) x site + SMI + IL13 | | | | |
|---|---|---|---|---|
| | Estimate | ±SE | t value | p |
| (Intercept) | 4.22 | 0.32 | 13.11 | *** |
| age_l | -0.01 | 0.01 | -0.51 | - |
| siteGL | 0.39 | 0.32 | 1.20 | - |
| siteHW | 0.26 | 0.28 | 0.93 | - |
| siteJB | 0.84 | 0.31 | 2.75 | ** |
| siteLU | 0.87 | 0.54 | 1.62 | - |
| sitePH | 0.29 | 0.29 | 1.02 | - |
| siteSK | 0.86 | 0.33 | 2.59 | ** |
| siteST | 0.23 | 0.31 | 0.73 | - |
| siteW | 0.80 | 0.33 | 2.44 | * |
| SMI | 0.10 | 0.01 | 9.82 | *** |
| IL13 (TC) | 0.31 | 0.21 | 1.49 | - |
| IL13 (TT) | 1.17 | 0.50 | 2.32 | * |
| age_l x siteGL | 0.00 | 0.02 | -0.22 | - |
| age_l x siteHW | -0.02 | 0.02 | -1.28 | - |
| age_l x siteJB | -0.06 | 0.02 | -3.21 | ** |
| age_l x siteLU | -0.04 | 0.03 | -1.45 | - |
| age_l x sitePH | -0.02 | 0.02 | -1.48 | - |
| age_l x siteSK | -0.06 | 0.02 | -2.87 | ** |
| age_l x siteST | 0.00 | 0.02 | 0.02 | - |
| age_l x siteW | -0.06 | 0.02 | -2.61 | ** |

| CD8+ T cells ~ H2Aa + age (linear) + SMI | | | | |
|---|---|---|---|---|
| | Estimate | ±SE | t value | p |
| (Intercept) | 5.72 | 0.11 | 50.27 | *** |
| age_l | -0.02 | 0.00 | -8.26 | *** |
| SMI | 0.08 | 0.01 | 12.16 | *** |
| H2Aa(AG) | -0.14 | 0.09 | -1.55 | - |
| H2Aa(GG) | -0.17 | 0.07 | -2.44 | * |

| CD8+ T cells ~ age (linear) + SMI + IL6 | | | | |
|---|---|---|---|---|
| | Estimate | ±SE | t value | p |
| (Intercept) | 5.56491 | 0.09 | 58.99 | *** |
| age_l | -0.0191 | 0 | -8.28 | *** |
| SMI | 0.07509 | 0.01 | 12.29 | *** |
| IL6(CT) | -0.1048 | 0.07 | -1.42 | - |
| IL6(TT) | 0.23306 | 0.1 | 2.317 | * |

| CD8+ T cells ~ age (linear) + SMI + IL10 | | | | |
|---|---|---|---|---|
| | Estimate | ±SE | t value | p |
| (Intercept) | 5.47913 | 0.11 | 50.2 | *** |
| age_l | -0.0198 | 0 | -8.619 | *** |
| SMI | 0.07426 | 0.01 | 12.25 | *** |
| IL10(CT) | -0.0649 | 0.11 | -0.584 | - |
| IL10(TT) | 0.12572 | 0.07 | 1.868 | - |

| CD4+ T cells~ H2Aa x site + age (linear) + SMI | | | | |
|---|---|---|---|---|
| | Estimate | ±SE | t value | p |
| (Intercept) | 5.57 | 0.19 | 29.02 | *** |
| siteGL | 0.48 | 0.19 | 2.59 | * |
| siteHW | 0.09 | 0.08 | 1.09 | - |
| siteJB | 0.12 | 0.09 | 1.37 | - |
| siteLU | 0.08 | 0.11 | 0.74 | - |
| sitePH | -0.01 | 0.09 | -0.06 | - |
| siteSK | 0.18 | 0.09 | 1.89 | . |
| siteST | 0.11 | 0.10 | 1.17 | - |
| siteWF+WT | 0.06 | 0.11 | 0.52 | - |
| H2Aa(AG) | 0.27 | 0.34 | 0.79 | - |
| H2Aa(GG) | 0.16 | 0.16 | 1.00 | - |
| age_l | -0.02 | 0.00 | -7.58 | *** |
| SMI | 0.07 | 0.01 | 13.00 | *** |
| siteGL x H2Aa(AG) | -0.75 | 0.37 | -2.05 | * |
| sitePH x H2Aa(AG) | -0.15 | 0.32 | -0.47 | - |
| siteST x H2Aa(AG) | 0.52 | 0.42 | 1.22 | - |
| siteWF+WT x H2Aa(AG) | -0.10 | 0.33 | -0.29 | - |
| siteGL x H2Aa(GG) | -0.67 | 0.34 | -1.97 | * |

| CD4+ T cells ~ H2Eb x site + age (linear) + SMI | | | | |
|---|---|---|---|---|
| | Estimate | ±SE | t value | p |
| (Intercept) | 5.32 | 0.17 | 30.43 | *** |
| siteGL | 0.21 | 0.09 | 2.26 | * |
| siteHW | 0.64 | 0.16 | 3.91 | *** |
| siteJB | 0.65 | 0.33 | 1.99 | * |
| siteLU | 0.08 | 0.11 | 0.73 | - |
| sitePH | -0.07 | 0.08 | -0.80 | - |
| siteSK | 0.16 | 0.09 | 1.69 | - |
| siteST | 0.36 | 0.12 | 3.02 | ** |
| siteWF+WT | 0.06 | 0.10 | 0.61 | - |
| H2Eb(AG) | 0.37 | 0.20 | 1.85 | - |
| H2Eb(GG) | 0.44 | 0.13 | 3.44 | *** |
| age_l | -0.02 | 0.00 | -8.23 | *** |
| SMI | 0.07 | 0.01 | 12.59 | *** |
| siteGL x H2Eb(AG) | -0.09 | 0.27 | -0.34 | - |
| siteHW x H2Eb(AG) | -0.45 | 0.22 | -2.07 | * |
| siteHW x H2Eb(GG) | -0.64 | 0.15 | -4.22 | *** |
| siteJB x H2Eb(GG) | -0.54 | 0.33 | -1.67 | - |

| CD4+ T cells ~ site x IL6 + age (linear) + SMI | | | | |
|---|---|---|---|---|
| | Estimate | ±SE | t value | p |
| (Intercept) | 5.55 | 0.13 | 44.02 | *** |
| siteGL | 0.41 | 0.12 | 3.47 | *** |
| siteHW | 0.29 | 0.11 | 2.69 | ** |
| siteJB | 0.33 | 0.12 | 2.77 | ** |
| siteLU | 0.29 | 0.14 | 2.16 | * |
| sitePH | 0.14 | 0.11 | 1.25 | - |
| siteSK | 0.38 | 0.12 | 3.15 | ** |
| siteST | 0.41 | 0.13 | 3.27 | ** |
| siteW | -0.02 | 0.23 | -0.07 | - |
| IL6(CT) | 0.36 | 0.16 | 2.29 | * |
| IL6(TT) | 0.47 | 0.20 | 2.34 | * |
| age_l | -0.02 | 0.00 | -8.18 | *** |
| SMI | 0.07 | 0.01 | 12.82 | *** |
| siteST x IL6(CT) | -0.69 | 0.22 | -3.15 | ** |
| siteW x IL6(CT) | -0.19 | 0.28 | -0.68 | - |
| siteW x IL6(TT) | 0.08 | 0.31 | 0.26 | - |

| CD4 T cells ~ site x IL10 + age (linear) + SMI | | | | |
|---|---|---|---|---|
| | Estimate | ±SE | t value | p |
| (Intercept) | 5.73 | 0.24 | 23.65 | *** |
| siteGL | -0.40 | 0.31 | -1.31 | - |
| siteHW | 0.05 | 0.08 | 0.60 | - |
| siteJB | 0.08 | 0.09 | 0.91 | - |
| siteLU | 0.07 | 0.11 | 0.58 | - |
| sitePH | -0.10 | 0.08 | -1.23 | - |
| siteSK | 0.18 | 0.24 | 0.75 | - |
| siteST | 0.07 | 0.10 | 0.67 | - |
| siteW | 0.05 | 0.10 | 0.49 | - |
| IL10(CT) | -0.42 | 0.37 | -1.14 | - |
| IL10(TT) | 0.05 | 0.22 | 0.22 | - |
| age_l | -0.02 | 0.00 | -8.84 | *** |
| SMI | 0.07 | 0.01 | 13.36 | *** |
| siteGL x IL10(CT) | 0.73 | 0.44 | 1.67 | - |
| siteST x IL10(CT) | 0.61 | 0.34 | 1.82 | - |
| siteGL x IL10(TT) | 0.77 | 0.31 | 2.50 | * |
| siteSK x IL10(TT) | 0.21 | 0.37 | 0.56 | - |

| CD4 T cells ~ site + age (linear) + SMI + Tnf | | | | |
|---|---|---|---|---|
| | Estimate | ±SE | t value | p |
| (Intercept) | 5.77 | 0.11 | 53.18 | *** |
| siteGL | 0.18 | 0.09 | 2.01 | * |
| siteHW | 0.06 | 0.08 | 0.83 | - |
| siteJB | 0.10 | 0.09 | 1.10 | - |
| siteLU | 0.27 | 0.13 | 2.05 | * |
| sitePH | -0.08 | 0.08 | -1.02 | - |
| siteSK | 0.16 | 0.09 | 1.66 | - |
| siteST | 0.22 | 0.10 | 2.14 | * |
| siteW | 0.12 | 0.10 | 1.27 | - |
| age_l | -0.02 | 0.00 | -8.44 | *** |
| SMI | 0.07 | 0.01 | 13.09 | *** |
| Tnf(TG) | -0.20 | 0.09 | -2.18 | * |
| Tnf(TT) | -0.37 | 0.14 | -2.76 | ** |

| CD4+ T cells ~ age (linear) x IL13 + site + SMI | | | | |
|---|---|---|---|---|
| | Estimate | ±SE | t value | p |
| (Intercept) | 5.61 | 0.12 | 47.39 | *** |
| age_l | -0.02 | 0.00 | -7.75 | *** |
| IL13(TC) | 0.61 | 0.23 | 2.68 | ** |
| IL13(TT) | 0.90 | 0.31 | 2.90 | ** |
| siteGL | 0.36 | 0.10 | 3.46 | *** |
| siteHW | 0.25 | 0.09 | 2.64 | ** |
| siteJB | 0.28 | 0.10 | 2.69 | ** |
| siteLU | 0.25 | 0.12 | 2.03 | * |
| sitePH | 0.10 | 0.10 | 1.01 | - |
| siteSK | 0.31 | 0.11 | 2.90 | ** |
| siteST | 0.27 | 0.10 | 2.61 | ** |
| siteW | 0.25 | 0.11 | 2.27 | * |
| SMI | 0.07 | 0.01 | 12.44 | *** |
| age_l x IL13 (TC) | -0.03 | 0.02 | -1.98 | * |

| Tregs ~ site x age (linear) + SMI + IL6 | | | | |
|---|---|---|---|---|
| | Estimate | ±SE | t value | p |
| (Intercept) | 4.44 | 0.22 | 19.95 | *** |
| siteGL | 0.18 | 0.23 | 0.75 | - |
| siteHW | 0.00 | 0.20 | 0.02 | - |
| siteJB | 0.10 | 0.21 | 0.48 | - |
| siteLU | 0.36 | 0.39 | 0.92 | - |
| sitePH | -0.35 | 0.20 | -1.76 | - |
| siteSK | -0.72 | 0.29 | -2.53 | * |
| siteST | 0.02 | 0.22 | 0.09 | - |
| siteW | 0.28 | 0.23 | 1.20 | - |
| age_l | -0.02 | 0.01 | -2.26 | * |
| SMI | 0.08 | 0.01 | 10.60 | *** |
| IL6(CT) | -0.13 | 0.13 | -0.95 | - |
| IL6(TT) | 0.22 | 0.17 | 1.31 | - |
| siteGL x age_l | -0.01 | 0.02 | -0.51 | - |
| siteHW x age_ | 0.00 | 0.01 | 0.06 | - |
| siteJB x age_l | -0.02 | 0.01 | -1.33 | - |
| siteLU x age_l | -0.03 | 0.02 | -1.38 | - |
| sitePH x age_l | 0.00 | 0.01 | 0.37 | - |
| siteSK x age_l | 0.06 | 0.03 | 2.15 | * |
| siteST x age_l | 0.00 | 0.02 | -0.04 | - |
| siteW x age_l | -0.03 | 0.02 | -1.72 | - |

| Treg cells ~ site x age (linear) + SMI + IL13 | | | | |
|---|---|---|---|---|
| | Estimate | ±SE | t value | p |
| (Intercept) | 4.18 | 0.25 | 17.04 | *** |
| siteGL | 0.43 | 0.25 | 1.74 | - |
| siteHW | 0.26 | 0.21 | 1.22 | - |
| siteJB | 0.35 | 0.23 | 1.50 | - |
| siteLU | 0.66 | 0.41 | 1.61 | - |
| sitePH | -0.10 | 0.22 | -0.47 | - |
| siteSK | -0.30 | 0.28 | -1.05 | - |
| siteST | 0.23 | 0.24 | 0.95 | - |
| siteW | 0.62 | 0.25 | 2.50 | * |
| age_l | -0.02 | 0.01 | -1.62 | - |
| SMI | 0.08 | 0.01 | 10.53 | *** |
| IL13 (TC) | 0.32 | 0.16 | 2.02 | * |
| IL13 (TT) | 0.99 | 0.38 | 2.61 | ** |
| siteGL x age_l | -0.01 | 0.02 | -0.87 | - |
| siteHW x age_l | -0.01 | 0.01 | -0.42 | - |
| siteJB x age_l | -0.02 | 0.01 | -1.63 | - |
| siteLU x age_l | -0.04 | 0.02 | -1.78 | - |
| sitePH x age_l | 0.00 | 0.01 | -0.02 | - |
| siteSK x age_l | 0.04 | 0.03 | 1.38 | - |
| siteST x age_l | -0.01 | 0.02 | -0.38 | - |
| siteW x age_l | -0.04 | 0.02 | -2.45 | * |

| Macrophages ~ age (linear) x IL13 + SMI | | | | |
|---|---|---|---|---|
| | Estimate | ±SE | t value | p |
| (Intercept) | 4.06 | 0.16 | 24.70 | *** |
| age_l | -0.04 | 0.00 | -9.10 | *** |
| IL13 (TC) | 1.08 | 0.42 | 2.58 | * |
| IL13 (TT) | 1.08 | 0.55 | 1.94 | - |
| SMI | 0.10 | 0.01 | 9.34 | *** |
| age_l x IL13 (TC) | -0.07 | 0.03 | -2.23 | * |

| Macrophages ~ age (linear) + SMI + IL10 | | | | |
|---|---|---|---|---|
| | Estimate | ±SE | t value | p |
| (Intercept) | 4.08 | 0.18 | 22.41 | *** |
| age_l | -0.04 | 0.00 | -9.83 | *** |
| SMI | 0.10 | 0.01 | 9.80 | *** |
| IL10(CT) | -0.56 | 0.19 | -2.94 | ** |
| IL10(TT) | 0.00 | 0.11 | 0.00 | - |

| HGM ~ site x age (linear) + SMI + sex + IL13 | | | | |
|---|---|---|---|---|
| | Estimate | ±SE | t value | p |
| (Intercept) | 2.62 | 0.49 | 5.39 | *** |
| siteGL | 0.35 | 0.49 | 0.72 | - |
| siteHW | 0.50 | 0.43 | 1.18 | - |
| siteJB | 1.31 | 0.46 | 2.86 | ** |
| siteLU | 0.66 | 0.80 | 0.82 | - |
| sitePH | -0.56 | 0.44 | -1.27 | - |
| siteSK | -0.39 | 0.50 | -0.79 | - |
| siteST | 0.02 | 0.47 | 0.04 | - |
| siteW | 0.82 | 0.49 | 1.66 | - |
| age_l | 0.00 | 0.02 | 0.10 | - |
| SMI | 0.14 | 0.02 | 8.71 | *** |
| sex (M) | -0.25 | 0.09 | -2.87 | ** |
| IL13 (TC) | 0.61 | 0.32 | 1.93 | - |
| IL13 (TT) | 1.56 | 0.75 | 2.09 | * |
| siteGL x age_l | -0.06 | 0.03 | -2.05 | * |
| siteHW x age_l | -0.04 | 0.02 | -1.53 | - |
| siteJB x age_l | -0.10 | 0.03 | -3.58 | *** |
| siteLU x age_l | -0.07 | 0.04 | -1.72 | - |
| sitePH x age_l | -0.04 | 0.02 | -1.53 | - |
| siteSK x age_l | 0.01 | 0.03 | 0.33 | - |
| siteST x age_l | -0.03 | 0.03 | -0.94 | - |
| siteW x age_l | -0.08 | 0.03 | -2.53 | * |

| PMN ~ site x IL17f + age (classes) + SMI | | | | |
|---|---|---|---|---|
| | Estimate | ±SE | t value | p |
| (Intercept) | 2.55 | 0.73 | 3.50 | *** |
| siteGL | 1.48 | 0.79 | 1.88 | - |
| siteHW | 0.55 | 0.71 | 0.77 | - |
| siteJB | 0.87 | 0.78 | 1.13 | - |
| siteLU | -0.01 | 0.43 | -0.02 | - |
| sitePH | -0.53 | 0.72 | -0.74 | - |
| siteSK | 0.54 | 0.87 | 0.63 | - |
| siteST | 1.09 | 1.00 | 1.09 | - |
| siteW | 0.65 | 0.74 | 0.88 | - |
| IL17f(CT) | 1.15 | 0.75 | 1.52 | - |
| IL17f(TT) | 0.53 | 0.76 | 0.70 | - |
| age_cl(IM) | 0.91 | 0.11 | 8.06 | *** |
| age_cl(YO) | 0.38 | 0.11 | 3.48 | *** |
| SMI | 0.12 | 0.01 | 8.30 | *** |
| siteGL x IL17f(CT) | -2.87 | 0.85 | -3.39 | *** |
| siteJB x IL17f(CT) | -1.42 | 0.84 | -1.69 | - |
| siteLU x IL17f(CT) | -0.82 | 0.60 | -1.37 | - |
| sitePH x IL17f(CT) | -0.68 | 0.79 | -0.87 | - |
| siteSK x IL17f(CT) | -1.09 | 1.00 | -1.09 | - |
| siteW x IL17f(CT) | -1.52 | 0.83 | -1.83 | - |
| siteGL x IL17f(TT) | -1.35 | 0.89 | -1.51 | - |
| siteHW x IL17f(TT) | 0.13 | 1.04 | 0.13 | - |
| siteJB x IL17f(TT) | -0.86 | 0.85 | -1.01 | - |
| sitePH x IL17f(TT) | -0.27 | 0.92 | -0.29 | - |
| siteSK x IL17f(TT) | -1.11 | 0.93 | -1.19 | - |
| siteST x IL17f(TT) | -1.40 | 1.05 | -1.33 | - |
| siteW x IL17f(TT) | -0.39 | 0.85 | -0.46 | - |

| PMN ~ site x IL17a_U + age (classes) + SMI | | | | |
|---|---|---|---|---|
| | Estimate | ±SE | t value | p |
| (Intercept) | 3.44 | 0.31 | 11.25 | *** |
| siteGL | -1.65 | 0.37 | -4.40 | *** |
| siteHW | -0.45 | 0.43 | -1.06 | - |
| siteJB | -0.22 | 0.24 | -0.91 | - |
| siteLU | -0.32 | 0.32 | -1.03 | - |
| sitePH | -1.04 | 0.29 | -3.60 | *** |
| siteSK | -0.72 | 0.24 | -2.98 | ** |
| siteST | -0.59 | 0.28 | -2.13 | * |
| siteW | -0.27 | 0.25 | -1.08 | - |
| IL17a_U(GA) | -0.27 | 0.27 | -1.01 | - |
| IL17a_U(GG) | 0.21 | 0.37 | 0.56 | - |
| age_cl(IM) | 0.87 | 0.11 | 7.71 | *** |
| age_cl(YO) | 0.33 | 0.11 | 3.05 | ** |
| SMI | 0.12 | 0.01 | 7.87 | *** |
| siteGL x IL17a_U(GA) | 1.66 | 0.45 | 3.70 | *** |
| siteLU x IL17a_U(GA) | -0.63 | 0.80 | -0.79 | - |
| siteGL x IL17a_U(GG) | 0.56 | 0.87 | 0.65 | - |
| siteJB x IL17a_U(GG) | -0.20 | 0.82 | -0.25 | - |
| sitePH x IL17a_U(GG) | -0.39 | 0.47 | -0.83 | - |
| siteSK x IL17a_U(GG) | -0.07 | 0.83 | -0.08 | - |

| Neutrophils ~ site x age (linear) + SMI + IL6 | | | | |
|---|---|---|---|---|
| | Estimate | ±SE | t value | p |
| (Intercept) | 4.00 | 0.30 | 13.42 | *** |
| siteGL | 0.27 | 0.30 | 0.91 | |
| siteHW | 0.17 | 0.26 | 0.65 | |
| siteJB | 0.33 | 0.29 | 1.14 | |
| siteLU | 1.24 | 0.95 | 1.31 | |
| sitePH | -0.34 | 0.29 | -1.16 | |
| siteSK | -0.48 | 0.32 | -1.47 | |
| siteST | -0.04 | 0.31 | -0.14 | |
| siteW | 0.74 | 0.30 | 2.45 | * |
| age_l | 0.01 | 0.01 | 0.55 | |
| SMI | 0.10 | 0.01 | 9.11 | *** |
| IL6(CT) | -0.43 | 0.17 | -2.49 | * |
| IL6(TT) | -0.13 | 0.22 | -0.60 | |
| siteGL x age_l | -0.04 | 0.02 | -2.00 | * |
| siteHW x age_l | -0.03 | 0.02 | -2.24 | * |
| siteJB x age_l | -0.05 | 0.02 | -3.00 | ** |
| siteLU x age_l | -0.09 | 0.05 | -1.98 | * |
| sitePH x age_l | -0.03 | 0.02 | -2.11 | * |
| siteSK x age_l | -0.01 | 0.02 | -0.73 | |
| siteST x age_l | -0.02 | 0.03 | -0.60 | |
| siteW x age_l | -0.07 | 0.02 | -3.50 | *** |

| Neutrophils ~ site x age (linear) + SMI + IL10 | | | | |
|---|---|---|---|---|
| | Estimate | ±SE | t value | p |
| (Intercept) | 3.45 | 0.35 | 9.84 | *** |
| siteGL | 0.74 | 0.29 | 2.54 | * |
| siteHW | 0.32 | 0.23 | 1.36 | - |
| siteJB | 0.52 | 0.26 | 2.03 | * |
| siteLU | 1.37 | 0.93 | 1.47 | - |
| sitePH | -0.18 | 0.26 | -0.69 | - |
| siteSK | 0.10 | 0.35 | 0.29 | - |
| siteST | 0.18 | 0.31 | 0.57 | - |
| siteW | 0.65 | 0.29 | 2.26 | * |
| age_l | 0.00 | 0.01 | 0.13 | - |
| SMI | 0.09 | 0.01 | 9.11 | *** |
| IL10(CT) | -0.17 | 0.25 | -0.69 | - |
| IL10(TT) | 0.45 | 0.22 | 2.09 | * |
| siteGL x age_l | -0.04 | 0.02 | -1.99 | * |
| siteHW x age_l | -0.03 | 0.02 | -1.89 | - |
| siteJB x age_l | -0.05 | 0.02 | -2.94 | ** |
| siteLU x age_l | -0.08 | 0.04 | -1.87 | - |
| sitePH x age_l | -0.03 | 0.02 | -1.80 | - |
| siteSK x age_l | -0.01 | 0.02 | -0.46 | - |
| siteST x age_l | 0.00 | 0.03 | -0.14 | - |
| siteW x age_l | -0.07 | 0.02 | -3.44 | *** |

| MDSC ~ site x age (linear) + SMI + H2Aa | | | | |
|---|---|---|---|---|
| | Estimate | ±SE | t value | p |
| (Intercept) | 3.48 | 0.34 | 10.23 | *** |
| siteGL | 0.40 | 0.33 | 1.24 | - |
| siteHW | 0.23 | 0.23 | 0.98 | - |
| siteJB | 0.47 | 0.25 | 1.84 | - |
| siteLU | 2.52 | 0.91 | 2.75 | ** |
| sitePH | -0.06 | 0.28 | -0.23 | - |
| siteSK | -0.61 | 0.29 | -2.13 | * |
| siteST | -0.27 | 0.30 | -0.92 | - |
| siteW | 0.99 | 0.30 | 3.31 | ** |
| age_l | 0.00 | 0.01 | 0.35 | - |
| SMI | 0.09 | 0.01 | 8.86 | *** |
| H2Aa(AG) | -0.39 | 0.19 | -2.10 | * |
| H2Aa(GG) | -0.07 | 0.21 | -0.34 | - |
| siteGL x age_l | -0.05 | 0.02 | -2.37 | * |
| siteHW x age_l | -0.03 | 0.01 | -1.82 | - |
| siteJB x age_l | -0.06 | 0.02 | -3.50 | *** |
| siteLU x age_l | -0.13 | 0.04 | -2.97 | ** |
| sitePH x age_l | -0.04 | 0.02 | -2.29 | * |
| siteSK x age_l | 0.03 | 0.02 | 1.44 | - |
| siteST x age_l | -0.03 | 0.03 | -0.94 | - |
| siteW x age_l | -0.09 | 0.02 | -4.26 | *** |

| MDSC ~ site x age (linear) + SMI + IL6 | | | | |
|---|---|---|---|---|
| | Estimate | ±SE | t value | p |
| (Intercept) | 3.53 | 0.29 | 12.22 | *** |
| siteGL | 0.19 | 0.29 | 0.63 | - |
| siteHW | 0.07 | 0.25 | 0.27 | - |
| siteJB | 0.27 | 0.28 | 0.97 | - |
| siteLU | 2.37 | 0.92 | 2.58 | * |
| sitePH | -0.43 | 0.28 | -1.53 | - |
| siteSK | -0.77 | 0.31 | -2.44 | * |
| siteST | -0.21 | 0.30 | -0.71 | - |
| siteW | 0.86 | 0.29 | 2.92 | ** |
| age_l | 0.01 | 0.01 | 0.86 | - |
| SMI | 0.10 | 0.01 | 9.21 | *** |
| IL6(CT) | -0.52 | 0.17 | -3.10 | ** |
| IL6(TT) | -0.20 | 0.21 | -0.95 | - |
| siteGL x age_l | -0.05 | 0.02 | -2.61 | ** |
| siteHW x age_l | -0.03 | 0.02 | -2.31 | * |
| siteJB x age_l | -0.07 | 0.02 | -3.74 | *** |
| siteLU x age_l | -0.14 | 0.04 | -3.14 | ** |
| sitePH x age_l | -0.04 | 0.02 | -2.58 | * |
| siteSK x age_l | 0.02 | 0.02 | 1.00 | - |
| siteST x age_l | -0.05 | 0.03 | -1.57 | - |
| siteW x age_l | -0.08 | 0.02 | -3.85 | *** |

| MDSC ~ site x age (linear) + SMI + IL10 | | | | |
|---|---|---|---|---|
| | Estimate | ±SE | t value | p |
| (Intercept) | 3.06 | 0.34 | 8.92 | *** |
| siteGL | 0.67 | 0.28 | 2.35 | * |
| siteHW | 0.26 | 0.23 | 1.15 | - |
| siteJB | 0.50 | 0.25 | 1.98 | * |
| siteLU | 2.55 | 0.91 | 2.81 | ** |
| sitePH | -0.23 | 0.26 | -0.89 | - |
| siteSK | -0.29 | 0.35 | -0.84 | - |
| siteST | -0.04 | 0.30 | -0.12 | - |
| siteW | 0.73 | 0.28 | 2.62 | ** |
| age_l | 0.01 | 0.01 | 0.38 | - |
| SMI | 0.09 | 0.01 | 9.13 | *** |
| IL10(CT) | -0.26 | 0.24 | -1.08 | - |
| IL10(TT) | 0.32 | 0.21 | 1.49 | - |
| siteGL x age_l | -0.05 | 0.02 | -2.56 | * |
| siteHW x age_l | -0.03 | 0.01 | -1.88 | - |
| siteJB x age_l | -0.06 | 0.02 | -3.55 | *** |
| siteLU x age_l | -0.13 | 0.04 | -3.00 | ** |
| sitePH x age_l | -0.03 | 0.02 | -2.21 | * |
| siteSK x age_l | 0.03 | 0.02 | 1.45 | - |
| siteST x age_l | -0.03 | 0.03 | -0.92 | - |
| siteW x age_l | -0.07 | 0.02 | -3.69 | *** |

| MDSC ~ site x age (linear) + SMI + IL17a_N | | | | |
|---|---|---|---|---|
| | Estimate | ±SE | t value | p |
| (Intercept) | 2.71 | 0.40 | 6.72 | *** |
| siteGL | 0.37 | 0.27 | 1.35 | - |
| siteHW | 0.26 | 0.23 | 1.11 | - |
| siteJB | 0.48 | 0.25 | 1.90 | - |
| siteLU | 2.53 | 0.92 | 2.76 | ** |
| sitePH | -0.23 | 0.26 | -0.87 | - |
| siteSK | 0.10 | 0.43 | 0.23 | - |
| siteST | -0.24 | 0.30 | -0.82 | - |
| siteW | 0.73 | 0.28 | 2.56 | * |
| age_l | 0.00 | 0.01 | 0.36 | - |
| SMI | 0.09 | 0.01 | 8.77 | *** |
| IL17a_N(AG) | 0.92 | 0.36 | 2.55 | * |
| IL17a_N(GG) | 0.69 | 0.31 | 2.21 | * |
| siteGL x age_l | -0.04 | 0.02 | -2.23 | * |
| siteHW x age_l | -0.03 | 0.01 | -1.82 | - |
| siteJB x age_l | -0.06 | 0.02 | -3.49 | *** |
| siteLU x age_l | -0.13 | 0.04 | -2.96 | ** |
| sitePH x age_l | -0.03 | 0.02 | -2.18 | * |
| siteSK x age_l | 0.02 | 0.02 | 0.87 | - |
| siteST x age_l | -0.03 | 0.03 | -0.95 | - |
| siteW x age_l | -0.07 | 0.02 | -3.60 | *** |

#### Macroparasites

| mites ~ H2Aa + site + age (classes) + SMI | | | | |
|---|---|---|---|---|
| | Estimate | ±SE | t value | p |
| 0\|50 | 0.33 | 0.98 | 0.33 | - |
| 50\|100 | 0.53 | 0.98 | 0.54 | - |
| 100\|200 | 1.11 | 0.98 | 1.14 | - |
| 200\|300 | 2.24 | 0.98 | 2.28 | * |
| 300\|400 | 2.62 | 0.99 | 2.66 | ** |
| 400\|500 | 2.75 | 0.99 | 2.79 | ** |
| 500\|1000 | 3.63 | 0.99 | 3.66 | *** |
| 1000\|1500 | 4.78 | 1.01 | 4.75 | *** |
| siteGL | 1.32 | 0.79 | 1.67 | - |
| siteHW | 1.67 | 0.37 | 4.55 | *** |
| siteJB | -0.08 | 0.48 | -0.18 | - |
| siteLU | -1.57 | 0.65 | -2.41 | * |
| sitePH | 1.32 | 0.49 | 2.71 | ** |
| siteSK | -2.65 | 0.82 | -3.22 | ** |
| siteST | 1.68 | 0.48 | 3.52 | *** |
| siteWF+WT | 1.16 | 0.54 | 2.16 | * |
| age_cl.L | -1.01 | 0.19 | -5.24 | *** |
| age_cl.Q | 0.30 | 0.17 | 1.75 | - |
| H2Aa(AG) | 1.91 | 0.76 | 2.50 | * |
| H2Aa(GG) | 1.80 | 0.84 | 2.15 | * |
| SMI | -0.10 | 0.04 | -2.79 | ** |

| mites ~ site + age (classes) + TLR9 + SMI | | | | |
|---|---|---|---|---|
| 0\|L50 | -1.53 | 0.57 | -2.69 | * |
| L50\|100 | -1.29 | 0.57 | -2.27 | - |
| 100\|200 | -0.67 | 0.56 | -1.19 | - |
| 200\|300 | 0.46 | 0.56 | 0.81 | - |
| 300\|400 | 0.85 | 0.56 | 1.51 | - |
| 400\|500 | 0.98 | 0.56 | 1.74 | - |
| 500\|1000 | 1.88 | 0.57 | 3.30 | *** |
| 1000\|1500 | 3.04 | 0.59 | 5.12 | *** |
| siteGL | 0.05 | 0.50 | 0.10 | - |
| siteHW | 1.70 | 0.37 | 4.60 | *** |
| siteJB | 0.84 | 0.53 | 1.57 | - |
| siteLU | -1.60 | 0.66 | -2.44 | * |
| sitePH | 1.35 | 0.45 | 3.00 | ** |
| siteSK | -2.70 | 0.82 | -3.27 | ** |
| siteST | 1.70 | 0.47 | 3.58 | *** |
| siteW | 1.21 | 0.49 | 2.48 | * |
| age_cl.L | -1.07 | 0.19 | -5.66 | *** |
| age_cl.Q | 0.27 | 0.17 | 1.60 | - |
| Tlr9(TC) | -2.50 | 0.78 | -3.19 | ** |
| SMI | -0.10 | 0.03 | -2.85 | ** |

| mites ~ site + age (classes) + TLR5 + SMI | | | | |
|---|---|---|---|---|
| 0\|L50 | 0.65 | 1.23 | 0.52 | - |
| L50\|100 | 0.88 | 1.23 | 0.71 | - |
| 100\|200 | 1.49 | 1.23 | 1.21 | - |
| 200\|300 | 2.60 | 1.24 | 2.11 | * |
| 300\|400 | 2.99 | 1.24 | 2.42 | * |
| 400\|500 | 3.12 | 1.24 | 2.52 | * |
| 500\|1000 | 4.01 | 1.24 | 3.23 | ** |
| 1000\|1500 | 5.16 | 1.25 | 4.12 | *** |
| siteGL | -0.02 | 0.53 | -0.04 | - |
| siteHW | 1.61 | 0.42 | 3.84 | *** |
| siteJB | -0.16 | 0.52 | -0.31 | - |
| siteLU | -1.65 | 0.69 | -2.41 | * |
| sitePH | 1.27 | 0.49 | 2.59 | ** |
| siteSK | -2.75 | 0.85 | -3.24 | ** |
| siteST | 1.66 | 0.49 | 3.39 | *** |
| siteW | 1.13 | 0.53 | 2.16 | * |
| age_cl.L | -1.03 | 0.19 | -5.48 | *** |
| age_cl.Q | 0.29 | 0.17 | 1.70 | - |
| Tlr5(AG) | 2.67 | 1.23 | 2.17 | * |
| Tlr5(GG) | 2.22 | 1.16 | 1.91 | - |
| SMI | -0.10 | 0.03 | -2.84 | ** |

| worms ~ H2Ab x site + age (linear) + SMI | | | | |
|---|---|---|---|---|
| | Estimate | ±SE | t value | p |
| (Intercept) | 0.69 | 0.22 | 3.15 | ** |
| siteGL | 0.36 | 0.39 | 0.91 | - |
| siteHW | 0.13 | 0.21 | 0.64 | - |
| siteJB | -0.06 | 0.20 | -0.29 | - |
| siteLU | -1.22 | 0.23 | -5.23 | *** |
| sitePH | 0.66 | 0.41 | 1.61 | - |
| siteSK | -1.12 | 0.40 | -2.83 | ** |
| siteST | 0.46 | 0.32 | 1.45 | - |
| siteWF+WT | 0.04 | 0.24 | 0.16 | - |
| H2Ab(AG) | -0.10 | 0.24 | -0.42 | - |
| H2Ab(GG) | -0.12 | 0.41 | -0.29 | - |
| age_l | 0.01 | 0.00 | 2.46 | * |
| SMI | 0.03 | 0.01 | 2.37 | * |
| siteGL x H2Ab(AG) | 0.33 | 0.78 | 0.42 | - |
| siteHW x H2Ab(AG) | 0.21 | 0.28 | 0.74 | - |
| siteJB x H2Ab(AG) | -1.02 | 0.70 | -1.47 | - |
| siteLU x H2Ab(AG) | 0.33 | 0.54 | 0.61 | - |
| sitePH x H2Ab(AG) | 0.05 | 0.48 | 0.10 | - |
| siteST x H2Ab(AG) | -0.37 | 0.40 | -0.93 | - |
| siteWF+WT x H2Ab(AG) | 0.85 | 0.38 | 2.21 | * |
| siteHW x H2Ab(GG) | 0.32 | 0.43 | 0.73 | - |
| siteJB x H2Ab(GG) | -1.01 | 0.77 | -1.31 | - |
| siteST x H2Ab(GG) | -1.21 | 0.55 | -2.19 | * |
| siteWF+WT x H2Ab(GG) | 0.89 | 0.63 | 1.41 | |

| worms ~ site + age (linear) + SMI + IL6 | | | | |
|---|---|---|---|---|
| (Intercept) | 0.66 | 0.19 | 3.40 | *** |
| siteGL | 0.26 | 0.18 | 1.49 | - |
| siteHW | 0.26 | 0.14 | 1.79 | - |
| siteJB | -0.13 | 0.17 | -0.77 | - |
| siteLU | -1.18 | 0.20 | -5.77 | *** |
| sitePH | 0.61 | 0.18 | 3.47 | *** |
| siteSK | -1.24 | 0.18 | -6.71 | *** |
| siteST | -0.06 | 0.18 | -0.35 | - |
| siteW | 0.46 | 0.20 | 2.34 | * |
| age_l | 0.01 | 0.00 | 2.73 | ** |
| SMI | 0.03 | 0.01 | 2.60 | ** |
| IL6(CT) | -0.37 | 0.18 | -2.07 | * |
| IL6(TT) | 0.28 | 0.24 | 1.19 | - |

#### Microparasites

| norovirus ~ site + H2Eb x age (classes) | | | | |
|---|---|---|---|---|
| | Estimate | ±SE | t value | p |
| 0\|1 | 1.90 | 0.64 | 2.49 | ** |
| 1\|2 | 4.41 | 0.69 | 6.35 | *** |
| 2\|3 | 6.17 | 0.77 | 7.99 | *** |
| siteGL | 3.35 | 0.60 | 5.58 | *** |
| siteHW | 1.25 | 0.52 | 2.41 | * |
| siteJB | 1.11 | 0.58 | 1.89 | - |
| siteLU | 5.17 | 0.75 | 6.85 | *** |
| sitePH | 0.88 | 0.59 | 1.49 | - |
| siteSK | 0.82 | 0.63 | 1.30 | - |
| siteST | 1.34 | 0.67 | 1.99 | * |
| siteW | 2.71 | 0.64 | 4.25 | *** |
| H2Eb(AG) | -0.64 | 0.57 | -1.13 | - |
| H2Eb(GG) | 0.42 | 0.46 | 0.92 | - |
| age_cl.L | 0.35 | 0.76 | 0.46 | - |
| age_cl.Q | -0.25 | 0.59 | -0.42 | - |
| H2Eb(AG) x age_cl.L | -1.32 | 1.10 | -1.20 | - |
| H2Eb(GG) x age_cl.L | 1.02 | 0.80 | 1.27 | - |
| H2Eb(AG) x age_cl.Q | -0.59 | 0.81 | -0.73 | - |
| H2Eb(GG) x age_cl.Q | 0.09 | 0.64 | 0.13 | - |

| norovirus ~ site + IL10 x age (classes) | | | | |
|---|---|---|---|---|
| | Estimate | ±SE | t value | p |
| 0\|1 | 4.81 | 31.17 | 0.15 | - |
| 1\|2 | 7.28 | 31.18 | 0.23 | - |
| 2\|3 | 9.09 | 31.18 | 0.29 | - |
| siteGL | 3.10 | 0.62 | 5.02 | *** |
| siteHW | 0.90 | 0.48 | 1.86 | - |
| siteJB | 1.05 | 0.58 | 1.82 | - |
| siteLU | 5.29 | 0.76 | 7.00 | *** |
| sitePH | 0.86 | 0.58 | 1.49 | - |
| siteSK | 0.46 | 1.29 | 0.35 | - |
| siteST | 0.87 | 0.66 | 1.32 | - |
| siteW | 2.63 | 0.63 | 4.15 | *** |
| IL10(CT) | 11.27 | 40.07 | 0.28 | - |
| IL10(TT) | 3.39 | 31.18 | 0.11 | - |
| age_cl.L | 9.35 | 66.11 | 0.14 | - |
| age_cl.Q | -3.25 | 38.18 | -0.09 | - |
| IL10(CT) x age_cl.L | 7.45 | 47.22 | 0.16 | - |
| IL10(TT) x age_cl.L | -8.54 | 66.11 | -0.13 | - |
| IL10(CT) x age_cl.Q | 11.46 | 49.08 | 0.23 | - |
| IL10(TT) x age_cl.Q | 3.12 | 38.18 | 0.08 | - |

| parvovirus ~ site + age (classes) + H2Eb | | | | |
|---|---|---|---|---|
| | Estimate | ±SE | t value | p |
| 0\|1 | -1.25 | 0.51 | -2.43 | * |
| 1\|2 | 0.18 | 0.50 | 0.35 | - |
| 2\|3 | 0.97 | 0.51 | 1.92 | - |
| 3\|4 | 3.13 | 0.54 | 5.84 | *** |
| siteGL | -0.75 | 0.49 | -1.53 | - |
| siteHW | 0.75 | 0.40 | 1.87 | - |
| siteJB | -0.29 | 0.47 | -0.62 | - |
| siteLU | -0.32 | 0.58 | -0.54 | - |
| sitePH | 1.83 | 0.48 | 3.82 | *** |
| siteSK | 0.18 | 0.52 | 0.34 | - |
| siteST | -0.71 | 0.56 | -1.27 | - |
| siteW | 0.30 | 0.48 | 0.61 | - |
| age_cl.L | 1.40 | 0.20 | 7.15 | *** |
| age_cl.Q | -0.19 | 0.18 | -1.10 | - |
| H2Eb(AG) | 1.02 | 0.40 | 2.51 | * |
| H2Eb(GG) | 0.56 | 0.38 | 1.48 | - |

| minute virus ~ site + age (classes) + H2Ab | | | | |
|---|---|---|---|---|
| | Estimate | ±SE | t value | p |
| 0\|1 | 1.59 | 0.53 | 3.03 | ** |
| 1\|2 | 2.99 | 0.55 | 5.40 | *** |
| 2\|3 | 3.82 | 0.59 | 6.53 | *** |
| siteGL | -1.74 | 0.84 | -2.06 | * |
| siteHW | -0.04 | 0.55 | -0.07 | - |
| siteJB | 0.26 | 0.67 | 0.39 | - |
| siteLU | 0.24 | 0.81 | 0.30 | - |
| sitePH | 2.23 | 0.66 | 3.39 | *** |
| siteSK | 0.39 | 0.72 | 0.54 | - |
| siteST | 0.05 | 0.76 | 0.07 | - |
| siteW | 2.45 | 0.66 | 3.69 | *** |
| age_cl.L | 1.27 | 0.26 | 4.88 | *** |
| age_cl.Q | 0.21 | 0.24 | 0.88 | - |
| H2Ab(AG) | -0.25 | 0.45 | -0.55 | - |
| H2Ab(GG) | 0.76 | 0.45 | 1.69 | - |

| minute virus ~ site + age (classes) + IL1a | | | | |
|---|---|---|---|---|
| | Estimate | ±SE | t value | p |
| 0\|1 | 2.56 | 0.66 | 3.86 | *** |
| 1\|2 | 3.94 | 0.69 | 5.73 | *** |
| 2\|3 | 4.73 | 0.71 | 6.62 | *** |
| siteGL | -0.82 | 0.79 | -1.04 | - |
| siteHW | 0.42 | 0.53 | 0.79 | - |
| siteJB | 0.48 | 0.66 | 0.74 | - |
| siteLU | 0.46 | 0.80 | 0.57 | - |
| sitePH | 3.44 | 0.76 | 4.52 | *** |
| siteSK | 1.96 | 0.78 | 2.51 | * |
| siteST | 0.21 | 0.75 | 0.29 | - |
| siteW | 2.83 | 0.71 | 4.00 | *** |
| age_cl.L | 1.23 | 0.25 | 4.87 | *** |
| age_cl.Q | 0.22 | 0.23 | 0.97 | - |
| IL1a(TC) | 1.10 | 0.46 | 2.37 | * |
| IL1a(TT) | 0.75 | 0.52 | 1.43 | - |

| MHV ~ site + age (classes) + TNF | | | | |
|---|---|---|---|---|
| | Estimate | ±SE | t value | p |
| 0\|1 | -1.88 | 0.44 | -4.31 | *** |
| 1\|2 | -0.04 | 0.41 | -0.09 | - |
| 2\|3 | 1.60 | 0.43 | 3.74 | *** |
| siteGL | -4.58 | 0.76 | -6.00 | *** |
| siteHW | -2.86 | 0.47 | -6.11 | *** |
| siteJB | -4.13 | 0.70 | -5.89 | *** |
| siteLU | 1.51 | 0.98 | 1.55 | - |
| sitePH | -2.73 | 0.58 | -4.70 | *** |
| siteSK | -0.89 | 0.53 | -1.69 | - |
| siteST | -2.56 | 0.71 | -3.62 | *** |
| siteW | -2.52 | 0.64 | -3.95 | *** |
| age_cl.L | 1.78 | 0.28 | 6.40 | *** |
| age_cl.Q | -0.67 | 0.22 | -2.98 | ** |
| Tnf (TG) | -1.80 | 0.89 | -2.03 | * |
| Tnf (TT) | -1.31 | 1.21 | -1.08 | - |

| MHV ~ site x age (classes) + IL6 | | | | |
|---|---|---|---|---|
| | Estimate | ±SE | t value | p |
| 0\|1 | -3.00 | 0.54 | -5.54 | *** |
| 1\|2 | -0.76 | 0.49 | 1.53 | - |
| 2\|3 | 1.11 | 0.52 | 2.14 | * |
| siteGL | -7.45 | 14.31 | -0.52 | - |
| siteHW | -3.97 | 0.58 | -6.88 | *** |
| siteJB | -7.45 | 18.55 | -0.40 | - |
| siteLU | 0.12 | 0.79 | 0.16 | - |
| sitePH | -9.27 | 0.69 | -13.37 | *** |
| siteSK | -2.10 | 0.67 | -3.13 | ** |
| siteST | -3.76 | 0.80 | -4.68 | *** |
| siteW | -5.62 | 76.52 | -0.07 | - |
| age_cl.L | -0.16 | 0.76 | -0.21 | - |
| age_cl.Q | -2.34 | 0.71 | -3.28 | ** |
| IL6(CT) | -2.51 | 0.77 | -3.26 | ** |
| IL6(TT) | -1.41 | 0.98 | -1.44 | - |
| siteGL x age_cl.L | 5.27 | 30.34 | 0.17 | - |
| siteHW x age_cl.L | 1.63 | 0.87 | 1.86 | - |
| siteJB x age_cl.L | 7.05 | 39.33 | 0.18 | - |
| siteLU x age_cl.L | 1.64 | 1.10 | 1.49 | - |
| sitePH x age_cl.L | 14.96 | 0.50 | 30.11 | *** |
| siteSK x age_cl.L | 3.43 | 1.16 | 2.96 | ** |
| siteST x age_cl.L | 1.53 | 1.41 | 1.08 | - |
| siteW x age_cl.L | 9.92 | 162.30 | 0.06 | - |
| siteGL x age_cl.Q | -1.71 | 17.54 | -0.10 | - |
| siteHW x age_cl.Q | 1.59 | 0.78 | 2.04 | * |
| siteJB x age_cl.Q | -0.16 | 22.73 | -0.01 | - |
| sitePH x age_cl.Q | -2.28 | 1.01 | -2.26 | * |
| siteSK x age_cl.Q | 1.34 | 1.01 | 1.33 | - |
| siteST x age_cl.Q | 1.38 | 1.16 | 1.19 | - |
| siteW x age_cl.Q | -2.94 | 93.71 | -0.03 | - |

| MHV ~ site + age (classes) + H2Aa | | | | |
|---|---|---|---|---|
| | Estimate | ±SE | t value | p |
| 0\|1 | -1.90 | 1.88 | -1.01 | - |
| 1\|2 | -0.02 | 1.87 | -0.01 | - |
| 2\|3 | 1.61 | 1.88 | 0.86 | - |
| siteGL | -4.47 | 1.89 | -2.37 | * |
| siteHW | -2.84 | 0.46 | -6.19 | *** |
| siteJB | -4.12 | 0.69 | -5.93 | *** |
| siteLU | 0.14 | 0.63 | 0.22 | - |
| sitePH | -1.81 | 0.62 | -2.94 | ** |
| siteSK | -0.85 | 0.52 | -1.63 | - |
| siteST | -2.48 | 0.69 | -3.59 | *** |
| siteW | -2.66 | 0.69 | -3.86 | *** |
| age_cl.L | 1.77 | 0.28 | 6.38 | *** |
| age_cl.Q | -0.74 | 0.23 | -3.25 | ** |
| H2Aa(AG) | -2.64 | 1.97 | -1.34 | - |
| H2Aa(GG) | -0.04 | 1.84 | -0.02 | - |

| sendai virus ~ site + TLR9 x age (classes) | | | | |
|---|---|---|---|---|
| | Estimate | ±SE | t value | p |
| 0\|1 | 0.95 | 0.41 | 2.29 | * |
| 1\|2 | 4.22 | 0.48 | 8.75 | *** |
| 2\|3 | 6.66 | 0.83 | 8.03 | *** |
| siteGL | 0.34 | 0.56 | 0.60 | - |
| siteHW | 1.57 | 0.45 | 3.48 | *** |
| siteJB | 1.57 | 0.64 | 2.46 | * |
| siteLU | 1.33 | 0.69 | 1.93 | - |
| sitePH | 0.94 | 0.55 | 1.71 | - |
| siteSK | 1.50 | 0.57 | 2.63 | ** |
| siteST | 1.63 | 0.59 | 2.75 | ** |
| siteW | 1.42 | 0.62 | 2.29 | * |
| Tlr9(TC) | -0.43 | 0.77 | -0.56 | - |
| age_cl.L | 0.55 | 0.21 | 2.65 | ** |
| age_cl.Q | -0.60 | 0.20 | -3.08 | - |
| Tlr9(TC) x age_cl.L | 2.34 | 1.13 | 2.08 | * |
| Tlr9(TC) x age_cl.Q | 1.73 | 0.97 | 1.78 | - |

| sendai virus ~ site + age (classes) + IL6 | | | | |
|---|---|---|---|---|
| | Estimate | ±SE | t value | p |
| 0\|1 | 1.58 | 0.53 | 2.94 | * |
| 1\|2 | 4.79 | 0.59 | 8.06 | *** |
| 2\|3 | 7.20 | 0.89 | 8.03 | *** |
| siteGL | 0.95 | 0.66 | 1.44 | - |
| siteHW | 2.19 | 0.57 | 3.86 | *** |
| siteJB | 1.89 | 0.65 | 2.90 | ** |
| siteLU | 1.86 | 0.77 | 2.42 | * |
| sitePH | 1.59 | 0.65 | 2.45 | * |
| siteSK | 2.17 | 0.67 | 3.25 | ** |
| siteST | 2.04 | 0.64 | 3.18 | ** |
| siteW | 0.57 | 0.72 | 0.79 | - |
| age_cl.L | 0.63 | 0.21 | 3.03 | ** |
| age_cl.Q | -0.44 | 0.19 | -2.28 | * |
| IL6(CT) | 1.27 | 0.65 | 1.94 | - |
| IL6(TT) | 1.92 | 0.88 | 2.20 | * |

| sendai virus ~ site x Tnf + age (classes) | | | | |
|---|---|---|---|---|
| | Estimate | ±SE | t value | p |
| 0\|1 | 1.06 | 0.43 | 2.47 | * |
| 1\|2 | 4.36 | 0.50 | 8.77 | *** |
| 2\|3 | 6.79 | 0.84 | 8.10 | *** |
| siteGL | 0.41 | 0.58 | 0.72 | - |
| siteHW | 1.71 | 0.47 | 3.66 | *** |
| siteJB | 1.34 | 0.56 | 2.36 | * |
| siteLU | 2.37 | 1.44 | 1.64 | - |
| sitePH | 1.06 | 0.56 | 1.89 | - |
| siteSK | 1.58 | 0.58 | 2.70 | ** |
| siteST | 1.50 | 0.67 | 2.22 | * |
| siteW | 2.33 | 0.69 | 3.35 | *** |
| Tnf(TG) | 2.34 | 1.98 | 1.18 | - |
| Tnf(TT) | -18.03 | 0.96 | -18.80 | *** |
| age_cl.L | 0.74 | 0.21 | 3.46 | *** |
| age_cl.Q | -0.45 | 0.19 | -2.34 | * |
| siteLU x Tnf(TG) | -4.43 | 2.52 | -1.76 | - |
| siteS(TT)nf(TG) | -0.78 | 2.18 | -0.36 | - |
| siteW x Tnf(TG) | -15.12 | 174.00 | -0.08 | - |
| siteLU x Tnf(TT) | 18.56 | 0.96 | 19.35 | *** |
| siteS(TT)nf(TT) | 1.75 | 0.00 | 200.84 | *** |

| corona virus ~ site + IL17a_N x age (linear) | | | | |
|---|---|---|---|---|
| | Estimate | ±SE | t value | p |
| 0\|1 | 118.55 | 1.66 | 72.13 | *** |
| 1\|2 | 121.85 | 1.67 | 73.09 | *** |
| siteGL | -4.30 | 0.72 | -5.96 | *** |
| siteHW | -5.10 | 0.62 | -8.22 | *** |
| siteJB | -4.12 | 0.71 | -5.76 | *** |
| siteLU | -0.90 | 0.67 | -1.33 | - |
| sitePH | -6.15 | 1.16 | -5.30 | *** |
| siteSK | -22.83 | 3.26 | -6.99 | *** |
| siteST | -3.94 | 0.76 | -5.20 | *** |
| siteW | -5.69 | 1.16 | -4.91 | *** |
| IL17a_N(AG) | 68.24 | 0.01 | 6734.50 | *** |
| IL17a_N(GG) | 121.32 | 1.66 | 73.19 | *** |
| age_l | 6.15 | 39.97 | 0.15 | - |
| IL17a_N(AG) x age_l | -2.26 | 0.14 | -16.65 | *** |
| IL17a_N(GG) x age_l | -6.08 | 39.97 | -0.15 | - |

| M. pulmonis ~ site + age (classes) + IL6 | | | | |
|---|---|---|---|---|
| | Estimate | ±SE | t value | p |
| 0\|1 | 1.56 | 0.53 | 2.90 | ** |
| 1\|2 | 4.89 | 0.60 | 8.21 | *** |
| 2\|3 | 639.36 | 0.60 | 1073.58 | *** |
| siteGL | 0.75 | 0.67 | 1.11 | - |
| siteHW | 2.08 | 0.57 | 3.67 | *** |
| siteJB | 1.93 | 0.65 | 2.96 | ** |
| siteLU | 2.58 | 0.78 | 3.33 | *** |
| sitePH | 1.93 | 0.65 | 2.97 | ** |
| siteSK | 2.25 | 0.67 | 3.34 | *** |
| siteST | 2.61 | 0.65 | 4.03 | *** |
| siteW | 0.49 | 0.70 | 0.70 | - |
| age_cl.L | 0.71 | 0.21 | 3.40 | *** |
| age_cl.Q | -0.30 | 0.19 | -1.57 | - |
| IL6(CT) | 0.88 | 0.65 | 1.35 | - |
| IL6(TT) | 2.29 | 0.88 | 2.62 | ** |

Supplementary 12. P-values of the deletion and the log-likelihood ratio test, assessing the magnitude of the genetic effect retained in the model selection process. The non-significant p-values are not reported here. Statistical significance is represented by * for p < 0.05, ** for p < 0.01 and *** for p < 0.001.

| Phenotypic trait | | Loci | Nested model | | Model selected with the SNP | | Chisq | P-value | |
|---|---|---|---|---|---|---|---|---|---|
| | | | **model** | **log-lik** | **model** | **log-lik** | | | |
| **Immune cells** | **NKp46+ NK cells** | IL-6 | site x age (linear) + SMI | -160.0 | site x age (linear) + SMI + IL6 | -155.0 | 9.95 | 0.006 | ** |
| | | IL-10 | site x age (linear) + SMI | -160.0 | site x age (linear) + SMI + IL10 | -153.9 | 12.29 | 0.002 | ** |
| | | IL-13 | site x age (linear) + SMI | -159.4 | site x age (linear) + SMI + IL13 | -155.6 | 7.74 | 0.020 | * |
| | | TNF | site x age (linear) + SMI | -160.0 | site x age (linear) + SMI + TNF | -155.3 | 9.55 | 0.008 | ** |
| | **CD19+ B cells** | IL-10 | sex x age (linear) + site + SMI | -181.4 | sex x age (linear) + site + SMI + IL10 | -172.8 | 17.18 | 1.8E-04 | *** |
| | | TNF | sex x age (linear) + site + SMI | -181.4 | sex x age (linear) + site + SMI + TNF | -176.2 | 10.39 | 0.006 | ** |
| | | IL-13 | sex x age (linear) + site + SMI | -180.2 | sex x age (linear) + site + SMI + IL13 | -176.7 | 6.91 | 0.031 | * |
| | **CD11c+ DC** | IL-6 | site x age (linear) + SMI | -204.0 | site x age (linear) + SMI + IL6 | -199.2 | 9.55 | 0.008 | ** |
| | | IL-10 | site x age (linear) + SMI | -205.3 | site x age (linear) + SMI + IL10 | -192.8 | 24.95 | 1.8E-06 | *** |
| | | IL-13 | age (linear) x site + SMI | -204.2 | age (linear) x site + SMI + IL13 | -200.7 | 7.01 | 0.030 | * |
| | **CD4+ T cells** | H2-Aa | site + age (linear) + SMI | -60.7 | site x H2Aa + age (linear) + SMI | -50.4 | 20.72 | 0.004 | ** |
| | | H2-Eb | site + age (linear) + SMI | -65.5 | site x H2Eb + age (linear) + SMI | -55.1 | 20.87 | 0.002 | ** |
| | | IL-6 | site + age (linear) + SMI | -66.7 | site x IL6 + age (linear) + SMI | -54.8 | 23.92 | 2.2E-04 | *** |
| | | IL-10 | site + age (linear) + SMI | -66.1 | site x IL10 + age (linear) + SMI | -50.5 | 31.36 | 2.2E-05 | *** |
| | | TNF | site + age (linear) + SMI | -66.1 | age (linear) + site + SMI + TNF | -61.2 | 9.88 | 7.2E-03 | ** |
| | | IL-13 | site + age (linear) + SMI | -65.9 | age (linear) x IL13 + site + SMI | -58.7 | 14.33 | 0.002 | ** |
| | **CD8+ T cells** | H2-Aa | age (linear) + SMI | -109.1 | age (linear) + SMI + H2Aa | -106.1 | 6.01 | 0.049 | * |
| | | IL-6 | age (linear) + SMI | -112.2 | age (linear) + SMI + IL6 | -108.3 | 7.82 | 0.020 | * |
| | | IL-10 | age (linear) + SMI | -11.9 | age (linear) + SMI + IL10 | -108.3 | 7.23 | 0.027 | * |
| | **CD25+ FoxP3+ Tregs** | IL-6 | site x age (linear) + SMI | -107.4 | site x age (linear) + SMI + IL6 | -104.0 | 6.89 | 0.032 | * |
| | | IL-13 | site x age (linear) + SMI | -107.9 | site x age (linear) + SMI + IL13 | -103.0 | 9.89 | 0.007 | ** |
| | **F4/80+ macrophages** | IL-10 | age (linear) + SMI | -252.6 | age (linear) + SMI + IL10 | -246.7 | 11.83 | 0.003 | ** |
| | | IL-13 | age (linear) + SMI | -251.3 | age (linear) x IL13 + SMI | -246.1 | 10.38 | 0.016 | * |
| | **HGM** | IL-13 | site x age (linear) + sex + SMI | -302.0 | site x age (linear) + sex + SMI + IL13 | -299.0 | 7.48 | 0.024 | * |
| | **PMN** | IL-17a_U | site + age (classes) + SMI | -316.5 | site x IL17a_U + age (classes) + SMI | -307.2 | 18.71 | 0.017 | * |
| | | IL-17f | site + age (classes) + SMI | -326.5 | site x IL17f + age (classes) + SMI | -306.8 | 39.42 | 0.001 | *** |
| | **Neutrophils** | IL-6 | site x age (linear) + SMI | -142.0 | site x age (linear) + SMI + IL6 | -138.0 | 8.05 | 0.017 | * |
| | | IL-10 | site x age (linear) + SMI | -143.2 | site x age (linear) + SMI + IL10 | -134.8 | 16.81 | 2.2E-04 | *** |
| | **MDSC** | H2-Aa | site x age (linear) + SMI | -128.8 | site x age (linear) + SMI + H2Aa | -125.1 | 7.36 | 0.025 | * |
| | | IL-6 | site x age (linear) + SMI | -135.7 | site x age (linear) + SMI + IL6 | -129.9 | 11.63 | 0.003 | ** |
| | | IL-10 | site x age (linear) + SMI | -136.2 | site x age (linear) + SMI + IL10 | -129.3 | 13.88 | 9.7E-04 | *** |
| | | IL-17a_N | site x age (linear) + SMI | -136.2 | site x age (linear) + SMI + IL17a_N | -132.1 | 8.28 | 0.016 | * |
| **Antibodies** | **IgG** | H2-Ab | site + age (classes) | -162.4 | site + age (classes) + H2Ab | -159.2 | 6.52 | 0.038 | * |
| | **IgA** | IL-6 | site x sex + age (classes) | -199.6 | site x sex + age (classes) x IL6 | -192.4 | 14.49 | 0.024 | * |
| | | TLR-9 | site x sex + age (classes) | -200.9 | site x sex + age (classes) x TLR9 | -194.5 | 12.78 | 0.005 | ** |
| | **IgE** | IL-1a | site + sex + age (classes) | -244.0 | site x IL1a + sex + age (classes) | -221.7 | 44.51 | 1.0E-05 | *** |
| | | IL-1b_U | site + sex + age (classes) | -242.4 | site + sex + age (classes) + IL1b_U | -235.2 | 14.30 | 0.001 | *** |
| | | IL-6 | site + sex + age (classes) | -251.1 | site x IL6 + sex + age (classes) | -244.7 | 12.78 | 0.026 | * |
| | | IL-17a_U | site + sex + age (classes) | -238.4 | site x IL17a_U + sex + age (classes) | -221.6 | 33.68 | 2.0E-04 | *** |
| | | TNF | site + sex + age (classes) | -251.7 | site x TNF + sex + age (classes) | -243.8 | 15.86 | 0.015 | * |
| **Cytokines** | **PC1** | H2-Eb | site x sex | -453.7 | site x sex + H2Eb | -432.6 | 6.23 | 0.044 | * |
| | | IL-1b_U | site x sex | -435.7 | site x sex + IL1b_U | -432.2 | 7.02 | 0.030 | * |
| | | IL-6 | site x sex | -435.7 | site x sex + IL6 | -431.7 | 7.99 | 0.018 | * |
| | | IL-10 | site x sex | -435.7 | site x sex + IL10 | -423.0 | 25.28 | 4.9E-07 | *** |
| | | TNF | site x sex | -435.7 | site x sex + TNF | -428.2 | 15.05 | 0.001 | *** |
| | **PC2** | IL-10 | site x sex | -358.5 | site x sex + IL10 | -379.7 | 11.69 | 0.001 | *** |
| | | IL-17a_U | site x sex | -385.5 | site x sex + IL17a_U | -382.4 | 6.27 | 0.044 | * |
| **Macro-parasites** | Mites | H2-Aa | site + age (classes) + SMI | -674.8 | site + age (classes) + SMI + H2Aa | -678.6 | 7.62 | 0.022 | * |
| | | TLR-9 | site + age (classes) + SMI | -695.2 | site + age (classes) + SMI + TLR9 | -690.9 | 8.64 | 0.003 | ** |
| | Worms | H2-Ab | site + age (linear) + SMI | -375.4 | site x H2Ab + age (linear) + SMI | -360.4 | 29.92 | 0.005 | ** |
| | | IL-6 | site + age (linear) + SMI | -387.4 | site + age (linear) + SMI + IL6 | -381.8 | 11.17 | 0.004 | ** |
| **Micro-parasites** | Norovirus | H2-Eb | site + age (classes) | -291.7 | site + age (classes) x H2Eb | -284.2 | 15.08 | 0.020 | * |
| | | IL-10 | site + age (classes) | -295.3 | site + age (classes) x IL10 | -285.9 | 18.77 | 0.005 | ** |
| | Parvovirus | H2-Eb | site + age (classes) | -479.6 | site + age (classes) + H2Eb | -476.4 | 6.47 | 0.039 | * |
| | Minute virus | H2-Ab | site + age (classes) | -237.2 | site + age (classes) + H2Ab | -233.3 | 7.81 | 0.020 | * |
| | | IL-1a | site + age (classes) | -248.2 | site + age (classes) + IL1a | -245.0 | 6.34 | 0.042 | * |
| | Mouse hepatitis virus | H2-Aa | site + age (classes) | -260.4 | site + age (classes) + H2Aa | -254.0 | 12.78 | 0.002 | ** |
| | | IL-6 | site + age (classes) | -260.1 | site + age (classes) + IL6 | -256.1 | 7.90 | 0.019 | * |
| | Sendai virus | IL-6 | site + age (classes) | -295.1 | site + age (classes) + IL6 | -292.1 | 6.05 | 0.049 | * |
| | | TNF | site + age (classes) | -295.7 | site x TNF + age (classes) | -284.5 | 22.52 | 0.002 | ** |

| | | | | | | | | | |
|---|---|---|---|---|---|---|---|---|---|
| | | TLR-9 | site + age (classes) | -295.4 | site + age (classes) x TLR9 | -291.2 | 8.36 | 0.039 | * |
| | Coronavirus | IL17a_N | site + age (linear) | -129.0 | site + age (linear) x IL17a_N | -123.5 | 10.89 | 0.028 | * |
| | *M. pulmonis* | IL-6 | site + age (classes) | -283.6 | site + age (classes) + IL6 | -279.9 | 7.30 | 0.026 | * |
